# A framework for community curation of interspecies interactions literature

**DOI:** 10.1101/2022.12.15.520601

**Authors:** Alayne Cuzick, James Seager, Valerie Wood, Martin Urban, Kim Rutherford, Kim E. Hammond-Kosack

## Abstract

The quantity and complexity of data being generated and published in biology has increased substantially, but few methods exist for capturing knowledge about phenotypes derived from molecular interactions between diverse groups of species, in such a way that is amenable to data-driven biology and research. To improve access to this knowledge, we have constructed a framework for the curation of the scientific literature studying interspecies interactions, using data curated for the Pathogen-Host Interactions Database (PHI-base) as a case study. The framework provides a curation tool, phenotype ontology and controlled vocabularies to curate pathogen-host interaction data (at the level of the host, pathogen, strain, gene and genotype). The concept of a multispecies genotype, the ‘metagenotype’, is introduced to facilitate capturing changes in the pathogens’ disease-causing abilities, and host resistance or susceptibility observed by gene alterations. We report on this framework and describe PHI-Canto, a community curation tool for use by publication authors.

## Introduction

Recent technological advancements across the biological sciences have resulted in an increasing volume of peer-reviewed publications reporting experimental data and conclusions. To increase the value of this highly fragmented knowledge, biocurators manually extract the data from publications and represent it in a standardized and interconnected way in accordance with the FAIR (Findable, Accessible, Interoperable and Reusable) Data Principles (International Society for Biocuration, 2018; Wilkinson et al., 2016). The curated data is then made available in online databases, either organism- or clade-specific (e.g., model organism databases) or those supporting multiple kingdoms of life (e.g., PHI-base, Alliance of Genomes Resources (Alliance of Genome Resources Consortium, 2020; Urban et al., 2021)). Due to the complexity of the biology, manual biocuration is currently the only way to reliably represent information about function and phenotype in databases and knowledge bases (Wood, Sternberg, & Lipshitz, 2022). The development of curation tools with clear workflows supporting the use of biological ontologies and controlled vocabularies has standardized curation efforts, reduced ambiguity in annotation and improved the maintenance of the curated corpus as biological knowledge evolves (International Society for Biocuration, 2018).

The pathogen-host interaction research communities are an example of a domain of the biological sciences exhibiting a literature deluge (Figure 1). The Pathogen-Host Interactions Database, PHI-base (phi-base.org), is an open access FAIR biological database containing data on bacterial, fungal and protist genes proven to affect the outcome of pathogen-host interactions (Rodriguez-Iglesias et al., 2016; Urban et al., 2021). Viruses are not included in PHI-base. Since 2005, PHI-base has manually curated phenotype data associated with underlying genome-level changes from peer-reviewed pathogen-host interaction literature. Information is also provided on the target sites of some anti-infective chemistries (Urban et al., 2020). This type of data is increasingly relevant, as infectious microbes continually threaten global food security, human health across the life course, farmed animal health and wellbeing, tree health and ecosystem resilience (Brown et al., 2012; Fisher, Hawkins, Sanglard, & Gurr, 2018; Fisher et al., 2012; Smith, Machalaba, Seifman, Feferholtz, & Karesh, 2019). Rising resistance to antimicrobial compounds, increased globalization, and climate change indicate that infectious microbes will present ever greater economic and societal threats (Bebber, Ramotowski, & Gurr, 2013; Chaloner, Gurr, & Bebber, 2021; Cook et al., 2021). In order to curate relevant publications into PHI-base (version 4) professional curators have, since 2011, entered 81 different data types into a text file (Urban et al., 2017). However, increasing publication numbers and data complexity required more robust curation procedures.

**Figure 1.**
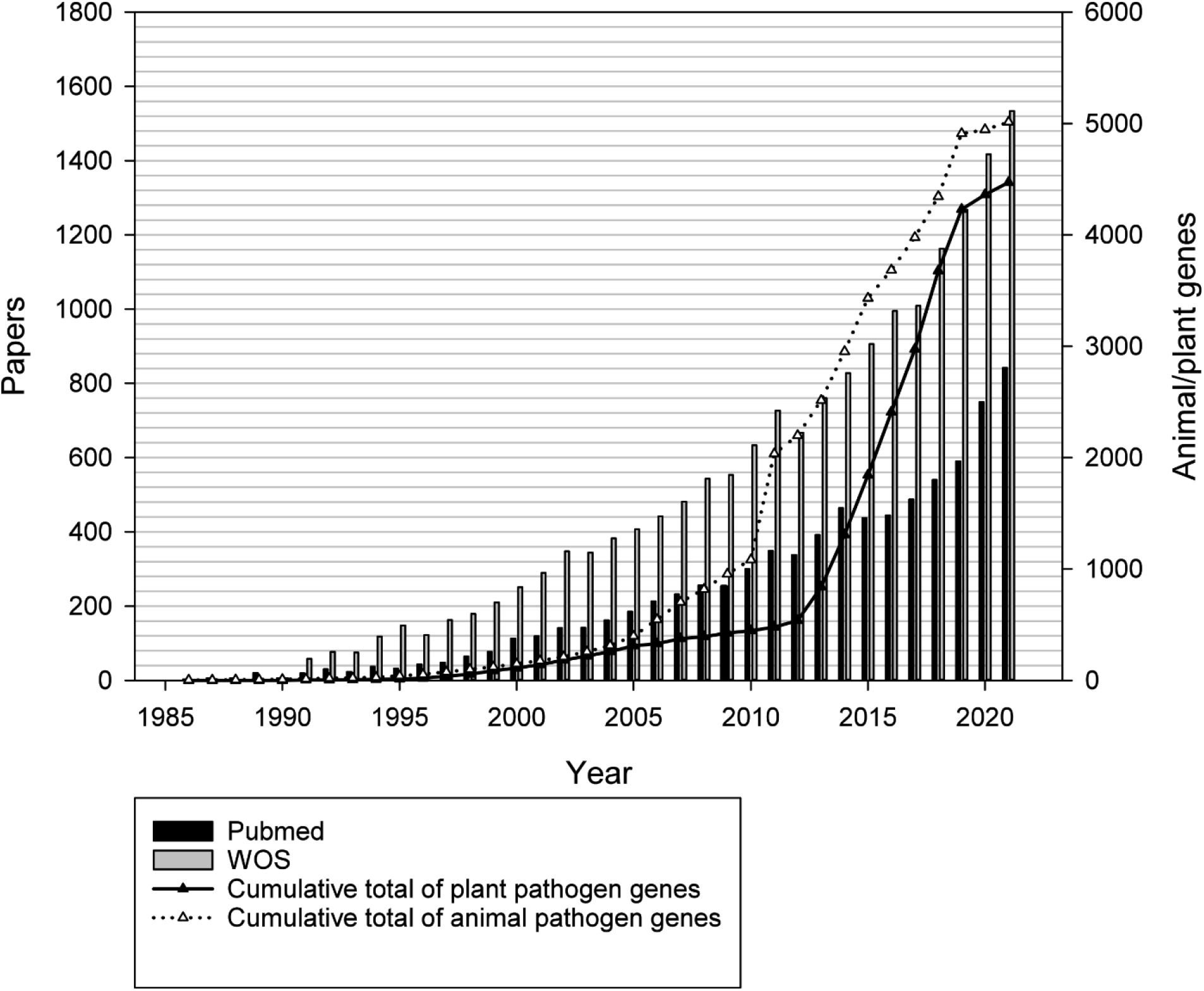
Increase of molecular host-pathogen interaction publications and gene phenotype information during the last 35 years curated in PHI-base. Grey bars show the number of publications in the Web of Science Core Collection database retrieved with search term “(fung* or yeast) and (gene or factor) and (pathogenicity or virulen* or avirulence gene*)”. Black vertical bars show the number of articles retrieved from PubMed (searching on title and abstract). Black and white triangles show the number of curated animal and plant pathogen genes, respectively.

We were unable to locate any curation frameworks or tools capable of capturing the interspecies interactions required for PHI-base. PomBase, the fission yeast (*Schizosaccharomyces pombe*) database developed Canto, a web-based tool supporting curation by both professional biocurators and publication authors (Rutherford, Harris, Lock, Oliver, & Wood, 2014). While Canto could support annotation for multiple species, it could not annotate interactions between species. Therefore, we extended and customized Canto to support interspecies interactions, creating a new tool: PHI-Canto (the Pathogen-Host Interaction Community Annotation Tool). Likewise, there were no existing biomedical ontologies that could accurately describe pathogen-host interaction phenotypes at the depth and breadth required for PHI-base. Infectious disease formation depends on a series of complex and dynamic interactions between pathogenic species and their potential hosts, and also requires the correct biotic and/or abiotic environmental conditions (Scholthof, 2007), as illustrated by the concept of the ‘disease triangle’ (Figure 2). All these interrelated factors must be recorded in order to sufficiently describe a pathogen-host interaction.

**Figure 2.**
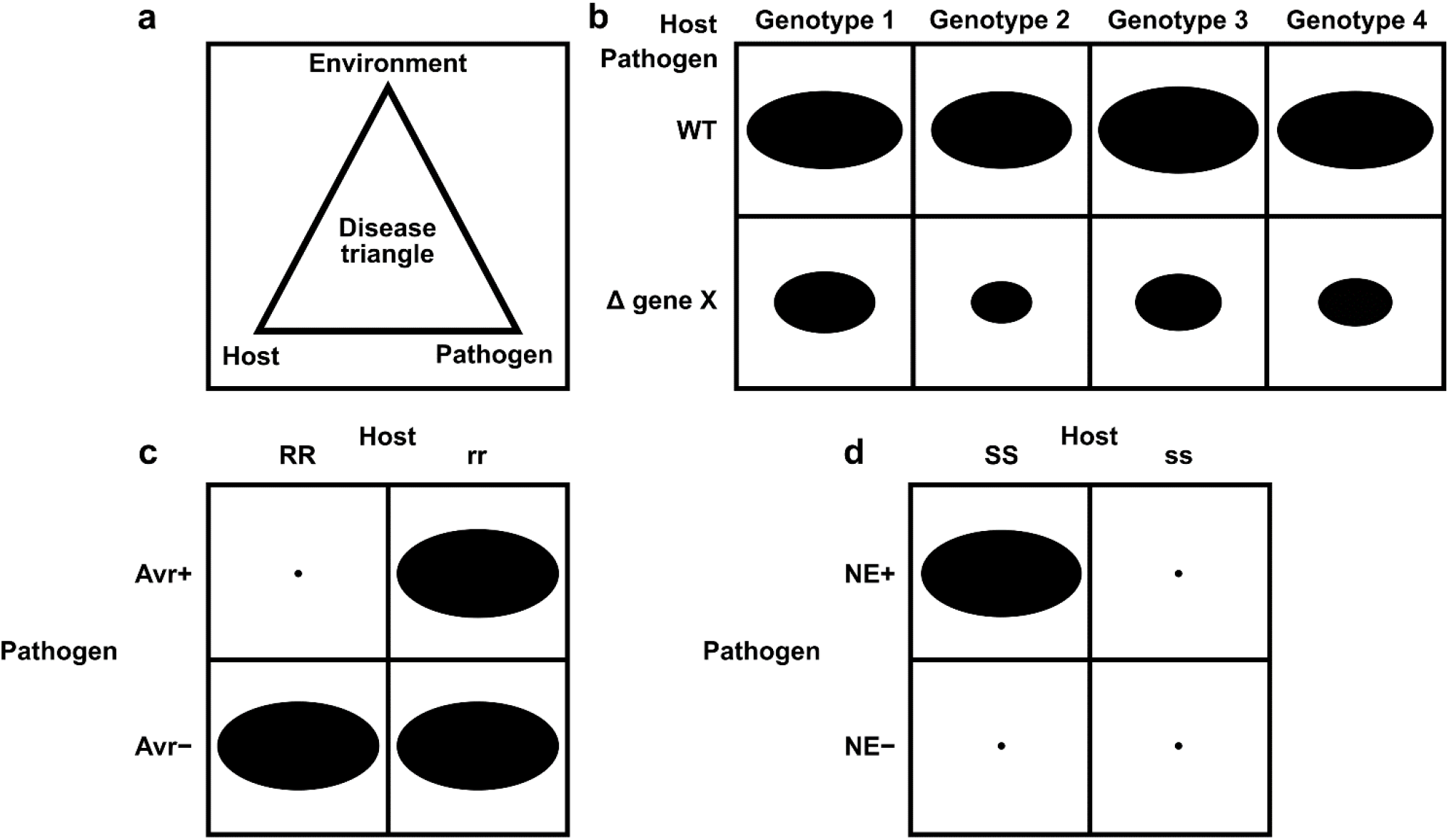
Schematic representation of pathogen-host interactions. **(a)** the disease triangle illustrates the requirement for the correct abiotic and biotic environmental conditions to ensure disease when an adapted pathogen encounters a suitable host; **(b)** a non gene-for-gene genetic relationship where compatible interactions result in disease on all host genotypes (depicted as genotypes 1–4), but the extent of disease formation is influenced to a greater or lesser extent by the presence or absence of a single pathogen virulence gene product X. In host genotypes 1 and 3, the pathogen gene product X is the least required for disease formation. The size of each black oval in each of the eight genetic interactions indicates the severity of the disease phenotype observed, with a larger oval indicating greater severity: **(c)** a gene-for-gene genetic relationship. In this genetic system, considerable specificity is observed, which is based on the direct or indirect interaction of a pathogen avirulence (*Avr*) effector gene product with a host resistance (*R*) gene product to determine specific recognition (an incompatible interaction), which is typically observed in biotrophic interactions ((Jones & Dangl, 2006)). In one scenario, the product of the *Avr* effector gene binds to the product of the *R* gene (a receptor) to activate host resistance mechanisms. In another scenario, the product of the *Avr* effector gene binds to an essential host target which is guarded by the product of the *R* gene (a receptor). Once *Avr* effector binding is detected, host resistance mechanisms are activated. The absence of the *Avr* effector product or the absence of the *R* gene product leads to susceptibility (a compatible interaction). The small black dot indicates no disease formation, and the large black oval indicates full disease formation, and **(d)** an inverse gene-for-gene genetic relationship. Again, considerable specificity is observed based on the interaction of a pathogen necrotrophic effector (NE) with a host susceptibility (*S*) target to determine specific recognition. The product of the pathogen NE gene binds to the product of the *S* gene (a receptor) to activate host susceptibility mechanisms.

In this study three key issues were addressed in order to develop the curation framework for interspecies interactions: firstly, to support the classification of genes as ‘pathogen’ or ‘host’, and enable the variations of the same gene in different strains to be captured; secondly, formulating the concept of a ‘metagenotype’ to represent the interaction between specific strains of both a pathogen and a host within a multispecies genotype; and thirdly, developing supporting ontologies and controlled vocabularies, including the generic Pathogen-Host Interaction Phenotype Ontology (PHIPO), to annotate phenotypes connected to genotypes at the level of a single species (pathogen or host) and multiple species (pathogen-host interaction phenotypes).

## Results

### Enabling multispecies curation with UniProtKB accessions

In any curation context, stable identifiers are required for annotated entities. The UniProt Knowledgebase (UniProtKB) (UniProt Consortium, 2021) is universally recognized, provides broad taxonomic protein coverage, and manually curates standard nomenclature across protein families. Protein sequences are both manually and computationally annotated in UniProtKB, providing a wealth of data on catalytic activities, protein structures and protein-protein interactions, Gene Ontology (GO) annotations and links to PHI-base phenotypes (Ashburner et al., 2000; Gene Ontology Consortium, 2021; Urban et al., 2021). To improve interoperability with other resources, we used UniProtKB accession numbers for retrieving protein entities, gene names and species information for display in PHI-Canto. PHI-Canto accesses the UniProtKB API to automatically retrieve the entities and their associated data.

### Developing the metagenotype to capture interspecies interactions

To enable annotation of interspecies interactions, we developed the concept of a ‘metagenotype’, that represents the combination of a pathogen genotype and a host genotype (Figure 3). A metagenotype is created after the individual genotypes from both species are created. Each metagenotype can be annotated with pathogen-host interaction phenotypes to capture changes in pathogenicity (caused by alterations to the pathogen) and changes in virulence (caused by alterations to the host and/or the pathogen). Metagenotypes must always include at least one named pathogen gene with a genotype of interest, but a metagenotype can be composed from a pathogen genotype and a host species (and strain) if no specific host gene is referenced in an experiment.

**Figure 3.**
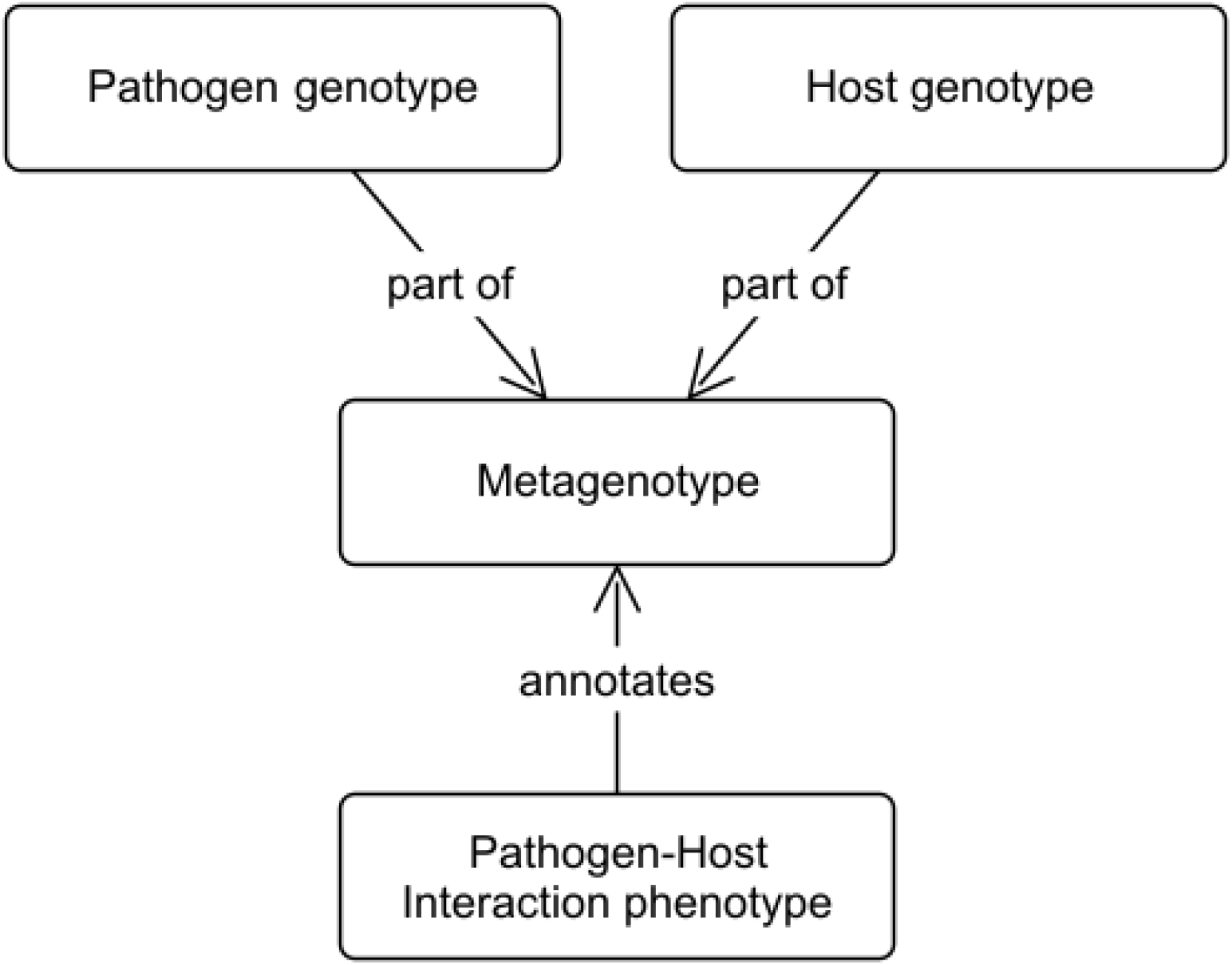
Conceptual model showing the relationship between metagenotypes, genotypes and annotations. The curator selects a pathogen genotype and a host genotype to combine into a metagenotype. The metagenotype can be annotated with pathogen-host interaction phenotypes from PHIPO (the Pathogen-Host Interaction Phenotype Ontology).

### Annotation types and annotation extensions in PHI-Canto

In PHI-Canto, ‘annotation’ is the task of relating a specific piece of knowledge to a biological feature. To curate a wide variety of experiment types, three groupings of annotation types are available in PHI-Canto, covering ‘metagenotype’, ‘genotype’ (of a single species) and ‘gene’ annotation types (Table 1). To capture additional biologically relevant information associated with an annotation, curators use annotation extensions (Huntley et al., 2014) to extend the primary annotation. For the purpose of Canto and PHI-Canto, the meaning of ‘annotation extension’ was broadened to capture additional properties related to the annotation, such as the metagenotype used as an experimental control. The additional properties that may be related to an annotation are simply referred to as ‘annotation extensions’ (AEs) in this manuscript (Table 1, Supplementary file 1 and Supplementary file 2). Descriptions of the new AEs for PHI-Canto and the core collection of AEs from Canto are available in the PHI-Canto user documentation (see the Code availability section).

**Table 1.**
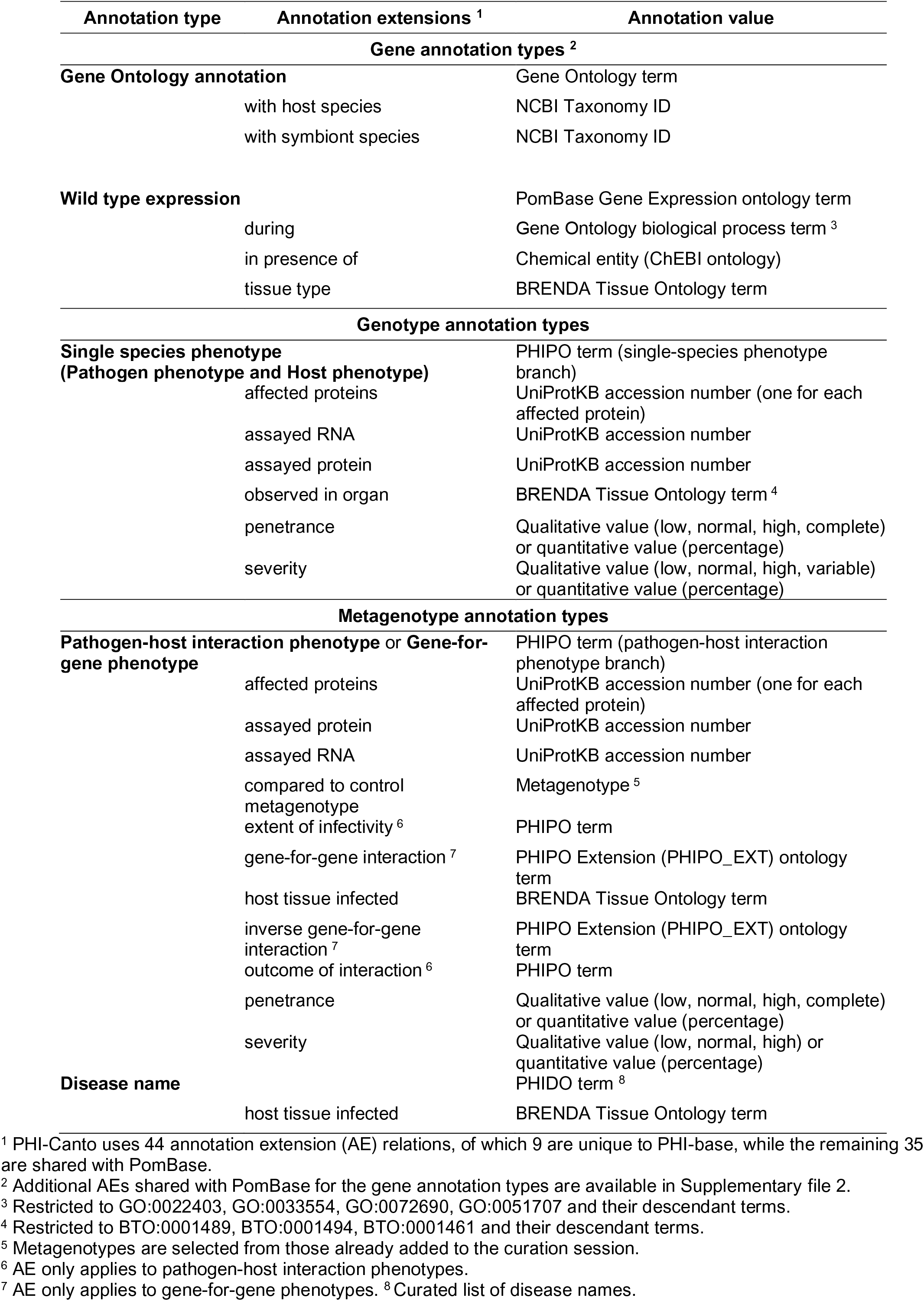
Annotation types and selected annotation extensions used in PHI-Canto.

Metagenotypes can be annotated with terms from an ontology or controlled vocabulary following either the ‘pathogen-host interaction phenotype’, ‘gene-for-gene phenotype’ or ‘disease name’ annotation types (Table 1). Phenotype annotations can be supported by AEs providing additional qualifying information required to fully interpret the experiment, such as the infected tissue of the host.

Phenotypes can also be curated for single species experiments, involving either the pathogen or host, following the ‘single species phenotype’ annotation workflow (Table 1). Single species phenotype annotations have a selection of AEs available, including the protein assayed in the experiment and the severity of the observed phenotype (see example from PMID:22314539 in Appendix 1).

PHI-Canto also supports the annotation of gene and gene product attributes to represent the evolved functional role of a gene product, described here as the ‘gene annotation’ workflow (Table 1). The Gene Ontology is used for annotation of a gene product’s molecular functions, biological processes and cellular components, while PSI-MOD is used for the annotation of protein modifications (Montecchi-Palazzi et al., 2008), and BioGRID experiment types are used to capture genetic and physical interactions (Oughtred et al., 2021). GO annotations are submitted to the EBI GO Annotation Database (GOA), from where they are propagated to the main GO database (Gene Ontology Consortium, 2021; Huntley et al., 2015).

### Curation of interspecies interaction publications

Ten publications covering a wide range of typical plant, human, and animal pathogen-host interactions were selected for trial curation in PHI-Canto (Table 2). These publications included experiments with early acting pathogen virulence proteins, the first host targets of pathogen effectors, and resistance to antifungal chemistries. These publications guided development of the ontology and controlled vocabulary terms required for PHI-Canto, as well as the curation methods required for different experiments. Major curation problems and their solutions are summarized in Table 3, and example annotations are described below and provided in Appendix 1 and Appendix 2.

**Table 2.** Publications selected for trial curation using PHI-Canto.

| Subject of publication | PMID | Publication title | Genotype <sup>10</sup> annotated with | Metagenotype <sup>11</sup> annotated with |
| --- | --- | --- | --- | --- |
| **Bacteria–human interaction** | 28715477 <sup>1</sup> | The RhlR quorum-sensing receptor controls *Pseudomonas aeruginosa* pathogenesis and biofilm development independently of its canonical homoserine lactone autoinducer. | Pathogen phenotype | unaffected pathogenicity<br>altered pathogenicity or virulence |
| **Fungal-human interaction/novel antifungal target** | 28720735 <sup>2</sup> | A nonredundant phosphopantetheinyl transferase, PptA, is a novel antifungal target that directs secondary metabolite, siderophore, and lysine biosynthesis in *Aspergillus fumigatus* and is critical for pathogenicity. | Pathogen phenotype | unaffected pathogenicity<br>altered pathogenicity or virulence |
| **Secondary metabolite clusters required for pathogen virulence** | 30459352 <sup>2</sup> | Phosphopantetheinyl transferase (Ppt)-mediated biosynthesis of lysine, but not siderophores or DHN melanin, is required for virulence of *Zymoseptoria tritici* on wheat. | Pathogen phenotype | unaffected pathogenicity<br>altered pathogenicity or virulence |
| **Early acting virulence proteins** | 29020037 <sup>2,3</sup> | A conserved fungal glycosyltransferase facilitates pathogenesis of plants by enabling hyphal growth on solid surfaces. | Pathogen phenotype | altered pathogenicity or virulence |
| **Mutualism interaction** | 16517760 <sup>4</sup> | Reactive oxygen species play a role in regulating a fungus-perennial ryegrass mutualistic interaction | Pathogen phenotype | mutualism |
| **First host targets of pathogen effectors** | 31804478 <sup>2,5</sup> | An effector protein of the wheat stripe rust fungus targets chloroplasts and suppresses chloroplast function. | N/A | altered pathogenicity or virulence<br>a pathogen effector |
| **Receptor decoys** | 30220500 <sup>5</sup> | Suppression of plant immunity by fungal chitinase-like effectors. | Pathogen phenotype | a pathogen effector |
| **R-Avr interactions** | 20601497 <sup>6,7</sup> | Activation of an Arabidopsis resistance protein is specified by the *in planta* association of its leucine-rich repeat domain with the cognate oomycete effector. | Host phenotype | a pathogen effector<br>a gene-for-gene interaction |
| **Fungal toxins required for virulence on plants** | 22241993 <sup>8</sup> | The cysteine rich necrotrophic effector SnTox1 produced by *Stagonospora nodorum* triggers susceptibility of wheat lines harboring Snn1. | N/A | a pathogen effector<br>a gene-for-gene interaction (inverse) |
| **Resistance to antifungal chemistries** | 22314539 <sup>9</sup> | The T788G mutation in the cyp51C gene confers voriconazole resistance in *Aspergillus flavus* causing aspergillosis. | Pathogen phenotype<br>Pathogen chemistry phenotype |  |

<sup>1</sup> Example of curating 'unaffected pathogenicity' available in Appendix 1.<sup>2</sup> Example of curating 'altered pathogenicity or virulence' available in Appendix 1 and Appendix 2.<sup>3</sup> Example of 'in vitro pathogen phenotype' available in Appendix 1.<sup>4</sup> Example of curating 'mutualism' available in Appendix 1.<sup>5</sup> Example of curating 'a pathogen effector' available in Appendix 1.<sup>6</sup> Example of curating 'a gene-for-gene interaction' available in Appendix 1.<sup>7</sup> Example of 'in vitro host phenotype' available in Appendix 1.<sup>8</sup> Example of curating 'an inverse gene-for-gene interaction' available in Appendix 1.<sup>9</sup> Example of 'in vitro pathogen chemistry phenotype' available in Appendix 1.<sup>10</sup> Single species genotypes could be annotated with either a pathogen phenotype, a pathogen chemistry phenotype, or a host phenotype. Genotypes are annotated with *in vitro* or *in vivo* phenotypes from PHIPO, using either the Pathogen phenotype or Host phenotype annotation type workflow.<sup>11</sup> Metagenotype comprises of a pathogen and a host genotype in combination. Phenotypes from PHIPO can be annotated to metagenotypes using either the 'Pathogen-Host Interaction Phenotype' or 'Gene-for-Gene Phenotype' annotation type workflow.

**Table 3.** Issues encountered whilst curating ten example publications with PHI-Canto.

| Curated feature | Problem description | Solution | Context in PHI-Canto | Example |
| --- | --- | --- | --- | --- |
| Species strain | UniProtKB sequence information is commonly from a reference genome strain. This sequence may differ from the experimental strain curated in PHI-Canto. | Develop a selectable list of strains for curators to assign to the genotype (and metagenotype). | Strain selected after UniProtKB entry on gene entry page. Strain used within genotype creation. | URL<sup>1</sup><br>All phenotype annotation examples in Appendix 1 contain a 'strain name' within the genotype / metagenotype. |
| Delivery mechanism | Pathogen-host interaction experiments use a wide array of mechanisms to deliver the treatment of choice (to cells, tissues, and host and non-host species) which are required for experimental interpretation. | Develop terms prefixed with 'delivery mechanism' in the Pathogen-Host Interaction Experimental Conditions Ontology (PHI-ECO). | Selection of experimental conditions whilst making a phenotype annotation to a metagenotype. | URL<sup>2</sup><br>Examples in Appendix 1 PMID:20601497, PMID:31804478 and PMID:22241993. |
| Physical interaction | Physical interactions (i.e., protein-protein interactions) could only be annotated between proteins of the same species, so it was not possible to annotate interactions between a pathogen effector and its first host target. | Adapt the 'Physical Interaction' annotation type to store gene and species information from two organisms (instead of one). | Physical Interaction annotation type. | URL<sup>3</sup> |
| Pathogen effector | There was no available ontology term to describe a 'class' pathogen effector (a 'transferred entity from pathogen to host'), because effectors have heterogeneous functions (specific enzyme inhibitors, modulating host immune responses, and targeting host gene-silencing mechanisms). Effector is not a phenotype, and so did not fit into the Pathogen-Host Interaction Phenotype Ontology (PHIPO). | Develop new Gene Ontology (GO) biological process terms (and children), to group 'effector-mediated' processes. | GO Biological Process annotation on a pathogen gene. | URL<sup>4</sup><br>Example in Appendix 1 PMID:31804478. |
| Wild type control phenotypes | Natural sequence variation between strains of both pathogen and host organisms can alter the phenotypic outcome within an interaction. The wild type metagenotype phenotype needs to be curated so that the phenotype of an altered metagenotype is informative. | Allow creation of metagenotypes containing wild type genes. Develop a new annotation extension (AE) property 'compared to control', used in annotation of altered metagenotypes. | Annotation of phenotypes and AEs to metagenotypes (using the 'PHI phenotype' or 'Gene for Gene phenotype' annotation type). | URL<sup>5</sup><br>Examples in Appendix 1 PMID:28715477, PMID:16517760, PMID:29020037, PMID:20601497, PMID:22241993. |
| Chemistry | How to record chemicals for resistance or sensitivity phenotypes. | Follow PomBase model to pre-compose PHIPO terms to include chemical names from the ChEBI ontology. | Annotation of phenotypes to single species genotypes. | URL<sup>4</sup><br>Example in Appendix 1 PMID:22314539. |
| Gene for gene interactions | Complex gene-for-gene interactions within plant pathogen-host interactions required additional detail to describe the function of the pathogen and host genes within the metagenotype (including the specified strains). | Develop the additional metagenotype curation type 'Gene for Gene Phenotype'. Develop two new AEs, 'gene_for_gene interaction' and 'inverse gene_for_gene interaction', using PHIPO_EXT terms describing three components of the interaction. | Annotation of phenotypes and AEs to metagenotypes using the 'Gene for Gene Phenotype' annotation type. | URL<sup>4</sup><br>Examples in Appendix 1 PMID:20601497 and PMID:22241993. |
| Nine high-level legacy terms (from PHI-base 4) | PHI-base should incorporate legacy data from PHI-base 4 into new PHI-base 5 gene-centric pages. | Maintain the nine high level terms as 'tags' within the new PHI-base 5 user interface. Develop mapping methods to enable this. | Three locations described in Supplementary file 3. | Urban et al., 2015 NAR (PMID:25414340). |

\*Namely, i) the compatibility of the interaction ii) the functional status of the pathogen gene and iii) the functional status of the host gene. URL<sup>1</sup> https://canto.phi-base.org/docs/getting_started#adding_strains, URL<sup>2</sup> https://canto.phi-base.org/docs/phipo_annotation#experimental_conditions, URL<sup>3</sup> https://canto.phi-base.org/docs/physical_interaction_annotation, URL<sup>4</sup> https://canto.phi-base.org/docs/phipo_annotation#pathogen_host_interaction_phenotypes, URL<sup>5</sup> https://canto.phi-base.org/docs/genotypes#metagenotype_management

#### Curating an experiment with a metagenotype

A large proportion of the curation in PHI-Canto requires the use of metagenotypes: one of the simpler cases involves early acting virulence proteins, where a genetically modified pathogen is inoculated onto a host (without a specified host gene). A metagenotype is created and annotated with a phenotype term. These experiments are curated following the ‘pathogen-host interaction phenotype’ workflow, including any relevant AEs (Table 1). This two-step curation process is illustrated by PMID:29020037 curation (Table 2, Appendix 1 and Appendix 2). The GT2 gene is deleted from the fungal plant pathogen *Zymoseptoria septoria* and inoculated onto wheat plants; the observed phenotype ‘absence of pathogen-associated host lesions’ (PHIPO:0000481) is annotated to the metagenotype; and the AE for ‘infective ability’ is annotated with ‘loss of pathogenicity’.

#### Curating pathogen effector experiments

A pathogen effector is defined as an entity transferred between the pathogen and the host that is known or suspected to be responsible for either activating or suppressing a host process commonly involved in defense (Houterman et al., 2009; Jones & Dangl, 2006) (Figure 2). To curate an effector experiment, first a metagenotype is created, then annotated with a phenotype term. To indicate that the pathogen gene functions as an effector, it is necessary to also make a concurrent ‘gene annotation’ (Table 1) with the GO biological process term ‘effector-mediated modulation of host process’ (GO:0140418) or an appropriate descendant term. This GO term has been created (with descendants) in collaboration with the Gene Ontology Consortium (GOC) and is used to identify pathogen effectors in PHI-base (version 5) (Supplementary file 3). Molecular functions of the pathogen gene can be curated with a GO molecular function term, if reported in the literature, and connected to the GO biological process term. An example of curation of a pathogen effector experiment is illustrated using PMID:31804478 (Table 2 and Appendix 1) where the pathogen effector Pst_12806 from *Puccinia striiformis* suppresses pattern-triggered immunity in a tobacco leaf model. Here, the metagenotype is curated with the phenotype ‘decreased level of host defense-induced callose deposition’ (PHIPO:0001015) and the effector is annotated with ‘effector-mediated suppression of host pattern-triggered immunity’ (GO:0052034). A further experiment demonstrated that the pathogen effector protein was able to bind to the natural host (wheat) protein PetC and inhibit the enzyme activity of PetC, resulting in a GO molecular function annotation ‘enzyme inhibitor activity’ (GO:0004857) on Pst_12806, with PetC captured as the target protein in an AE (see Appendix 1).

#### Curating experiments with a gene-for-gene relationship

For a gene-for-gene pathogen-host interaction type (when a known genetic interaction is conferred by a specific pathogen avirulence gene product and its cognate host resistance gene product) (Figure 2c, d, further described in the figure legend) (Flor, 1956; Jones & Dangl, 2006; Kanyuka, Igna, Solomon, & Oliver, 2022) the ‘gene-for-gene phenotype’ metagenotype workflow is followed. The metagenotypes and phenotype annotations are made in the same way as the standard ‘pathogen-host interaction phenotype’ workflow, but with different supporting data. A new AE was developed to indicate the following three components of the interaction: i) the compatibility of the interaction, ii) the functional status of the pathogen gene, and iii) the functional status of the host gene. An example of an annotation for a biotrophic pathogen gene-for-gene interaction has been illustrated with PMID:20601497 (Table 2 and Appendix 1). Inverse gene-for-gene relationships occur with necrotrophic pathogens, where the pathogen necrotrophic effector interacts with a gene product from the corresponding host susceptibility locus and activates a host response that benefits the pathogen (a compatible interaction). If the necrotrophic effector cannot interact with the host target, then no disease occurs (an incompatible interaction) (Breen, Williams, Winterberg, Kobe, & Solomon, 2016). An example of an inverse gene-for-gene interaction using the appropriate AEs is illustrated with PMID:22241993 (Table 2 and Appendix 1).

#### Curating an experiment with a single species genotype in the presence or absence of a chemical

Single species genotypes (pathogen or host) can also be annotated with phenotypes following the ‘single species phenotype annotation type’ workflow (Table 1). This is illustrated using PMID:22314539 in Table 2 (and Appendix 1) with an example of an *in vitro* pathogen chemistry phenotype, where a single nucleotide mutation in the *Aspergillus flavus* CYP51c gene confers ‘resistance to voriconazole’ (PHIPO:0000590), an antifungal agent.

#### Supporting curation of legacy information

PHI-Canto’s curation workflows maintain support for nine high-level terms that describe phenotypic outcomes essential for taxonomically diverse interspecies comparisons, which were the primary annotation method used in previous versions of PHI-base (Urban et al., 2015) and which are displayed in the Ensembl Genomes browser (Yates et al., 2021). For example, the ‘infective ability’ AE can be used to annotate the following subset of high-level terms: ‘loss of pathogenicity’, ‘unaffected pathogenicity’, ‘reduced virulence’, ‘increased virulence’ and ‘loss of mutualism’ (formerly ‘enhanced antagonism’). The mapping between the nine high-level terms and the PHI-Canto curation process is further described in Supplementary file 3.

#### Resolving additional problems with curating complex pathogen-host interactions

Table 3 shows a selection of the problems encountered during the development of PHI-Canto and the solutions we identified. For example, recording the delivery mechanism used within the pathogen-host interaction experiment. New experimental condition terms were developed with a prefix of ‘delivery mechanism’, for example, ‘delivery mechanism: agrobacterium’, ‘delivery mechanism: heterologous organism’, and ‘delivery mechanism: pathogen inoculation’. Another issue encountered was how to record a ‘physical interaction’ between proteins of different species, especially for the curation of pathogen effector first host targets. This was resolved by adapting the existing Canto module for curating physical interactions to support two different species.

### Development of the Pathogen-Host Interaction Phenotype Ontology and additional data lists

To support the annotation of phenotypes in PHI-Canto, the Pathogen-Host Interaction Phenotype Ontology (PHIPO) was developed. PHIPO is a species-neutral phenotype ontology that describes a broad range of pathogen-host interaction phenotypes. PHIPO’s terms were developed following a pre-compositional approach, where the term names and semantics are composed from existing terms from other ontologies, in order to make the curation process easier. For example, the curator annotates ‘resistance to penicillin’ (PHIPO:0000692) instead of ‘increased resistance to chemical’ (PHIPO:0000022) and ‘penicillin’ (CHEBI:17334) separately. Terms in PHIPO have logical definitions that follow design patterns from the uPheno ontology (Shefchek et al., 2020), and mapping PHIPO terms to uPheno patterns is an ongoing effort. These logical definitions provide relations between phenotypes in PHIPO and terms in other ontologies, such as PATO, GO, and ChEBI. PHIPO is available in OWL and OBO formats from the OBO Foundry (Jackson et al., 2021).

PHI-Canto uses additional controlled vocabularies derived from data in PHI-base. To enable PHI-Canto to distinguish between pathogen and host organisms, we extracted a list of > 250 pathogen and > 200 host species from PHI-base (Supplementary file 4). A curated list of strain names and their synonyms for the species currently curated in PHI-base was also developed for use in PHI-Canto (Supplementary file 4 and 5). PHI-base uses ‘strain’ as a grouping term for natural pathogen isolates, host cultivars and landraces, all of which are included in the curated list. The curation of pathogen strain designations was motivated by the NCBI Taxonomy’s decision to discontinue the assignment of strain-level taxonomic identifiers (Federhen et al., 2014) and a lack of standardized nomenclature for natural isolates of non-model species. New strain designations can be requested by curators and are reviewed by an expert prior to inclusion to ensure that each describes a novel strain designation rather than a new synonym for an existing strain.

Annotations in PHI-Canto include experimental evidence, which is specified by a term from a subset of the Evidence & Conclusion Ontology (ECO) (Giglio et al., 2019). Experimental evidence codes specific to pathogen-host interaction experiments have been developed and submitted to ECO. Phenotype annotations also include experimental conditions that are relevant to the experiment being curated, which are sourced from the PHI-base Experimental Conditions Ontology (PHI-ECO).

PHI-Canto includes a ‘disease name’ annotation type (Table 1) for annotating the name of the disease caused by an interaction between the pathogen and host specified in a wild type metagenotype (this annotation type is described in the PHI-Canto user documentation). Diseases are specified by a controlled vocabulary of disease names (called PHIDO), which was derived from disease names curated in previous versions of PHI-base.

### Summary of the PHI-Canto curation process

The PHI-Canto curation process is outlined in Figure 4, Figure 4 – figure supplement 1, the PHI-Canto user documentation and a detailed worked example is provided in Appendix 2. Each curation session is associated with one publication (using its PubMed identifier). One or more curators can collaborate on curating the same publication. An instructional email is sent to curators when they begin a new curation session, and PHI-base provides further guidelines on what information is needed in order to curate a publication in PHI-Canto (Figure 4 – figure supplement 2) and how to identify UniProtKB accession numbers from reference proteomes (Figure 4 – figure supplement 3).

**Figure 4.**
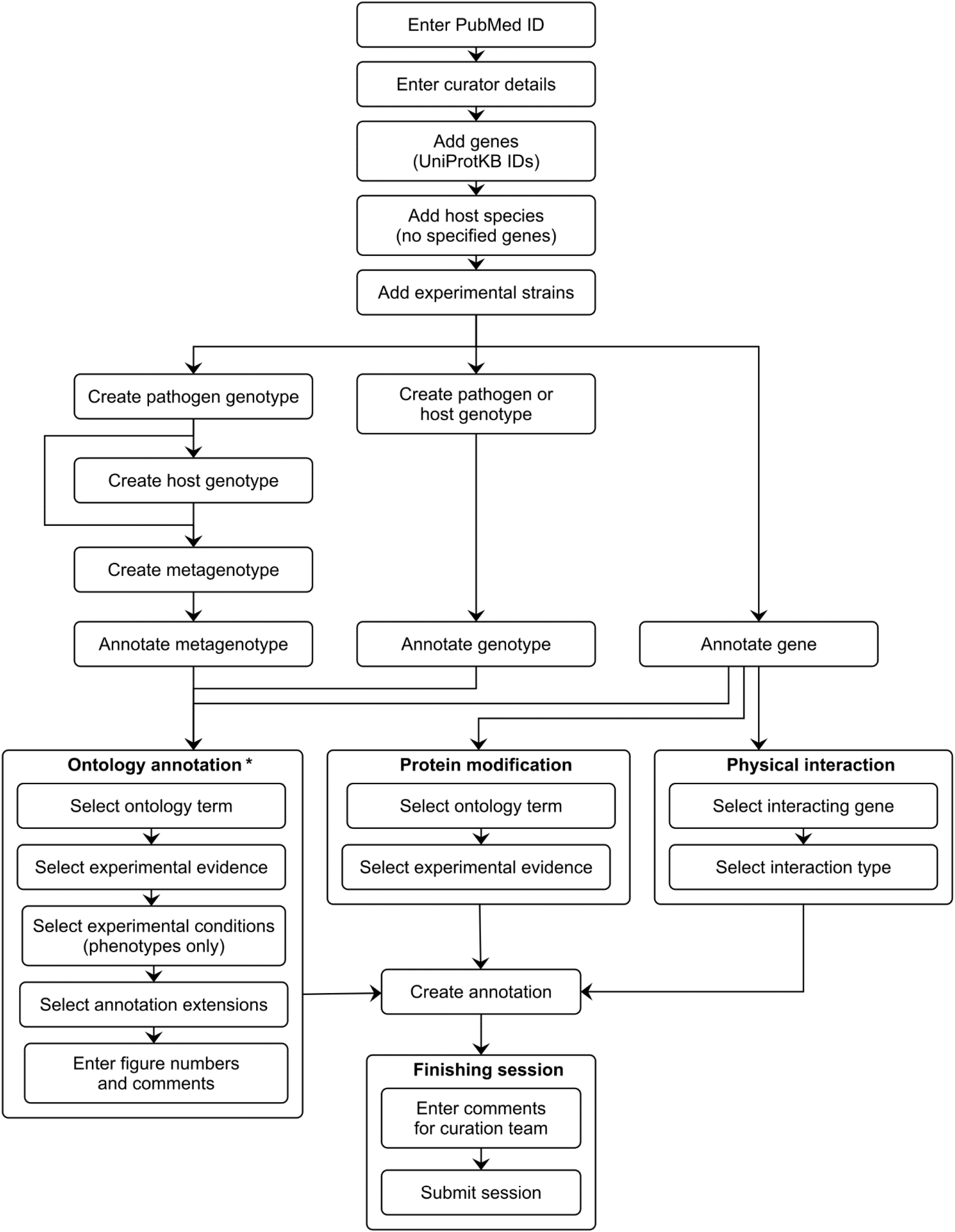
PHI-Canto curation workflow diagram. This diagram shows the curation workflow from the start of a curation session to its submission. The PubMed ID of the publication to be curated is entered and the title is automatically retrieved. The curator enters their name, email address and ORCID iD. On the species and genes page, the experimental pathogen and host genes are entered using UniProtKB accession numbers, and for experiments where a mutant pathogen genotype is assayed on a wild type host with no specified genes, there is the option to select the host species from an autocomplete menu. Information on the specific experimental strains used for each species is entered. After entering this initial information, the curator follows one of three distinct workflows depending on the biological feature the user wants to annotate (metagenotype, genotype or gene annotation type). Except for genes, biological features are created by composing less complex features: genotypes from alleles (generated in the pathogen or host genotype management pages), and metagenotypes from genotypes (generated in the metagenotype management page). Biological features are annotated with terms from a controlled vocabulary (usually an ontology), plus additional information that varies based on the annotation type. The curator has the option to generate further annotations after creating one, but this iterative process is not represented in the diagram for the sake of brevity. After all annotations have been made, the session is submitted to PHI-base. * Note that the ‘Ontology annotation’ group covers multiple annotation types, all of which annotate biological features with terms from an ontology or controlled vocabulary. These annotation types are described in Table 1.

The curator first adds genes from the publication, then creates alleles from genes, genotypes from alleles and metagenotypes from pathogen and host genotypes. Pathogen genotypes and host genotypes are created on separate pages, which only include genes from the relevant species. A genotype can consist of multiple alleles, and a metagenotype can contain multiple alleles from both the pathogen and the host. A ‘copy and edit’ feature allows the creation of multiple similar annotations.

To make annotations, the curator selects a gene, genotype, or metagenotype to annotate, then selects a term from a controlled vocabulary, adds experimental evidence, experimental conditions, AEs (where available), and any additional comments. In PHI-Canto, the curator can also specify a figure or table number from the original publication as part of the annotation. Curators can use PHI-Canto’s term suggestion feature to suggest new terms for any controlled vocabulary in PHI-Canto, and experimental conditions can be entered as free text if no suitable condition is found in PHI-ECO (new condition suggestions are reviewed and approved by expert curators). The curation session can be saved and paused at various stages in the entire process. Once the curation process is complete, the curator submits the session for review.

### Display and interoperability of data

The migration to incorporate FAIR principles fully into the PHI-base curation process will promote interoperability between various data resources (Wilkinson et al., 2016). Figure 5 illustrates the internal and external resource dependencies for curation in PHI-Canto. URLs and descriptions of the use of each resource are provided in Figure 5 – figure supplement 1. All data curated in PHI-Canto will be displayed in PHI-base version 5, introduced in (Urban et al., 2021). Additional detail on the data types displayed in PHI-base 5 is available in Table 4. Reciprocally, components of the interspecies curation framework (Figure 6a) provide data to other resources (Figure 6b). For example, GO terms will be used in curation with PHI-Canto and these annotations will be made available in the main GO database via the GOA Database. PHI-base is a member of ELIXIR, one of the leading organizations for biological resources and a major proponent of FAIR data.

**Figure 5.**
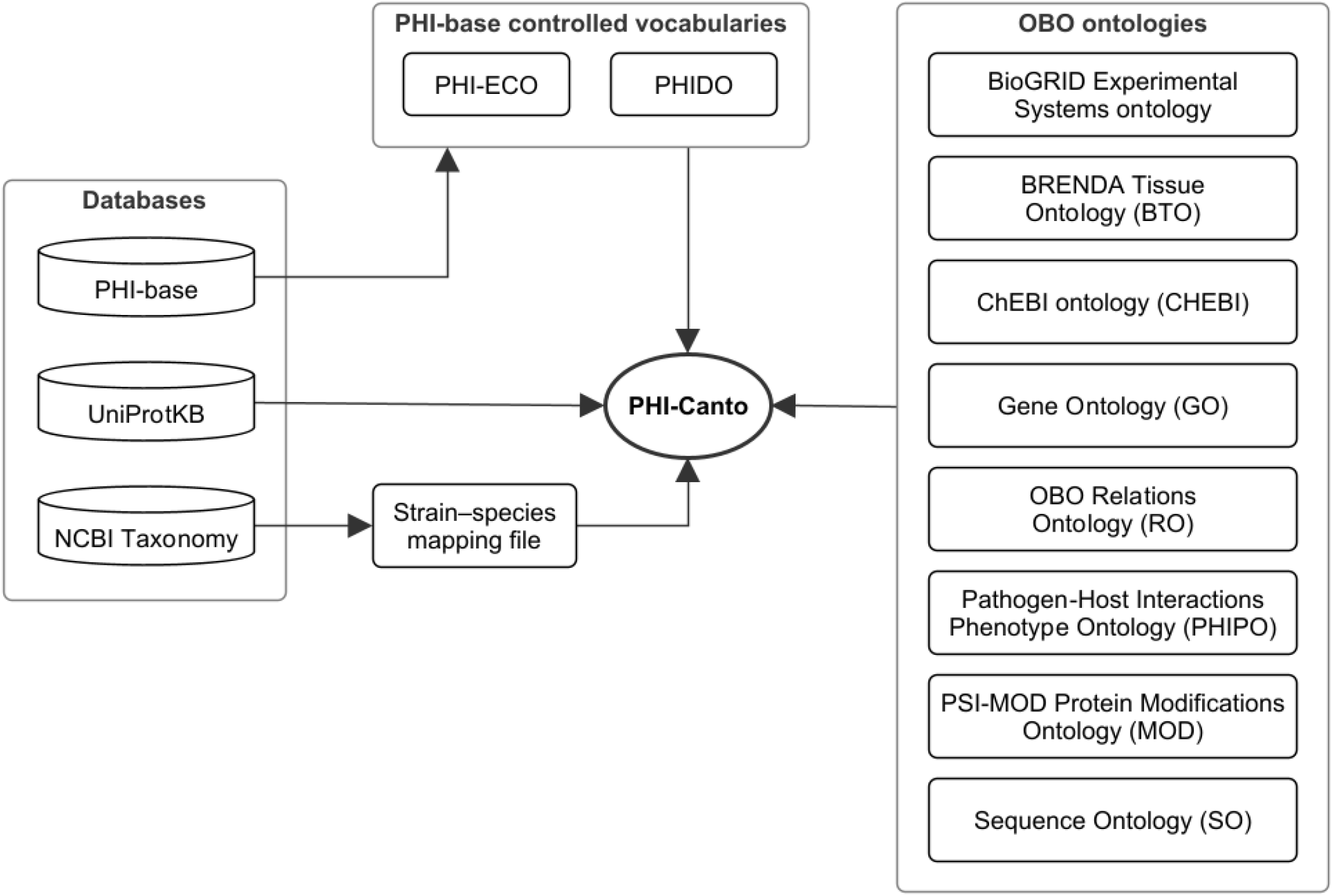
Network diagram showing the data resources used by PHI-Canto. Of the databases shown, PHI-base provides data (experimental conditions and disease names) used to create terms in the PHI-base controlled vocabularies; UniProtKB provides accession numbers for proteins that PHI-Canto uses to identify genes; and the NCBI Taxonomy database is used to generate a mapping file relating taxonomic identifiers lower than species rank to their nearest taxonomic identifiers at species rank. The OBO ontologies group contains ontologies in the OBO format that PHI-Canto uses for its annotation types. The parenthesized text after the ontology name indicates the term prefix for the ontology.

**Figure 6.**
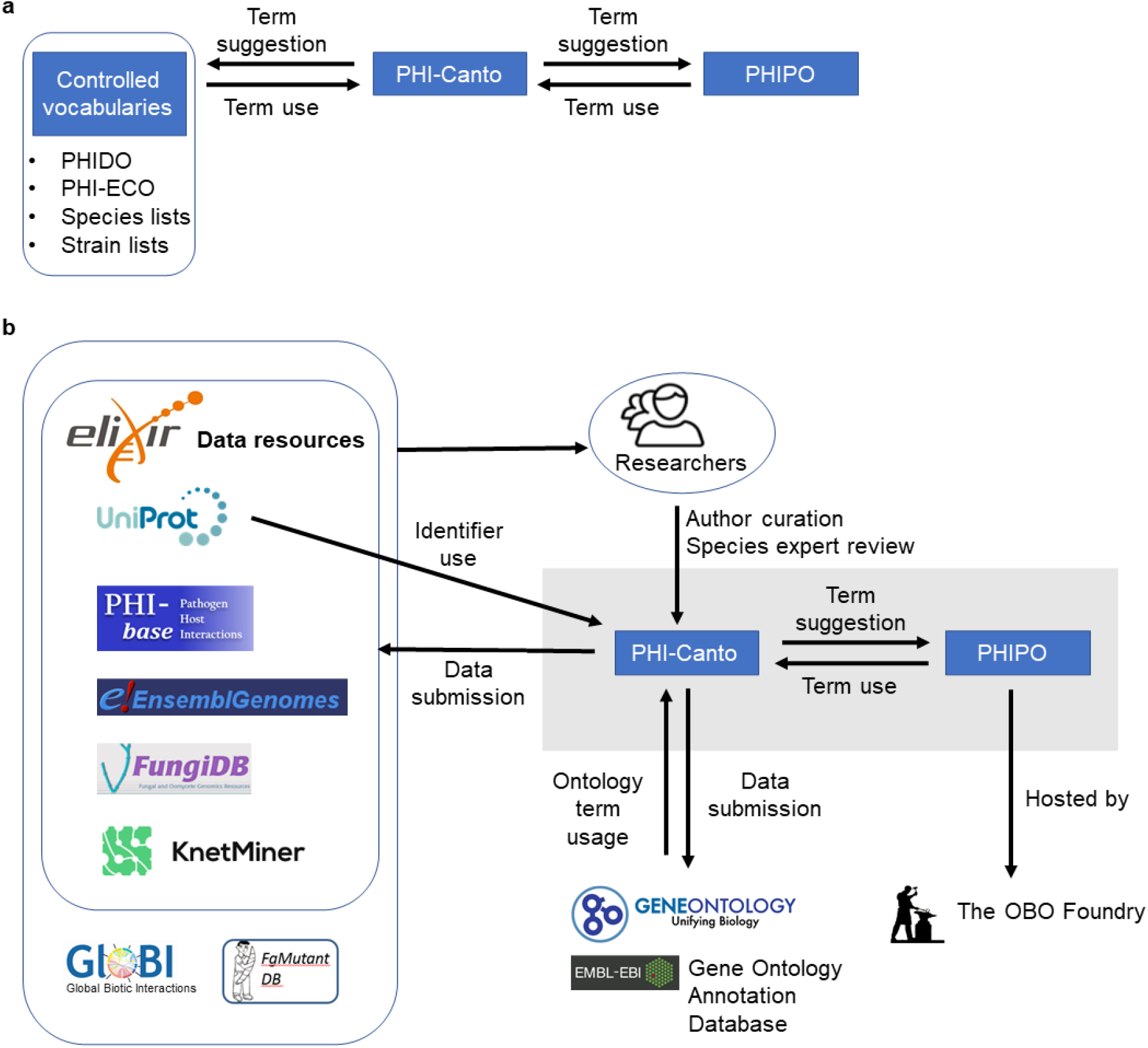
The interspecies curation framework and the interoperability of PHI-Canto. **(a)** The interspecies curation framework consists of three main components. Firstly, a curation tool called PHI-Canto (The Pathogen Host Interaction Community Annotation Tool), secondly, a new species neutral phenotype ontology called PHIPO (the Pathogen-Host Interaction Phenotype Ontology), and thirdly, a selection of additional controlled vocabularies for disease names (PHIDO), experimental conditions (PHI-ECO), pathogen and host species, and natural strains associated with each species. The two-way arrows indicate that terms from the ontology and controlled vocabularies are used in curation with PHI-Canto, and that new terms required for curation may be suggested for inclusion within the ontology and controlled vocabularies. **(b)** The PHI-Canto and PHIPO content curation framework (grey box) uses persistent identifiers and cross-referenced information from UniProt, Ensembl Genomes and the Gene Ontology. PHIPO is made available at the OBO Foundry. Newly minted wild type gene annotations are suggested for inclusion into the Gene Ontology via the EBI Gene Ontology Annotation database. Data curated in PHI-Canto will be shared with ELIXIR data resources such as UniProtKB, Ensembl Genomes, FungiDB, and KnetMiner, and will be provided on request to other databases (FgMutantDB, GloBI). Researchers can look up curated information via the PHI-base web interface or can download the whole dataset for inclusion in their bioinformatics pipelines. Authors can submit data to PHI-base by curating their publications into PHI-Canto. The origin of data is indicated by directional arrows.

**Table 4.** Automatically and manually curated types of data displayed in gene-centric PHI-base 5.

| Data type | Data source |
| --- | --- |
| **Metadata** |  |
| Entry Summary <sup>1</sup> | UniProtKB <sup>2</sup> |
| Pathogen species | NCBI Taxonomy <sup>2</sup> |
| Pathogen strain | PHI-base strain list |
| Host species | NCBI Taxonomy <sup>2</sup> |
| Host strain | PHI-base strain list |
| Publication | PubMed <sup>2</sup> |
| **Phenotype annotation sections** |  |
| Pathogen-Host Interaction Phenotype | PHIPO <sup>3</sup> pathogen-host interaction phenotype branch |
| Gene-for-Gene Phenotype | PHIPO pathogen-host interaction phenotype branch |
| Pathogen Phenotype | PHIPO single species phenotype branch |
| Host Phenotype | PHIPO single species phenotype branch |
| **Other annotation sections** |  |
| Disease name | PHIDO |
| GO Molecular Function | GO <sup>4</sup> |
| GO Biological Process | GO |
| GO Cellular Component | GO |
| Wild type RNA level | FYPO_EXT <sup>5</sup> |
| Wild type Protein level | FYPO_EXT |
| Physical Interaction | BioGRID <sup>6</sup> |
| Protein Modification | PSI-MOD <sup>7</sup> |

<sup>1</sup> The Entry Summary section includes information on which gene is being displayed in the gene-centric results page. The UniProtKB accession number is used to automatically retrieve the name and function of the protein, plus any cross-referenced identifiers from Ensembl Genomes and NCBI GenBank. The section also displays the PHI-base 5 gene identifier (PHIG) and any of the high-level terms (Supplementary file 3) annotated to the gene.
<sup>2</sup> Data from UniProtKB, NCBI Taxonomy and PubMed are automatically retrieved, while all other data are manually curated.
<sup>3</sup> PHIPO is the Pathogen-Host Interaction Phenotype Ontology.
<sup>4</sup> GO is the Gene Ontology.
<sup>5</sup> FYPO\_EXT is the Fission Yeast Phenotype Ontology Extension.
<sup>6</sup> BioGRID is the Biological General Repository for Interaction Datasets.
<sup>7</sup> PSI-MOD is the Human Proteome Organization (HUPO) Proteomics Standards Initiative (PSI) Protein Modifications Ontology.

## Discussion

Scalable and accurate curation of data within the scientific literature is of paramount importance due to the increasing quantity of publications and the complexity of experiments within each publication. PHI-base is an example of a freely available, manually curated database, which has been curating literature using professional curators since 2005 (Winnenburg et al., 2006). Here, we describe the development of PHI-Canto to allow the curation of the interspecies pathogen-host interaction literature by professional curators and publication authors. However, it should be noted that these developments – especially the concept of annotating metagenotypes – could be of use to communities focused on different types of interspecies interactions. Customizing Canto to use other ontologies and controlled vocabularies is as simple as editing a configuration file, as shown in Source code 1.

Several adaptations to the original single species community annotation tool, Canto (Rutherford et al., 2014), were required to convert this tool for interspecies use. Notably, the need to annotate an interaction involving two different organisms necessitated the development of a novel concept, the ‘metagenotype’, in order to record a combined experimental genotype involving both a pathogen and a host. This is, to our knowledge, the first example of such an approach to interspecies interaction curation.

Curation of pathogen-host interactions in PHI-Canto also necessitated the development of a new phenotype ontology (PHIPO) to annotate pathogen-host interaction phenotypes in sufficient detail across the broad range of host species that were curated in PHI-base. The functional annotation of genes involved in interspecies interactions is a complex and challenging task, requiring ongoing modifications to the Gene Ontology and occasionally major refactoring to deprecate legacy terms (Gene Ontology Consortium, 2021). PHIPO development and maintenance will also be an ongoing task, with both authors and professional curators requesting new terms and edits to existing terms and the ontology structure. Maintenance will be made more sustainable by the incorporation of logical definitions that are aligned across phenotype ontologies in collaboration with the uPheno project (Shefchek et al., 2020).

To improve the efficiency of the curation process, we are suggesting that authors follow an author checklist during manuscript preparation (Appendix 3). This will improve the key information (e.g., species names, gene identifiers etc.) in published manuscripts, thus enabling more efficient comprehensive curation that is both human- and machine-readable. The annotation procedures described here using PHI-Canto can be used to extract data buried in small-scale publications and increase the accessibility of the curated article to a wider range of potential users, for example computational biologists, thereby improving the FAIR status of the data. The current data in PHI-base has been obtained from > 200 journals (Figure 7) and therefore represents highly fragmented knowledge which is exceptionally difficult to use by professionals in other disciplines. The feasibility of scalable community curation with Canto is evidenced by PomBase, where Canto has been used by authors to curate ∼25% of the *S. pombe* literature, with the data being made available within 24 hours of curation review and approval (https://curation.pombase.org/pombe/stats/annotation).

**Figure 7.**
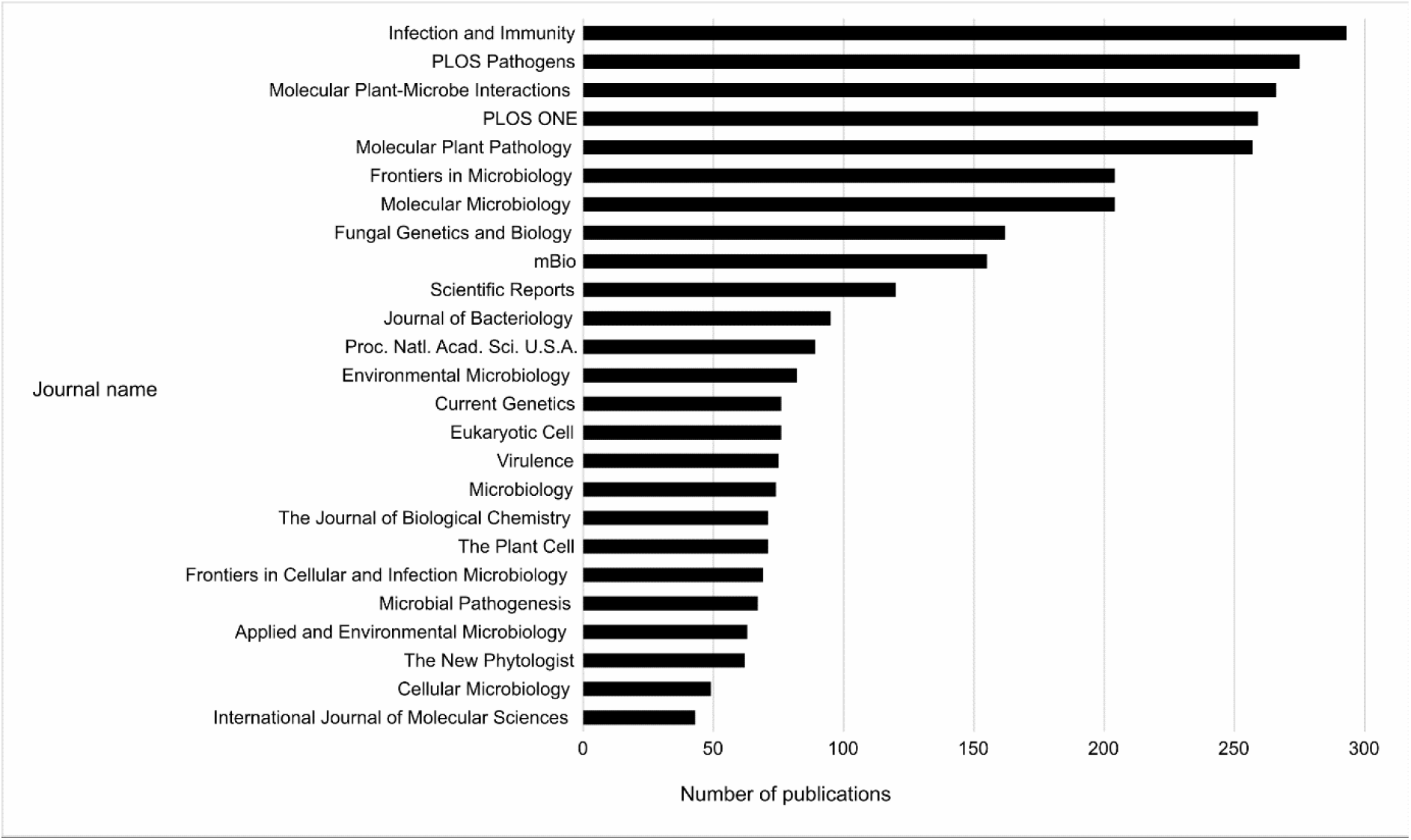
Top 25 Journals in PHI-base. Bar chart showing the top 25 journals by number of publications curated in PHI-base, as of version 4.13 (published 9 May 2022). Publication counts were generated by extracting every unique PubMed identifier (PMID) from PHI-base, then using the Entrez Programming Utilities (E-Utilities) to retrieve the journal name for each PMID, and finally summing the count of journal names.

Future plans for PHI-Canto include addressing issues with natural sequence variation between species strains. PHI-base contains information on numerous species with multiple experimental strains, and natural sequence variation between strains can result in alterations at the genome level that affect the subsequently observed phenotypes. Strain-specific sequence variation is not captured in the reference proteomes stored by UniProt, even though accession numbers from these proteomes are often used in PHI-Canto. Currently, when a curator enters a gene with a taxonomic identifier below the species rank, PHI-Canto maps the identifier to the corresponding identifier at the species rank (thus removing any strain details from the organism name), and the curator specifies a strain to differentiate gene variants in naturally occurring strains. However, this does not change the taxonomic identifier linked to the UniProtKB accession number (nor its sequence), so the potential for inaccuracy remains. To mitigate this, the future plan is to record the strain-specific sequence of the gene using an accession number from a database from the International Nucleotide Sequence Database Collaboration (Arita, Karsch-Mizrachi, & Cochrane, 2021).

The release of PHI-Canto to the community will occur gradually through various routes. Community curation will be promoted by working with journals to capture the publication data at source, at the point of manuscript acceptance. We will also target specific research communities (e.g., those working on a particular pathogen) by inviting authors to curate their own publications. Authors may contact us directly to request support while curating their publications in PHI-Canto.

## Methods

### Changes to the Canto data model and configuration

Several new entities were added to PHI-Canto’s data model in order to support pathogen-host curation, as well as new configuration options (the new entities are illustrated in Figure 3 – figure supplement 1).

#### Pathogen and host roles

Genotype entities in PHI-Canto’s data model were extended with an attribute indicating their status as a pathogen genotype or a host genotype. Genotypes inherit their status (as pathogen or host) from the organism, which in turn is classified as a pathogen or host based on a configuration file that contains the NCBI Taxonomy ID (taxid) (Schoch et al., 2020) of each host species in PHI-base. Only host taxids need to be specified since PHI-Canto defaults to classifying a species as a pathogen if its taxid is not found in the configuration file.

PHI-Canto also loads lists of pathogen and host species that specify the scientific name, taxid, and common name (if any) of each species. These species lists are used to specify which host species can be added as a component of the metagenotype in the absence of a specific studied gene, and to override the scientific name provided by UniProtKB in favor of the name used by the community (for example, to control whether the anamorph or teleomorph name of a fungal species is displayed in PHI-Canto’s user interface).

#### Metagenotype implementation

Metagenotypes were implemented by adding a ‘metagenotype’ entity to PHI-Canto’s data model. The metagenotype is the composition of two genotype entities. We also introduced new relations into the data model to allow annotations to be related to metagenotypes (previously, only genes and genotypes could be related to annotations).

#### Strain implementation

Support for strain curation was implemented by adding a ‘strain’ entity to PHI-Canto’s data model. Strains are related to an organism entity and its related genotype entities. In the user interface, PHI-Canto uses the taxid of the organism to filter an autocomplete system, such that only the strains of the specified organism are suggested. The autocomplete system can also use synonyms in the strain list to suggest a strain based on its synonymous names. Unknown strains are represented by a preset value of ‘Unknown strain’.

### Ontologies

PHIPO was developed using the Protégé ontology editor (Musen & Protégé Team, 2015). PHIPO uses OBO namespaces to allow PHI-Canto to filter the terms in the ontology by annotation type, ensuring that genotypes are annotated with single-species phenotypes and metagenotypes with pathogen-host interaction phenotypes.

PHI-ECO was developed using Protégé, starting from a list of experimental conditions originally developed by PomBase. PHIDO was initially derived from a list of diseases already curated in PHI-base and is now maintained as a flat file that is converted into an OBO file using ROBOT (Jackson et al., 2019).

## Data availability

Pathogen-Host Interaction Phenotype Ontology: http://purl.obolibrary.org/obo/phipo.owl

PHI-base Experimental Conditions Ontology: https://github.com/PHI-base/phi-eco

PHIDO, the controlled vocabulary of disease names: https://github.com/PHI-base/phido

PHIPO Extension Ontology for gene-for-gene phenotypes: https://github.com/PHI-base/phipo_ext

Location of species and strain lists used by PHI-Canto: https://github.com/PHI-base/data

PHI-Canto approved curation sessions (December 2022): https://doi.org/10.5281/zenodo.7428788

## Code availability

PHI-Canto’s source code is available on GitHub, at https://github.com/PHI-base/canto. PHI-Canto is freely licensed under the GNU General Public License version 3, with no restrictions on copying, distributing, or modifying the code, for commercial use or otherwise, provided any derivative works are licensed under the same terms. PHI-base provides an online demo version of PHI-Canto at https://demo-canto.phi-base.org/ which can be used for evaluating the tool. The demo version and the main version of PHI-Canto will remain freely available online for the foreseeable future.

Canto’s source code is available on GitHub, at https://github.com/pombase/canto. Canto is also freely licensed under the GNU General Public License version 3.

The source code for PHI-Canto’s user documentation is available on GitHub, at https://github.com/PHI-base/canto-docs. The user documentation is available online at https://canto.phi-base.org/docs/index.

The source code for PHIPO is available on GitHub under a Creative Commons Attribution 3.0 license, at https://github.com/PHI-base/phipo.

## Acknowledgements

We thank former post-doctoral PHI-base team member Dr Alistair Irvine for adding chemical entries to ChEBI. Dr Paul Kersey, formerly the non-vertebrate Ensembl team leader, is thanked for helpful discussions and ideas on community engagement. We thank Dr Midori Harris (formerly of University of Cambridge, UK) for providing valuable input into the development of PHIPO based on her extensive knowledge of FYPO. Dr Pascale Gaudet (Swiss-Prot, Swiss Institute of Bioinformatics) is thanked for the generation and editing of GO terms involved in interspecies interactions. We also thank Drs Chris Stephens and Ana Machado-Wood (both formerly of Rothamsted Research) for completing the trial curation of articles into beta versions of PHI-Canto and providing invaluable feedback and suggestions for further improvements. The Molecular Connections team based in Bangalore India while developing the PHI-base 5 website, provided useful feedback on data interoperability between PHI-Canto and the new gene-centric version of PHI-base.

## Author’s contributions

AC wrote the initial manuscript draft. JS, VW, KR, MU and KHK provided comments on various manuscript versions. AC, JS, MU and KHK prepared the figures and tables. AC and JS prepared the supplementary files.

## Corresponding authors

Correspondence to.

## Figure supplements

**Figure 3 – figure supplement 1.** Canto entity relationship diagram.

Simplified UML class diagram showing the relations between entities (things of interest) in a Canto curation session. The numbers on the connecting lines represent the cardinality of the relation, meaning how many of one entity can be related to another entity: 0..n means ‘zero or more’; 1..n means ‘one or more’. Lines with a hollow arrowhead indicate that the target entity (at the head of the arrow) is a generalization of the source entity (at the tail of the arrow). Boxes outlined in bold indicate new entities which were added to support curation in PHI-Canto.

**Figure 4 – figure supplement 1.** Alternative curation step workflow.

The flow diagram represents the PHI-Canto curation process from beginning to end in 5 steps. It is an alternative representation to the image depicted in Figure 4. During step 2 of the workflow, the curator chooses either the gene annotation or genotype / metagenotype annotation process. Multiple annotations can be made using both annotation processes which can then be submitted for review.

**Figure 4 - figure supplement 2.** What you need to curate a publication into PHI-Canto.

**Figure 4 - figure supplement 3.** Instructions on how to look up a UniProtKB ID.

**Figure 5 – figure supplement 1**. Resources relied upon by PHI-Canto.

## Source code

**Source code 1.** Main configuration file for PHI-Canto.

This is the main configuration file for PHI-Canto. Much of the configuration is inherited from Canto, the original curation application from which PHI-Canto is derived. Lines containing custom configuration for PHI-Canto have been indicated with comments.

## Appendix 1 How to use Annotation Extensions

This file provides information on Annotation Extensions (AE) and how to use them in PHI-Canto to curate a standard selection of experiments (Table 2). The first section provides four examples of using AEs for curating metagenotypes with pathogen-host interaction phenotypes. The second section provides examples of curating metagenotypes using the gene-for-gene phenotype workflow, including using the AEs for gene-for-gene interactions and inverse gene-for-gene interactions. The third section of this file illustrates three examples of using AEs for curating single species phenotypes.

Further information on how to use PHI-Canto to make annotations can be found in PHI-Canto’s user documentation, available at https://canto.phi-base.org/docs/index.

### SECTION 1 Annotation Extensions for curating pathogen-host interaction phenotypes on metagenotypes

When creating and annotating metagenotypes, it is advisable to also create and annotate a wild type control metagenotype where possible. This enables a better understanding of annotations made to altered metagenotypes.

(Note: it is also possible to use several of the AEs in the table documenting single species phenotype AEs, e.g. *penetrance* and *affected protein*)

**If you have a metagenotype phenotype recording ‘unaffected pathogenicity’ (corresponds to footnote 1 in Table 2**)

AE summary table

| AE name | Cardinality | Available terms |
| --- | --- | --- |
| compared to control genotype | 0, 1 | Metagenotype identifier |
| extent of infectivity | 0, 1 | 'unaffected pathogenicity' |
| host tissue affected | 0, *n* | BRENDA Tissue Ontology term |
| outcome of interaction | 0, 1 | 'disease present', 'disease absent' |

Example publication: The RhlR quorum-sensing receptor controls *Pseudomonas aeruginosa* pathogenesis and biofilm development independently of its canonical homoserine lactone autoinducer. (<u>PMID:28715477</u>)

#### Pathogen-host interaction phenotype

##### Control metagenotype

| Pathogen genotype | Host genotype | Term ID | Term name | Evidence code | Conditions | Figure | Annotation extension |
| --- | --- | --- | --- | --- | --- | --- | --- |
| rhII+[WT level]<br>*P. aeruginosa*<br>(PA14) | wild type<br>*C. elegans*<br>(N2) | PHIPO:0001069 | death of host organism with pathogen | Cell growth assay | agar plates | Fig 6a | **infects_tissue** whole body ,<br>**has_penetrance** 50% ,<br>**interaction_outcome** disease present |

##### Altered metagenotype

| Pathogen genotype | Host genotype | Term ID | Term name | Evidence code | Conditions | Figure | Annotation extension |
| --- | --- | --- | --- | --- | --- | --- | --- |
| rhIIΔ<br>*P. aeruginosa*<br>(PA14) | wild type<br>*C. elegans*<br>(N2) | PHIPO:0001069 | death of host organism with pathogen | Cell growth assay | agar plates | Fig 6a | **infects_tissue** whole body , **infective_ability** unaffected pathogenicity , **has_penetrance** 50% , **compared_to_control** rhII+[WT level] *Pseudomonas aeruginosa* (PA14) / wild type *Caenorhabditis elegans* (N2) , **interaction_outcome** disease present |

**If you have a metagenotype phenotype recording ‘altered pathogenicity or virulence’ (corresponds to footnote 2 in Table 2**)

#### AE summary table

| AE name | Cardinality | Available terms |
| --- | --- | --- |
| compared to control genotype | 0, 1 | Metagenotype identifier |
| extent of infectivity | 0, 1 | 'loss of pathogenicity', 'reduced virulence', 'increased virulence' |
| host tissue affected | 0, *n* | BRENDA Tissue Ontology term |
| outcome of interaction | 0, 1 | 'disease present', 'disease absent' |

Example publication: A conserved fungal glycosyltransferase facilitates pathogenesis of plants by enabling hyphal growth on solid surfaces (<u>PMID:29020037</u>)

A training video is available for the curation of this publication at https://youtu.be/44XGoi6Ijqk?t=1738

#### Pathogen-host interaction phenotype

##### Control metagenotype

| Pathogen genotype | Host genotype | Term ID | Term name | Evidence code | Conditions | Figure | Annotation extension |
| --- | --- | --- | --- | --- | --- | --- | --- |
| GT2+[WT level]<br>*Z. tritici*<br>(IPO323) | wild type<br>*T. aestivum*<br>(cv. Riband) | PHIPO:0000480 | presence of pathogen-associated host lesions | Macroscopic observation (qualitative observation) | 14 days post inoculation | Figure 2E | **infects_tissue** leaf , **interaction_outcome** disease present |

##### Altered metagenotype

| Pathogen genotype | Host genotype | Term ID | Term name | Evidence code | Conditions | Figure | Annotation extension |
| --- | --- | --- | --- | --- | --- | --- | --- |
| ΔGT2-19(deletion)<br>*Z. tritici*<br>(IPO323) | wild type<br>*T. aestivum*<br>(cv. Riband) | PHIPO:0000481 | absence of pathogen-associated host lesions | Macroscopic observation (qualitative observation) | 14 days post inoculation | Figure 2E | **infects_tissue** leaf , **infective_ability** loss of pathogenicity , **compared_to_control** GT2+[WT level] *Zymoseptoria tritici* (IPO323) / wild type *Triticum aestivum* (cv. Riband) , **interaction_outcome** disease absent |

**If you have a metagenotype phenotype recording ‘mutualism’ (corresponds to footnote 4 in Table 2**)

#### AE summary table

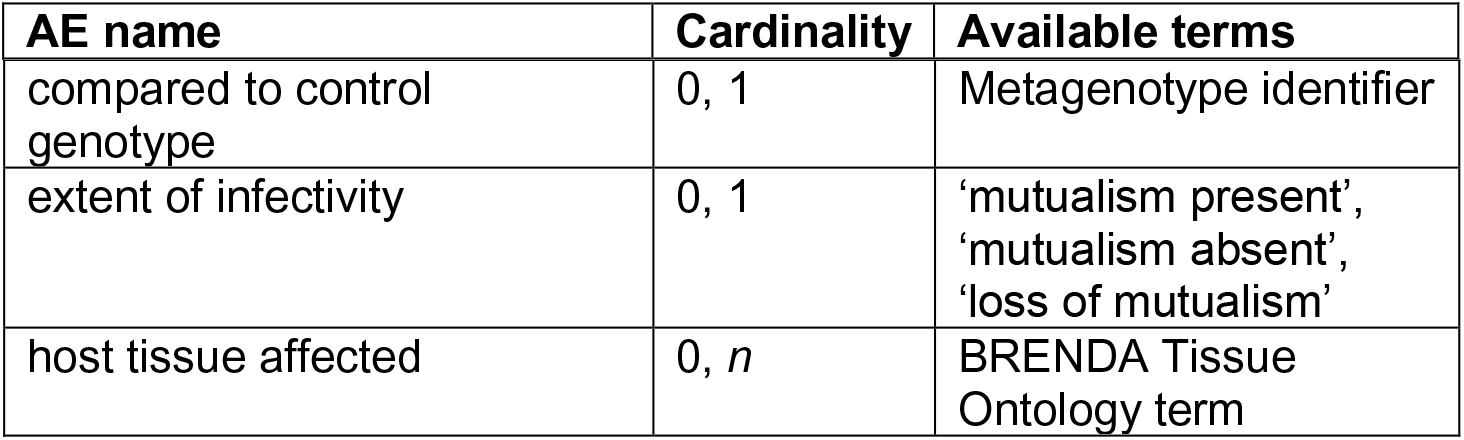

Note: The ‘Outcome of interaction’ AE is not relevant in this mutualism interaction.

Example publication: Reactive oxygen species play a role in regulating a fungus-perennial ryegrass mutualistic interaction (<u>PMID:16517760</u>)

#### Pathogen-host interaction phenotype: Example 1

Illustrating a phenotype associated with the <u>pathogen</u> component within the Pathogen-Host Interaction.

##### Control metagenotype

| Pathogen genotype | Host genotype | Term ID ↕ | Term name ↕ | Evidence code ↕ | Figure ↕ | Annotation extension ↕ |
| --- | --- | --- | --- | --- | --- | --- |
| noxA+[WT level]<br>*E. festucae*<br>(F11)<br>bkg: GFP | wild type<br>*L. perenne*<br>(Unknown strain) | PHIPO:0000954 | presence of pathogen growth within host | Microscopy | Figure 1c | **infects_tissue** leaf , **infective_ability** mutualism present |

##### Altered metagenotype

| Pathogen genotype | Host genotype | Term ID ↕ | Term name ↕ | Evidence code ↕ | Figure ↕ | Annotation extension ↕ |
| --- | --- | --- | --- | --- | --- | --- |
| noxA::pAN7-1(disruption)[Not assayed]<br>*E. festucae*<br>(F11)<br>bkg: GFP | wild type<br>*L. perenne*<br>(Unknown strain) | PHIPO:0000368 | increased pathogen growth within host | Microscopy | Figure 1d | **infects_tissue** leaf , **infective_ability** loss of mutualism , **compared_to_control** noxA+[WT level] *Epichloe festucae* (F11) / wild type *Lolium perenne* (Unknown strain) |

#### Pathogen-host interaction phenotype: Example 2

Illustrating a phenotype associated with the <u>host</u> component within the Pathogen-Host Interaction.

##### Control metagenotype

| Pathogen genotype | Host genotype | Term ID ↕ | Term name ↕ | Evidence code ↕ | Figure ↕ | Annotation extension ↕ |
| --- | --- | --- | --- | --- | --- | --- |
| noxA+[WT level]<br>*E. festucae*<br>(F11) | wild type<br>*L. perenne*<br>(Unknown strain) | PHIPO:0001005 | normal host morphology during pathogen invasion | Macroscopic observation (qualitative observation) | Figure 1a, 5c | **infects_tissue** whole plant , **infective_ability** mutualism present |

##### Altered metagenotype

Note: in this case, two separate annotations were made to the same metagenotype.

| Pathogen genotype | Host genotype | Term ID | Term name | Evidence code | Figure | Annotation extension |
| --- | --- | --- | --- | --- | --- | --- |
| noxA::pAN7-1(disruption)[Not assayed]<br>*E. festucae*<br>(F11) | wild type<br>*L. perenne*<br>(Unknown strain) | PHIPO:0001130 | stunted host growth during pathogen colonization | Macroscopic observation (qualitative observation) | Figure 1a | **infects_tissue** whole plant , **infective_ability** loss of mutualism , **compared_to_control** noxA+[WT level] *Epichloe festucae* (F11) / wild type *Lolium perenne* (Unknown strain) |
| noxA::pAN7-1(disruption)[Not assayed]<br>*E. festucae*<br>(F11) | wild type<br>*L. perenne*<br>(Unknown strain) | PHIPO:0001131 | increased number of host side shoots during pathogen colonization | Macroscopic observation (qualitative observation) | Figure 1a | **infects_tissue** whole plant , **infective_ability** loss of mutualism , **compared_to_control** noxA+[WT level] *Epichloe festucae* (F11) / wild type *Lolium perenne* (Unknown strain) |

**If you have a metagenotype phenotype recording ‘a pathogen effector’ (corresponds to footnote 5 in Table 2**)

If you have a biotrophic or necrotrophic plant pathogen effector which is involved in a gene-for-gene interaction, please see the AEs for the ‘gene-for-gene interaction’ or ‘inverse gene-for-gene interaction’ workflow (Section 2).

Annotate the pathogen effector with the GO Biological Process term ‘effector-mediated modulation of host process by symbiont’ (<u>GO:0140418</u>) or a descendant. If the GO Molecular Function term is known, then this can also be annotated and linked to the relevant GO effector term via an annotation extension.

#### AE summary table

| AE name | Cardinality | Available terms |
| --- | --- | --- |
| compared to control genotype | 0, 1 | Metagenotype identifier |
| extent of infectivity | 0, 1 | 'unaffected pathogenicity',<br>'loss of pathogenicity',<br>'reduced virulence',<br>'increased virulence' |
| host tissue affected | 0, *n* | BRENDA Tissue Ontology term |
| outcome of interaction | 0, 1 | 'disease present',<br>'disease absent' |

Example publication: An effector protein of the wheat stripe rust fungus targets chloroplasts and suppresses chloroplast function. (<u>PMID:31804478</u>)

#### GO biological process

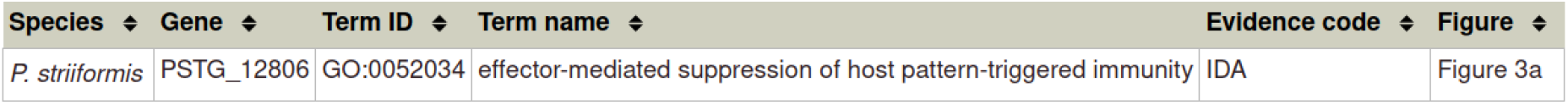

Note: ‘effector-mediated suppression of host pattern-triggered immunity’ (<u>GO:0052034</u>) is a descendant term of ‘effector-mediated modulation of host process by symbiont’ (<u>GO:0140418</u>).

#### GO molecular function

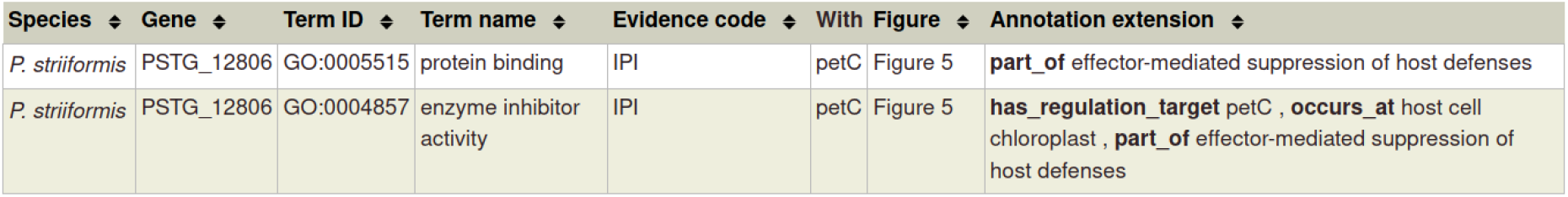

Please note that in the case of a physical interaction (protein–protein interaction) between the pathogen and host gene products (PSTG_12806 and PetC in the example above, respectively) this information can be curated using the Physical Interaction curation workflow, documented in https://canto.phi-base.org/docs/physical_interaction_annotation.

#### Pathogen-host interaction phenotype

##### Control metagenotype

In this case, there are no metagenotype control annotations. This is because it is not possible to create and annotate a metagenotype comprising of an empty vector control within the pathogen component of the metagenotype.

##### Altered metagenotype

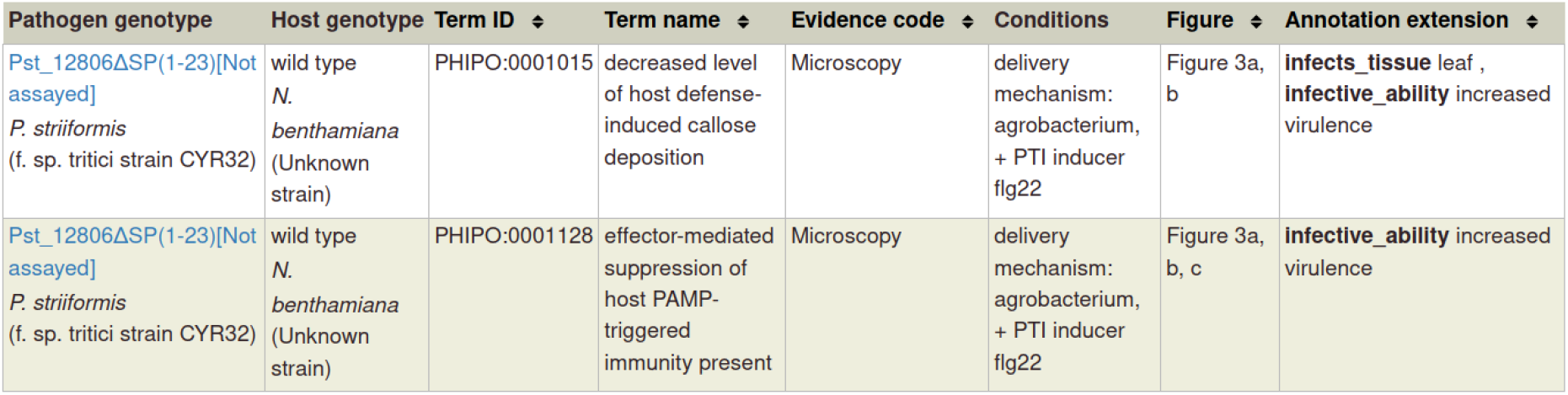

| Pathogen genotype | Host genotype | Term ID | Term name | Evidence code | Conditions | Figure | Annotation extension |
| --- | --- | --- | --- | --- | --- | --- | --- |
| Pst_12806ΔSP(1-23)[Not assayed]<br>*P. striiformis*<br>(f. sp. tritici strain CYR32) | wild type<br>*N. benthamiana*<br>(Unknown strain) | PHIPO:0001015 | decreased level of host defense-induced callose deposition | Microscopy | delivery mechanism: agrobacterium, + PTI inducer flg22 | Figure 3a, b | **infects_tissue** leaf , **infective_ability** increased virulence |
| Pst_12806ΔSP(1-23)[Not assayed]<br>*P. striiformis*<br>(f. sp. tritici strain CYR32) | wild type<br>*N. benthamiana*<br>(Unknown strain) | PHIPO:0001128 | effector-mediated suppression of host PAMP-triggered immunity present | Microscopy | delivery mechanism: agrobacterium, + PTI inducer flg22 | Figure 3a, b, c | **infective_ability** increased virulence |

### SECTION 2 Annotation Extensions for curating gene-for-gene phenotypes on metagenotypes

**If you have a metagenotype phenotype recording ‘a gene-for-gene interaction’ (corresponds to footnote 6 in Table 2**)

Annotate the pathogen effector with the GO Biological process term ‘effector-mediated modulation of host process by symbiont’ (<u>GO:0140418</u>) or a descendant. If the GO Molecular Function term is known, then this can also be annotated and linked to the relevant GO effector term via an annotation extension.

#### AE summary table

| AE name | Cardinality | Available terms |
| --- | --- | --- |
| compared to control genotype | 0, 1 | Metagenotype identifier |
| gene-for-gene phenotype | 0, 1 | 'incompatible interaction, recognizable pathogen effector present, functional host resistance gene present' |
|  |  | 'incompatible interaction, recognizable pathogen effector present, gain of functional host resistance gene'<br>'incompatible interaction, gain of recognizable pathogen effector, gain of functional host resistance gene'<br>'incompatible interaction, gain of recognizable pathogen effector, functional host resistance gene present'<br>'compatible interaction, recognizable pathogen effector present, functional host resistance gene absent'<br>'compatible interaction, recognizable pathogen effector absent, functional host resistance gene present'<br>'compatible interaction, recognizable pathogen effector present, compromised host resistance gene'<br>'compatible interaction, recognizable pathogen effector absent, functional host resistance gene absent'<br>'compatible interaction, recognizable pathogen effector absent, compromised functional host resistance gene' |
|  |  | 'compatible interaction, compromised recognizable pathogen effector, functional host resistance gene present'<br>'metagenotype outcome overcome by external condition' |
| host tissue affected | 0, *n* | BRENDA Tissue Ontology term |

Example publication: Activation of an Arabidopsis resistance protein is specified by the in planta association of its leucine-rich repeat domain with the cognate oomycete effector. (<u>PMID:20601497</u>).

#### GO biological process

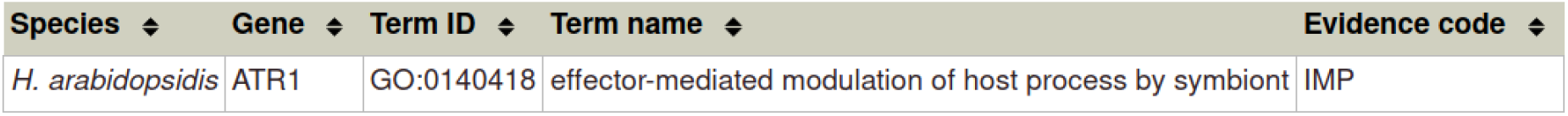

| Species ↕ | Gene ↕ | Term ID ↕ | Term name ↕ | Evidence code ↕ |
| --- | --- | --- | --- | --- |
| *H. arabidopsidis* | ATR1 | GO:0140418 | effector-mediated modulation of host process by symbiont | IMP |

#### GO molecular function

| Species ↕ | Gene ↕ | Term ID ↕ | Term name ↕ | Evidence code ↕ | With | Figure ↕ | Annotation extension ↕ |
| --- | --- | --- | --- | --- | --- | --- | --- |
| *H. arabidopsidis* | ATR1 | GO:0005515 | protein binding | IPI | RPP1 | Figure 4, 5 | **part_of** effector-mediated modulation of host process by symbiont |

#### Gene-for-gene phenotype

##### Control metagenotypes

###### Incompatible control

| Pathogen genotype | Host genotype | Term ID ↕ | Term name ↕ | Evidence code ↕ | Conditions | Figure ↕ | Annotation extension ↕ |
| --- | --- | --- | --- | --- | --- | --- | --- |
| ATR1-Δ51(1-51)[Not assayed]<br>*H. arabidopsidis* (Maks9)<br>**bkg**: Citrine tag | RPP1+[Not assayed]<br>*A. thaliana* (ecotype Ws-0)<br>**bkg**: HA tag | PHIPO:0000192 | presence of host-defense induced lesion by host hypersensitive response | Macroscopic observation (qualitative observation) | delivery mechanism: agrobacterium, heterologous species tobacco | Figure 3a | **infects_tissue** leaf , **gene_for_gene_interaction** incompatible interaction, recognizable pathogen effector present, functional host resistance gene present |

###### Compatible control

| Pathogen genotype | Host genotype | Term ID ↕ | Term name ↕ | Evidence code ↕ | Conditions | Figure ↕ | Annotation extension ↕ |
| --- | --- | --- | --- | --- | --- | --- | --- |
| ATR1-Δ51(1-51)[Not assayed]<br>*H. arabidopsidis* (Maks9)<br>**bkg**: Citrine tag | RPP1+[Not assayed]<br>*A. thaliana* (ecotype Nd-0)<br>**bkg**: HA tag | PHIPO:0000182 | absence of host-defense induced lesion by host hypersensitive response | Macroscopic observation (qualitative observation) | delivery mechanism: agrobacterium, heterologous species tobacco | Figure 3b, 8b | **infects_tissue** leaf , **gene_for_gene_interaction** compatible interaction, recognizable pathogen effector absent, functional host resistance gene present |

#### Altered metagenotype (shift from compatible to incompatible interaction)

| Pathogen genotype | Host genotype | Term ID | Term name | Evidence code | Conditions | Figure | Annotation extension |
| --- | --- | --- | --- | --- | --- | --- | --- |
| ATR1-Δ51-D191G(1-51, D191G)[Not assayed]<br>*H. arabidopsidis* (Maks9)<br>**bkg**: Citrine tag | RPP1+[Not assayed]<br>*A. thaliana* (ecotype Nd-0)<br>**bkg**: HA tag | PHIPO:0000192 | presence of host-defense induced lesion by host hypersensitive response | Macroscopic observation (qualitative observation) | delivery mechanism: agrobacterium, heterologous species tobacco | Figure 8b | **infects_tissue** leaf , **gene_for_gene_interaction** incompatible interaction, gain of recognizable pathogen effector, functional host resistance gene present , **compared_to_control** ATR1-delta51(1-51)[Not assayed] Hyaloperonospora arabidopsidis (Maks9) / RPP1+[Not assayed] Arabidopsis thaliana (ecotype Nd-0) |

**If you have a metagenotype phenotype recording an inverse gene-for-gene interaction (corresponds to footnote 8 in Table 2**)

Annotate the pathogen effector with the GO Biological process term ‘effector-mediated modulation of host process by symbiont’ (<u>GO:0140418</u>) or a descendant. If the GO Molecular Function term is known, then this can also be annotated and linked to the relevant GO effector term via an annotation extension.

#### AE summary table

| AE name | Cardinality | Available terms |
| --- | --- | --- |
| compared to control genotype | 0, 1 | Metagenotype identifier |
| inverse gene-for-gene phenotype | 0, 1 | 'compatible interaction, functional pathogen necrotrophic effector present, functional host susceptibility locus present'<br>'compatible interaction, functional pathogen necrotrophic effector present, gain of functional host susceptibility locus'<br>'compatible interaction, gain of functional pathogen necrotrophic effector, functional host susceptibility locus present'<br>'incompatible interaction, functional pathogen necrotrophic effector present, functional host susceptibility locus absent'<br>'incompatible |
|  |  | interaction, functional pathogen necrotrophic effector absent, functional host susceptibility locus present'<br>'incompatible interaction, functional pathogen necrotrophic effector present, functional host susceptibility locus compromised'<br>'incompatible interaction, compromised functional pathogen necrotrophic effector, functional host susceptibility locus present'<br>'incompatible interaction, gain of functional pathogen necrotrophic effector, functional host susceptibility locus compromised'<br>'metagenotype outcome overcome by external condition' |
| host tissue affected | 0, *n* | BRENDA Tissue Ontology term |

Example publication: The cysteine rich necrotrophic effector SnTox1 produced by Stagonospora nodorum triggers susceptibility of wheat lines harboring Snn1. (<u>PMID:22241993</u>).

#### GO biological process

| Species ↕ | Gene ↕ | Term ID ↕ | Term name ↕ | Evidence code ↕ |
| --- | --- | --- | --- | --- |
| *P. nodorum* | Tox1 | GO:0080185 | effector-mediated induction of plant hypersensitive response by symbiont | EXP |

#### GO molecular function

| Species ↕ | Gene ↕ | Term ID ↕ | Term name ↕ | Evidence code ↕ | Annotation extension ↕ |
| --- | --- | --- | --- | --- | --- |
| *P. nodorum* | Tox1 | GO:0140295 | pathogen-derived receptor ligand activity | EXP | has_input Snn1 , part_of effector-mediated induction of plant hypersensitive response by symbiont |

#### Gene-for-gene phenotype

##### Control metagenotypes

###### Compatible control

| Pathogen genotype | Host genotype | Term ID ↕ | Term name ↕ | Evidence code ↕ | Conditions | Figure ↕ | Annotation extension ↕ |
| --- | --- | --- | --- | --- | --- | --- | --- |
| Tox1+[WT level]<br>*P. nodorum*<br>(SN15) | Snn1+[WT level]<br>*T. aestivum*<br>(cv. Chinese Spring) | PHIPO:0000480 | presence of pathogen-associated host lesions | Macroscopic observation (qualitative observation) | delivery mechanism: culture infiltration | Figure 1 | **infects_tissue** leaf , **inverse_gene_for_gene** compatible interaction, functional pathogen necrotrophic effector present, functional host susceptibility locus present |

###### Incompatible control

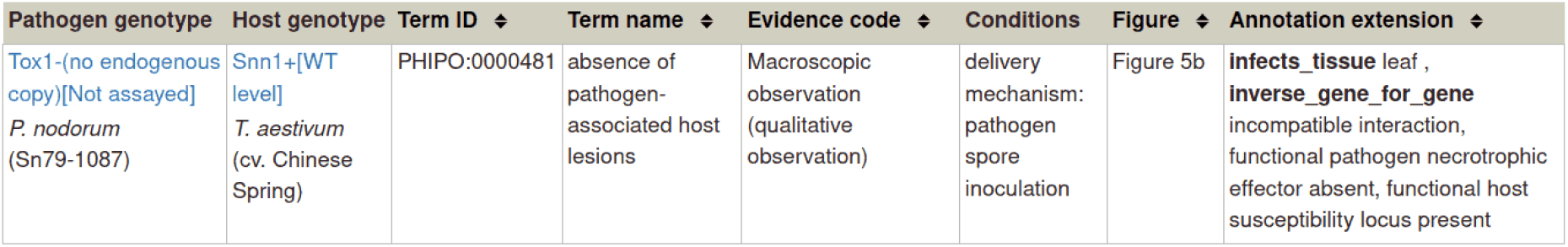

| Pathogen genotype | Host genotype | Term ID ↕ | Term name ↕ | Evidence code ↕ | Conditions | Figure ↕ | Annotation extension ↕ |
| --- | --- | --- | --- | --- | --- | --- | --- |
| Tox1-(no endogenous copy)[Not assayed]<br>*P. nodorum*<br>(Sn79-1087) | Snn1+[WT level]<br>*T. aestivum*<br>(cv. Chinese Spring) | PHIPO:0000481 | absence of pathogen-associated host lesions | Macroscopic observation (qualitative observation) | delivery mechanism: pathogen spore inoculation | Figure 5b | **infects_tissue** leaf , **inverse_gene_for_gene** incompatible interaction, functional pathogen necrotrophic effector absent, functional host susceptibility locus present |

##### Altered metagenotypes

###### Shift from compatible to incompatible interaction

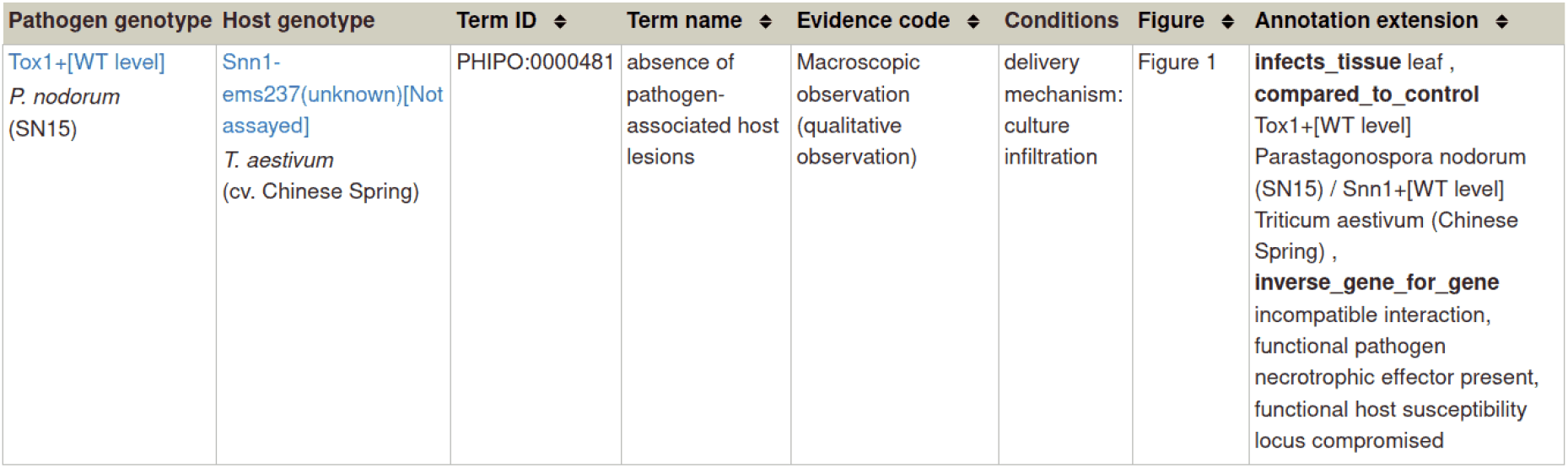

###### Shift from incompatible to compatible interaction

| Pathogen genotype | Host genotype | Term ID ↕ | Term name ↕ | Evidence code ↕ | Conditions | Figure ↕ | Annotation extension ↕ |
| --- | --- | --- | --- | --- | --- | --- | --- |
| +Sn15Tox1A1(transformant, no endogenous copy)[Not assayed]<br>*P. nodorum*<br>(Sn79-1087) | Snn1+[WT level]<br>*T. aestivum*<br>(cv. Chinese Spring) | PHIPO:0000480 | presence of pathogen-associated host lesions | Macroscopic observation (qualitative observation) | delivery mechanism: pathogen spore inoculation | Figure 5b | **infects_tissue** leaf , **compared_to_control** Tox1-(no endogenous copy)[Not assayed] *Parastagonospora nodorum* (Sn79-1087) / *Snn1+[WT level]* *Triticum aestivum* (Chinese Spring) , **inverse_gene_for_gene** compatible interaction, gain of functional pathogen necrotrophic effector, functional host susceptibility locus present |

###### No shift compared to control, still an incompatible interaction, despite alteration to both pathogen and host genotypes

Note: the AEs capture the detail of what has occurred within this pathogen-host interaction.

| Pathogen genotype | Host genotype | Term ID | Term name | Evidence code | Conditions | Figure | Annotation extension |
| --- | --- | --- | --- | --- | --- | --- | --- |
| +Sn15Tox1A1(transformant, no endogenous copy)[Not assayed]<br>*P. nodorum* (Sn79-1087) | Snn1-ems237(unknown)[Not assayed]<br>*T. aestivum* (cv. Chinese Spring) | PHIPO:0000481 | absence of pathogen-associated host lesions | Macroscopic observation (qualitative observation) | delivery mechanism: pathogen spore inoculation | Figure 5b | **infects_tissue** leaf , **compared_to_control** Tox1-(no endogenous copy)[Not assayed] Parastagonospora nodorum (Sn79-1087) / Snn1+[WT level] Triticum aestivum (Chinese Spring) , **inverse_gene_for_gene** incompatible interaction, gain of functional pathogen necrotrophic effector, functional host susceptibility locus compromised |

### SECTION 3 Annotation Extensions for curating single species phenotypes (pathogen phenotypes or host phenotypes)

#### AE summary table

SECTION 3: Annotation Extensions for curating single species phenotypes (pathogen phenotypes or host phenotypes) AE summary table
| AE name | Cardinality | Available terms |
| --- | --- | --- |
| affected proteins | 2 | UniProtKB accession number |
| assayed RNA | 0, 1 | UniProtKB accession number |
| assayed protein | 0, 1 | UniProtKB accession number |
| penetrance | 0, 1 | qualitative terms ('high', 'medium', 'low', or 'complete') or a quantitative value (a percentage) |
| severity | 0, 1 | 'high', 'medium', 'low', 'variable severity' |
| observed in organ | 0, 1 | BRENDA Tissue Ontology term |

**Example of an in vitro pathogen phenotype (corresponds to footnote 3 in Table 2**)

Example publication: A conserved fungal glycosyltransferase facilitates pathogenesis of plants by enabling hyphal growth on solid surfaces. (<u>PMID:29020037</u>)

A training video is available for the curation of this publication at https://youtu.be/44XGoi6Ijqk?t=1738

#### Pathogen phenotype

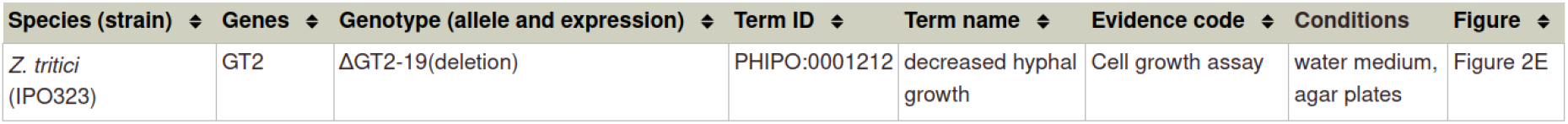

| Species (strain) | Genes | Genotype (allele and expression) | Term ID | Term name | Evidence code | Conditions | Figure |
| --- | --- | --- | --- | --- | --- | --- | --- |
| *Z. tritici* (IPO323) | GT2 | ΔGT2-19(deletion) | PHIPO:0001212 | decreased hyphal growth | Cell growth assay | water medium, agar plates | Figure 2E |

Please note that in this curation example, no AEs were required.

**Example of an in vitro pathogen chemistry phenotype (corresponds to footnote 9 in Table 2**)

Example publication: The T788G mutation in the cyp51C gene confers voriconazole resistance in Aspergillus flavus causing aspergillosis. (<u>PMID:22314539</u>)

#### Pathogen phenotype

| Species (strain) | Genes | Genotype (allele and expression) | Term ID | Term name | Evidence code | Conditions | Figure | Annotation extension |
| --- | --- | --- | --- | --- | --- | --- | --- | --- |
| *A. flavus*<br>(NRRL 3357) | *cyp51c* | *cyp51C*-T788G(aaS240A)[WT level] | PHIPO:0000590 | resistance to voriconazole | Cell growth assay | liquid culture, minimal medium, + voriconazole | Table 3 (footnote d), text page 2602 | has_severity high |
| *A. flavus*<br>(NRRL 3357) | *cyp51c* | *cyp51C*-T161C(aaM54T)[WT level] | PHIPO:0001219 | normal growth on voriconazole | Cell growth assay | liquid culture, minimal medium, + voriconazole | text on page 2602 |  |

**Example of an in vitro host phenotype (corresponds to footnote 7 in Table 2**)

Example publication: Activation of an Arabidopsis resistance protein is specified by the in planta association of its leucine-rich repeat domain with the cognate oomycete effector. (<u>PMID:20601497</u>)

#### Host phenotype

| Species (strain) | Genes | Background | Genotype (allele and expression) | Term ID | Term name | Evidence code | Conditions | Figure | Annotation extension |
| --- | --- | --- | --- | --- | --- | --- | --- | --- | --- |
| *A. thaliana*<br>(ecotype Ws-0) | RPP1 | GFP | RPP1-TIR(266-1221)[Not assayed] | PHIPO:0000467 | presence of effector-independent host hypersensitive response | Macroscopic observation (qualitative observation) | delivery mechanism: agrobacterium, heterologous species tobacco | Figure 7a, c | observed_organ leaf |
| *A. thaliana*<br>(ecotype Ws-0) | RPP1 | GFP | RPP1-TIRNBS(590-1221)[Not assayed] | PHIPO:0001180 | absence of effector-independent host hypersensitive response | Macroscopic observation (qualitative observation) | delivery mechanism: agrobacterium, heterologous species tobacco | Figure 7a | observed_organ leaf |
| *A. thaliana*<br>(ecotype Ws-0) | RPP1 | GFP | RPP1-TIR E158A(266-1221, E158A)[Not assayed] | PHIPO:0001180 | absence of effector-independent host hypersensitive response | Macroscopic observation (qualitative observation) | delivery mechanism: agrobacterium, heterologous species tobacco | Figure 7c | observed_organ leaf |

## Appendix 2 Worked example of a curation session

This document provides a worked example of the curation process in PHI-Canto for the publication by King et al. (2017), *A conserved fungal glycosyltransferase facilitates pathogenesis of plants by enabling hyphal growth on solid surfaces* (<u>PMID:29020037</u>).

The research study confirms the hypothesis that the GT2 gene is required for the fungal pathogens *Zymoseptoria tritici* and *Fusarium graminearum* to cause disease on wheat (*Triticum aestivum*). The curation session in PHI-Canto captures this conclusion by annotating a pathogen–host interaction between *Z. tritici* and *T. aestivum* to show that deletion of the GT2 gene causes loss of pathogenicity in the pathogen, and an absence of pathogen-associated lesions in the host. The wild type interaction between *Z. tritici* and *T. aestivum* is annotated to indicate the presence of disease (and lesions), and a corresponding pathogen–host interaction between *F. graminearum* and *T. aestivum* is annotated to show that deleting GT2 again causes a loss of pathogenicity and the absence of pathogen-associated lesions in the host.

The example starts with the entry of the publication into PHI-Canto (https://canto.phi-base.org/) and ends with the submission of the curation session for review by curators at PHI-base. The information curated from this publication is available on the new gene centric PHI-base 5 website (http://phi5.phi-base.org, search for PHIG:308 and PHIG:307).

### Entering the publication

The PHI-Canto homepage provides a text field where publications can be entered by providing their PubMed ID (PMID). The PMID in this case is 29020037.

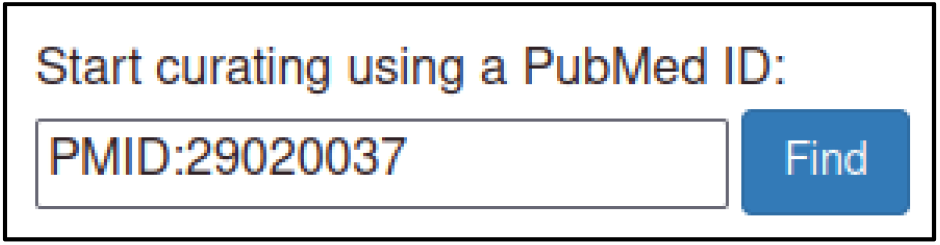

PHI-Canto will automatically retrieve details of the publication from PubMed so that the curator can confirm that they have entered the correct PMID.

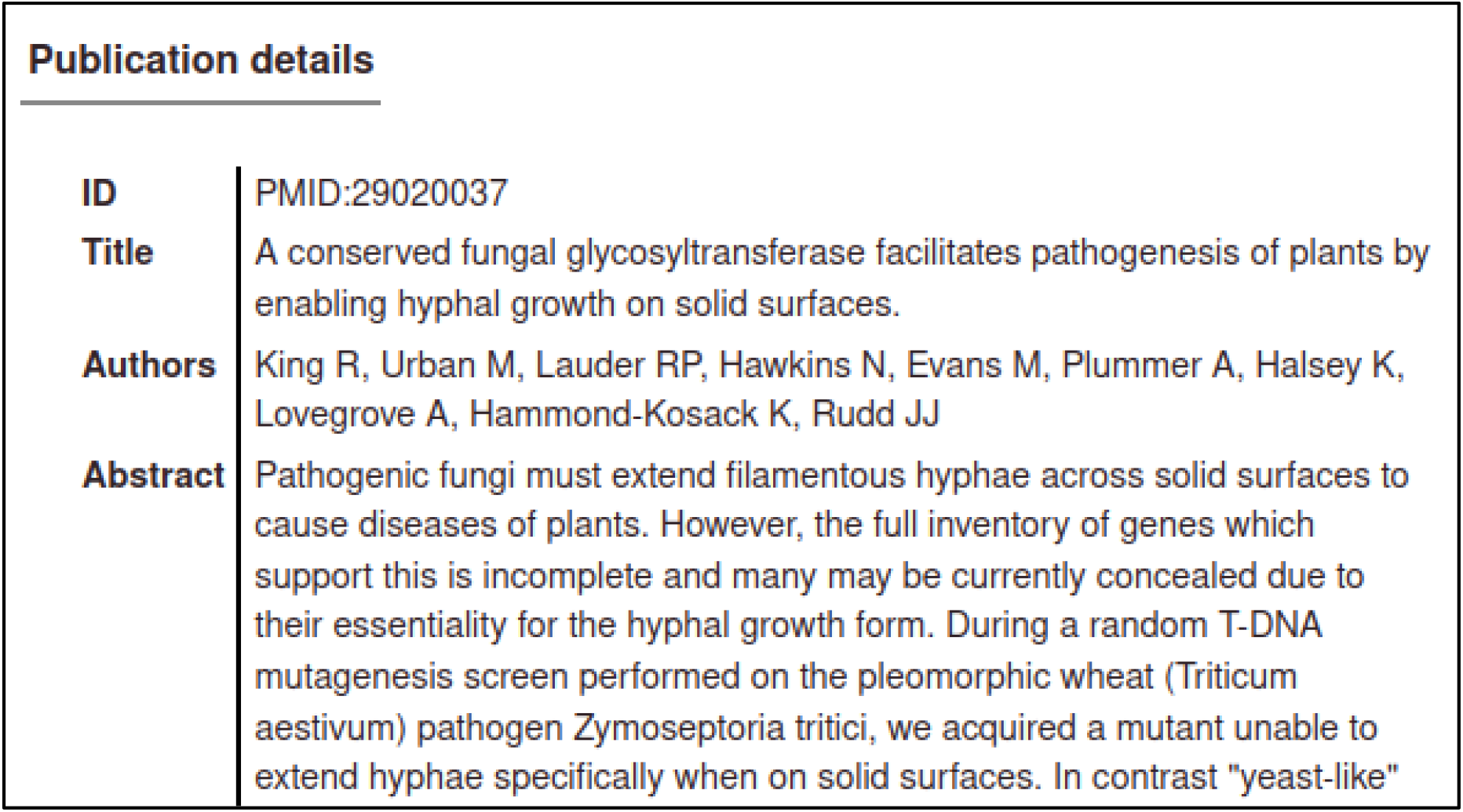

After accepting the publication, the curator is prompted for their name, email address, and (optionally) an ORCID ID, which are used to attribute the curation to the curator, and to contact the curator in case of problems with the curation session.

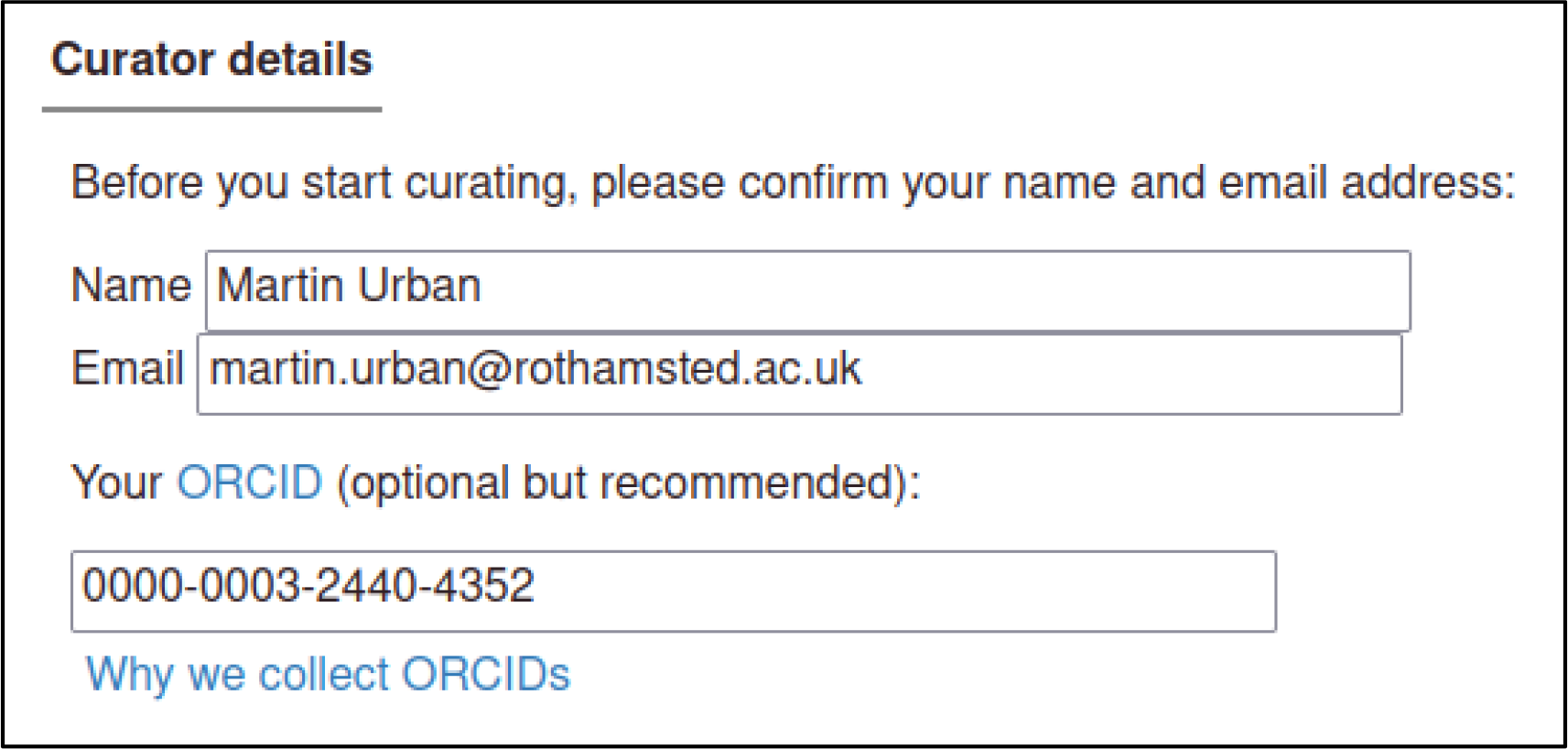

### Specifying genes and species

The gene is the most basic unit of annotation in PHI-Canto: every other biological feature that can be annotated involves a gene, so genes are entered first. PHI-Canto uses accession numbers from the UniProt Knowledgebase (UniProtKB) to uniquely identify proteins for the genes of interest in the curated publication.

The UniProtKB accession numbers for the publication are shown below.

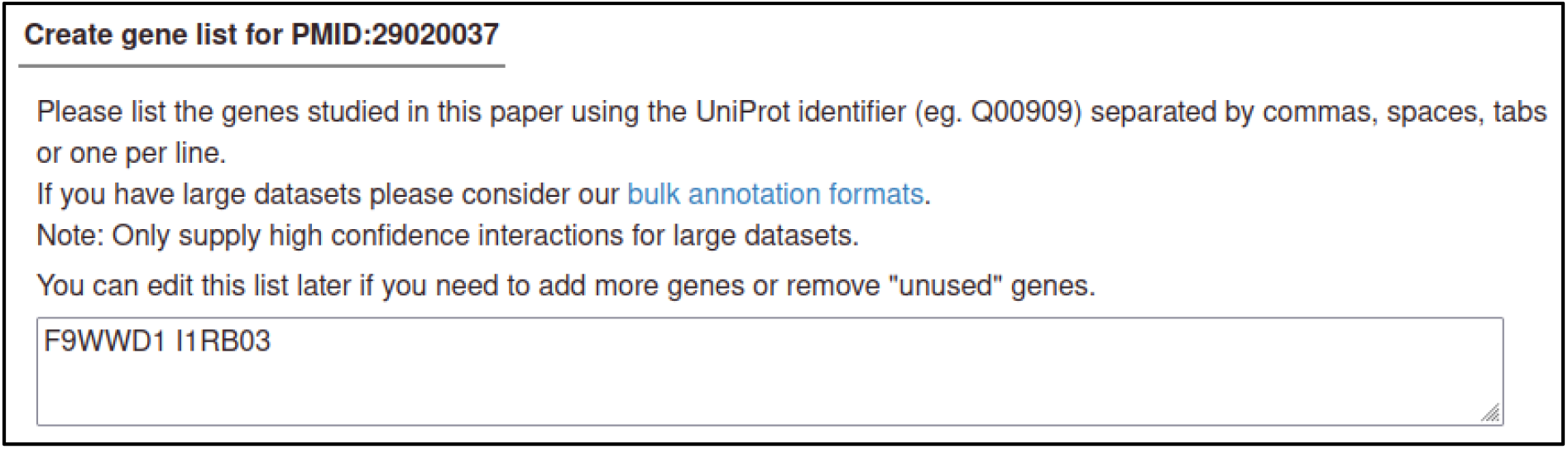

Since this publication describes a wild type host species (*T. aestivum*) with no specified genes of interest, the curator must add the host to the session by entering its NCBI Taxonomy ID in a separate field.

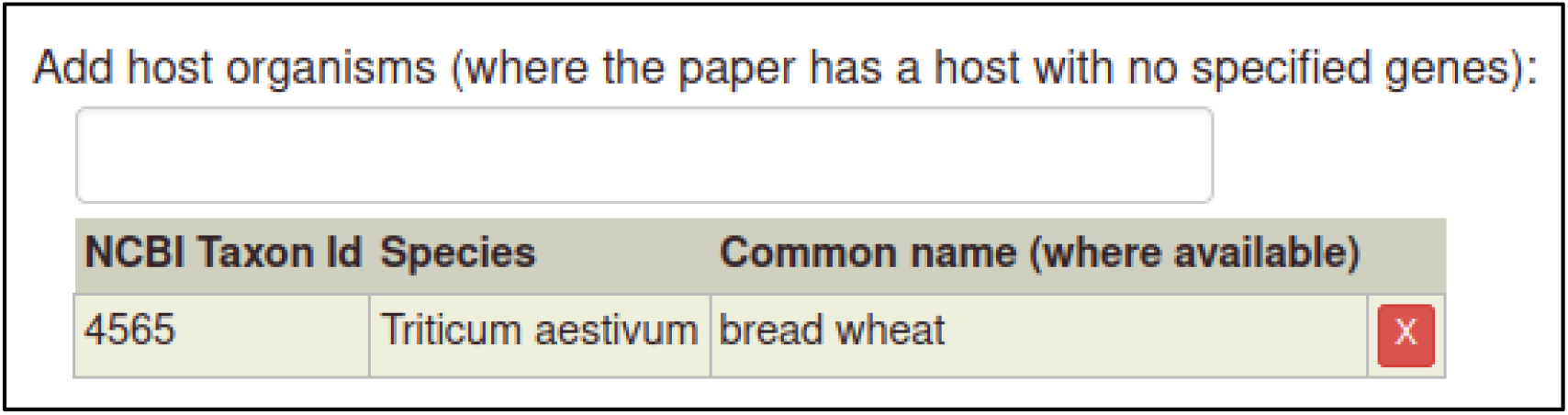

PHI-Canto automatically retrieves details of the proteins from UniProtKB, including the gene name, gene product, and taxonomy (e.g., the species name).

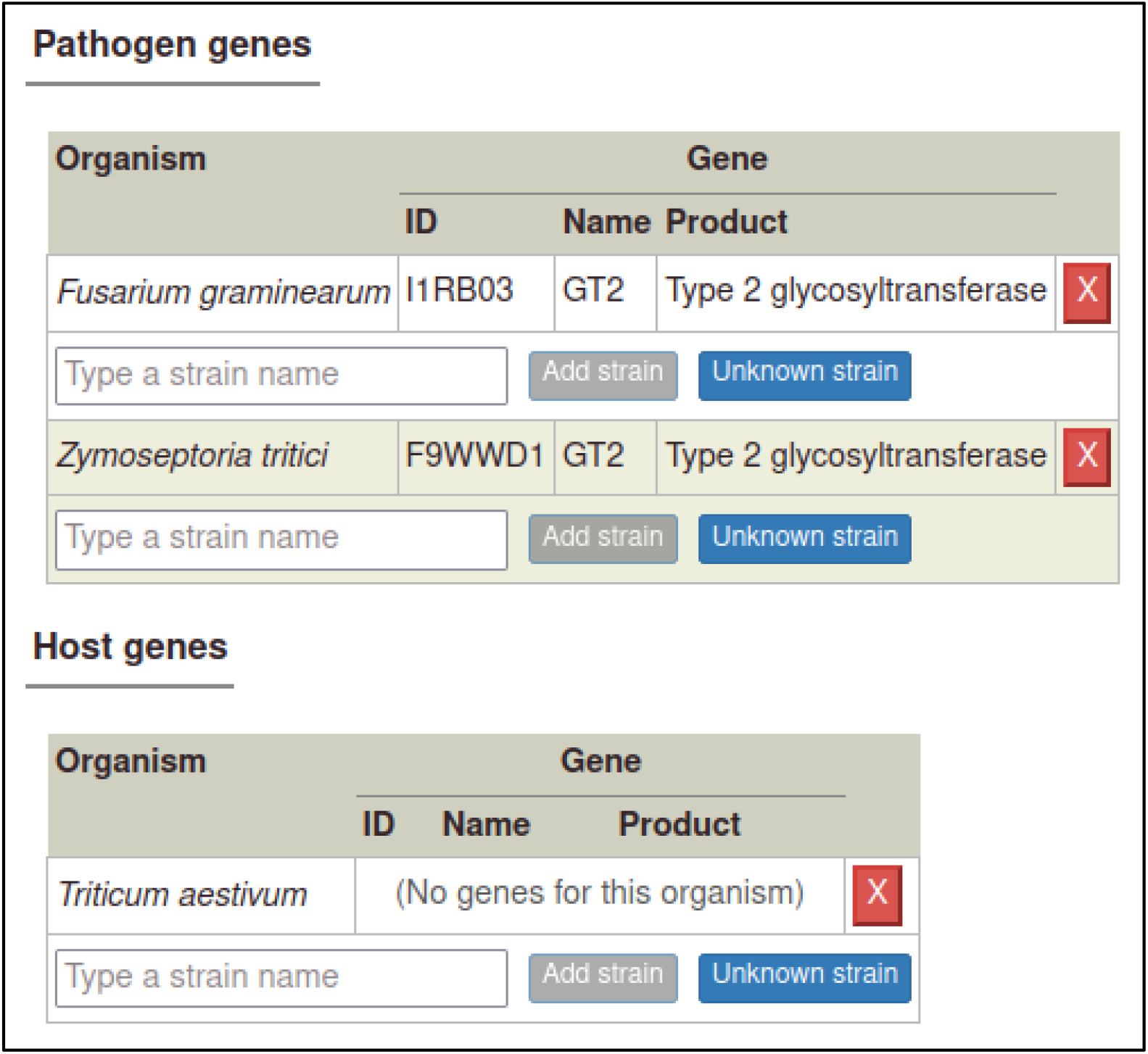

### Specifying strains

The curator must enter the strains for each organism studied in the publication or must specify when the strain was not known (or not specified in the publication). PHI-Canto provides a pre-populated list of strains for many species that the curator can select from, though they also have the option to specify a strain not in the list as free text.

In this publication, the pathogen strains are PH-1 for *F. graminearum* and IPO323 for *Z. tritici*. Two cultivars of *T. aestivum* were used: cv. Bobwhite and cv. Riband.

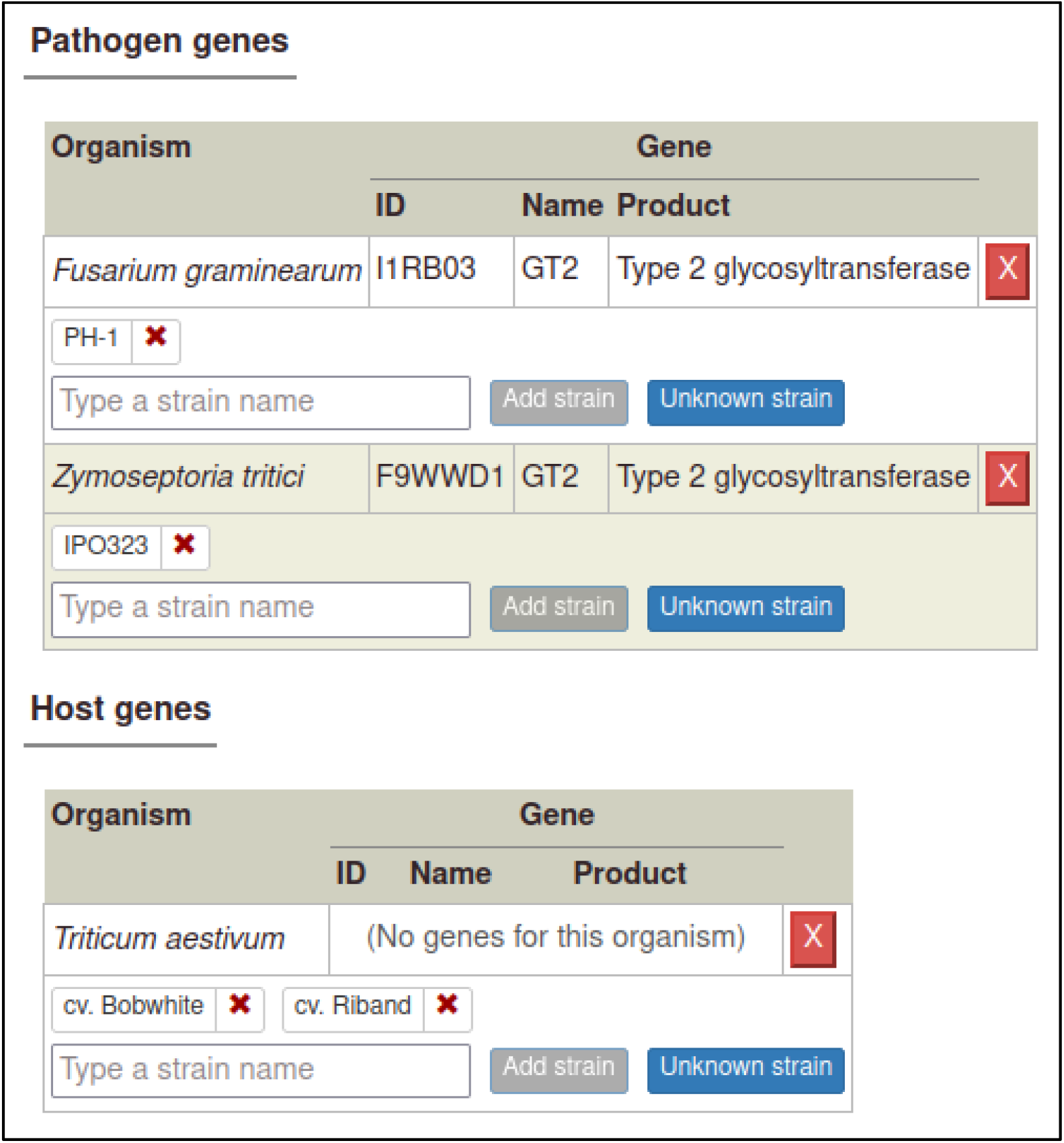

### Creating alleles and genotypes

In order to show that deleting GT2 in the pathogen causes a loss of pathogenicity, the curator must annotate the interaction between the mutant pathogen and its host with a phenotype, meaning the interaction must be added to the curation session. In PHI-Canto, interactions are represented as *metagenotypes*, which are the combined genotypes of the pathogen and host species.

Before the curator can create a metagenotype, they must first create a genotype. Genotypes are composed from alleles (except in the case of wild type host genotypes with no specified genes, as described later), and metagenotypes are composed from genotypes. So, the curator must first create an allele from a gene, then a genotype from an allele, then a metagenotype from two genotypes.

The curator starts from the Pathogen genotype management page, following a link from the Curation summary page.

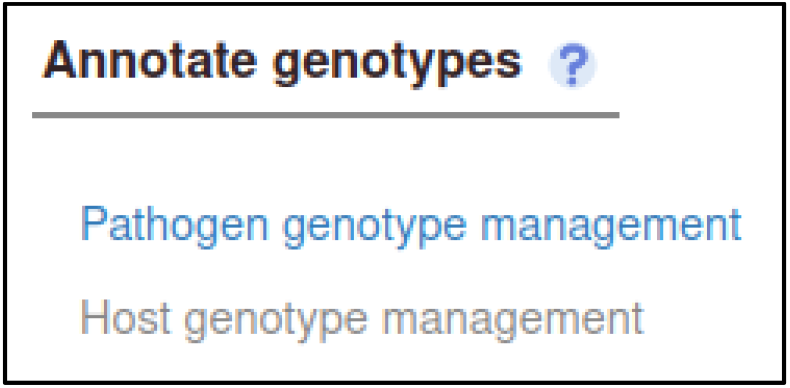

The curator then selects a pathogen species (*Z. tritici*) from a drop-down menu.

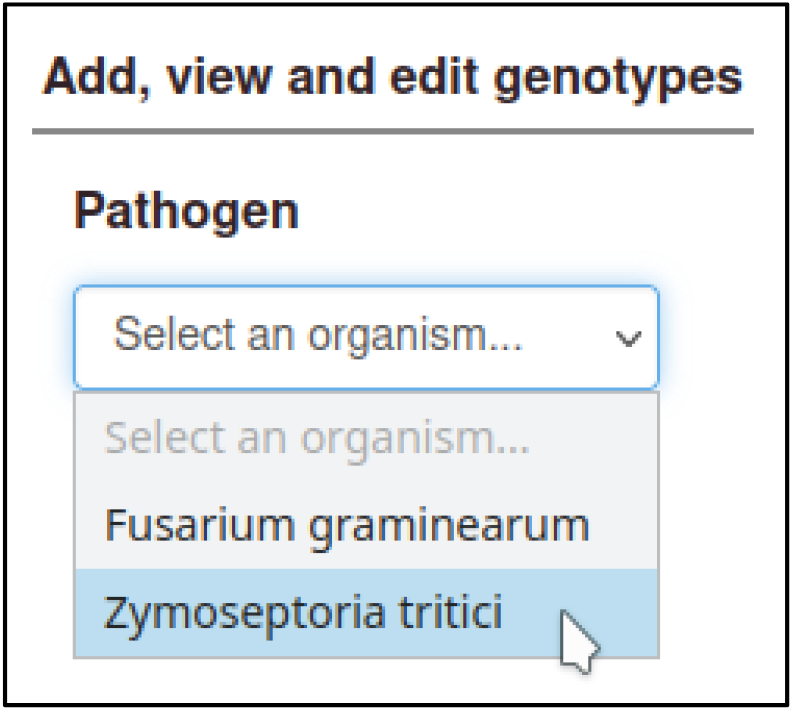

Selecting a pathogen species shows a list of genes for the species, with buttons to create types of alleles. Here, the curator selects ‘Deletion’ for a deletion allele.

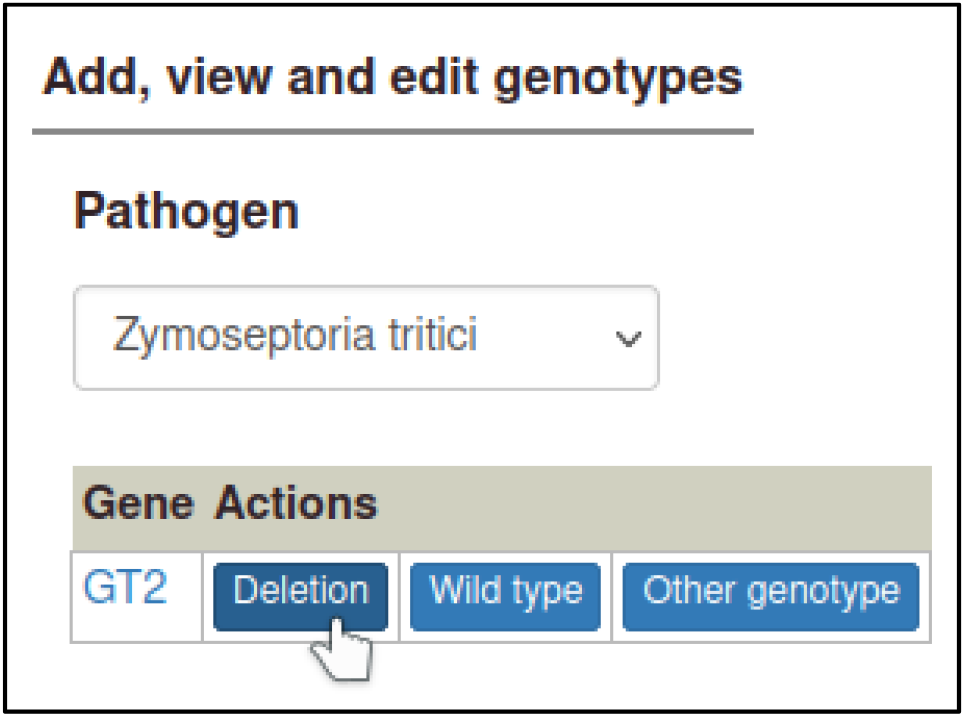

The curator is prompted for the strain the deletion occurred in.

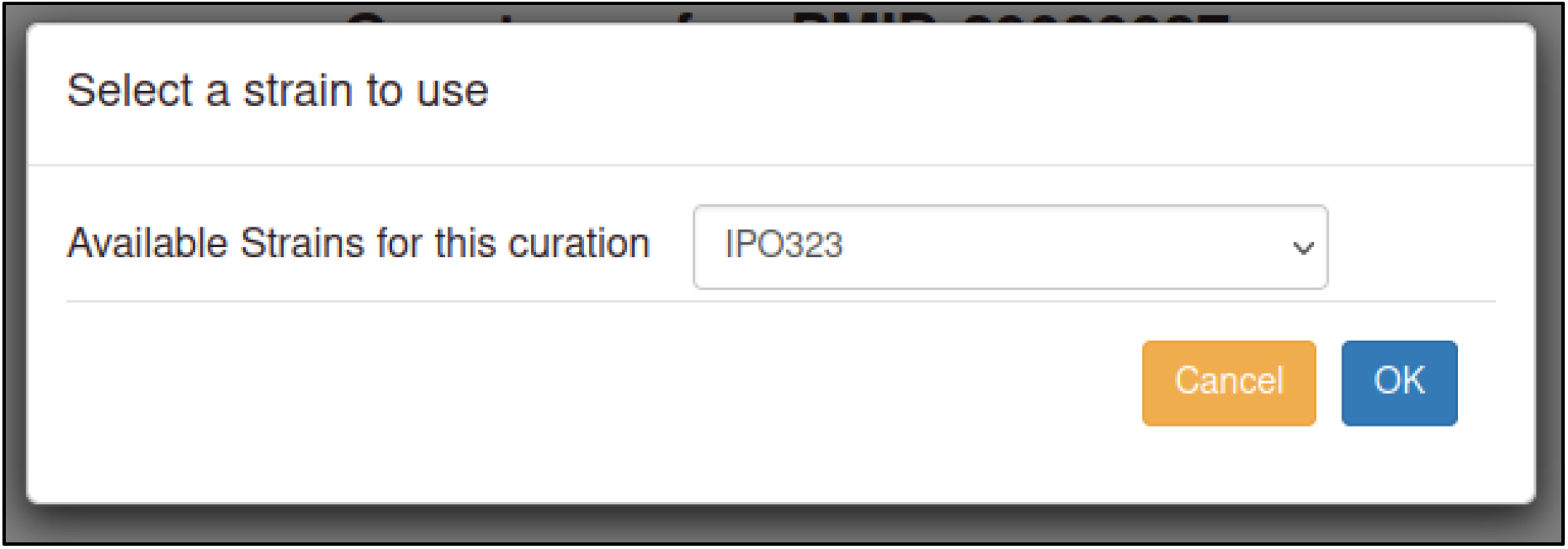

After selecting this, PHI-Canto creates a genotype containing a single allele, with the allele name automatically generated from the gene name followed by a delta symbol.

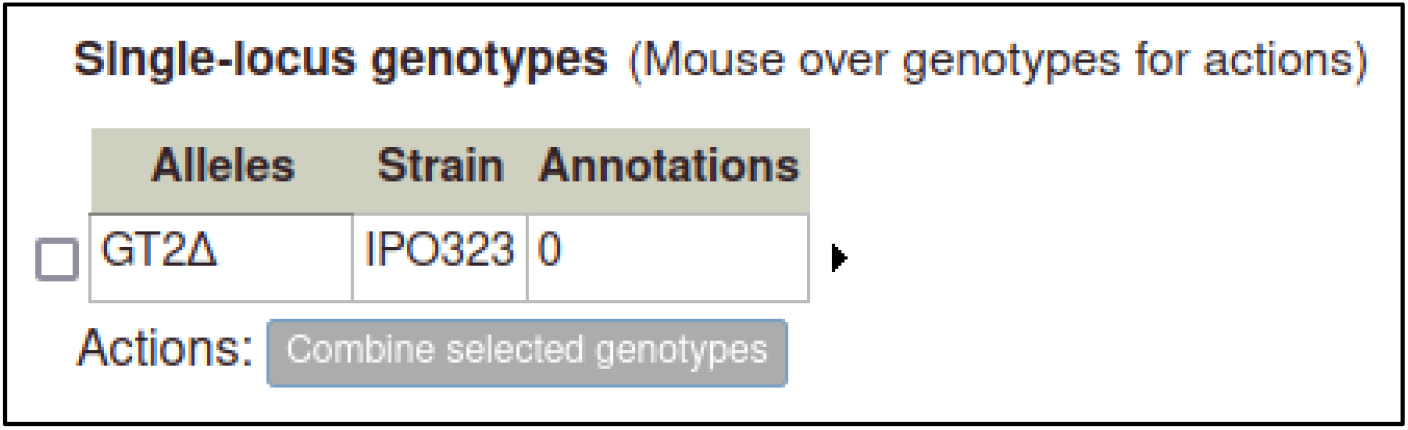

The curator will also need to prepare a wild type genotype for the pathogen GT2 gene, which can be added to the control metagenotype so that any changes in the phenotype (between the wild type pathogen and the altered pathogen inoculated onto the host) can be properly annotated. This first requires making a wild type allele for GT2, using the ‘Wild type’ allele type.

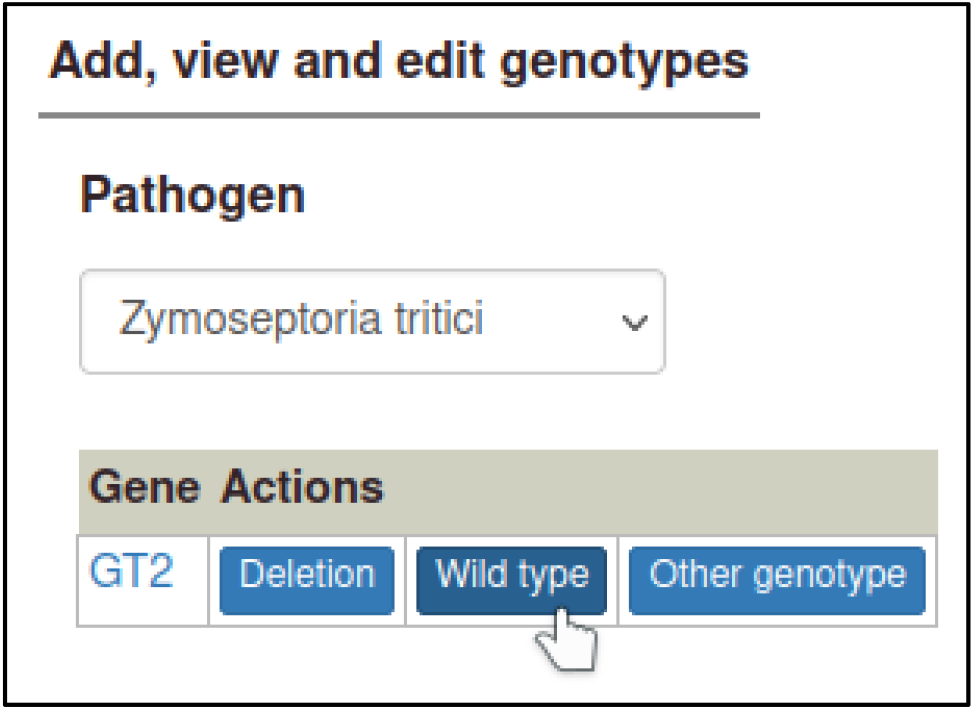

Wild type alleles require the gene expression level to be specified. In this case, there was no change in expression level, so the curator selects ‘Wild type product level’. PHI-Canto automatically creates an allele name by appending a plus symbol to the gene name.

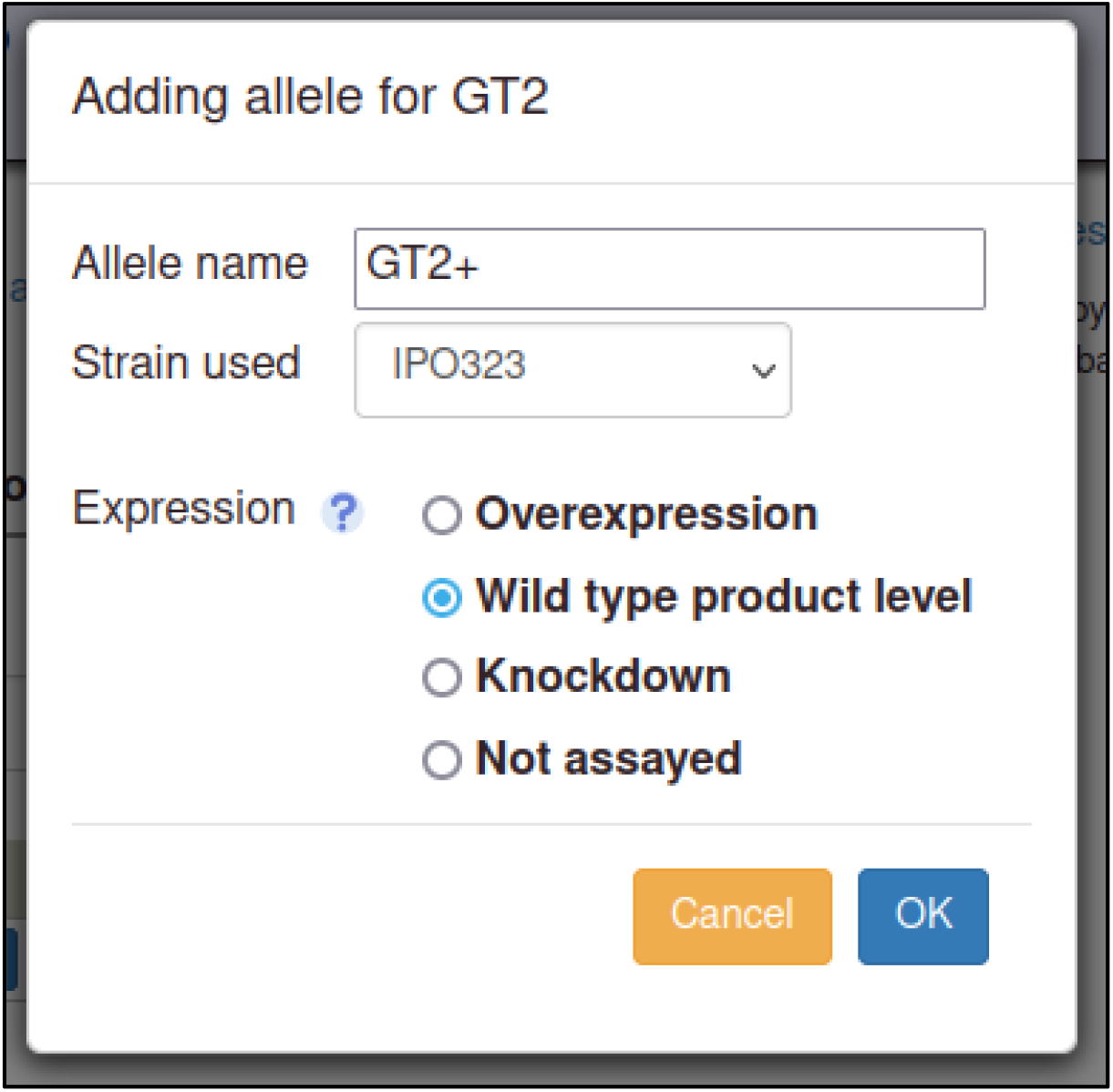

As genotypes are created, they are added to a table of genotypes on their respective genotype management page (Pathogen genotype management for pathogens, Host genotype management for hosts).

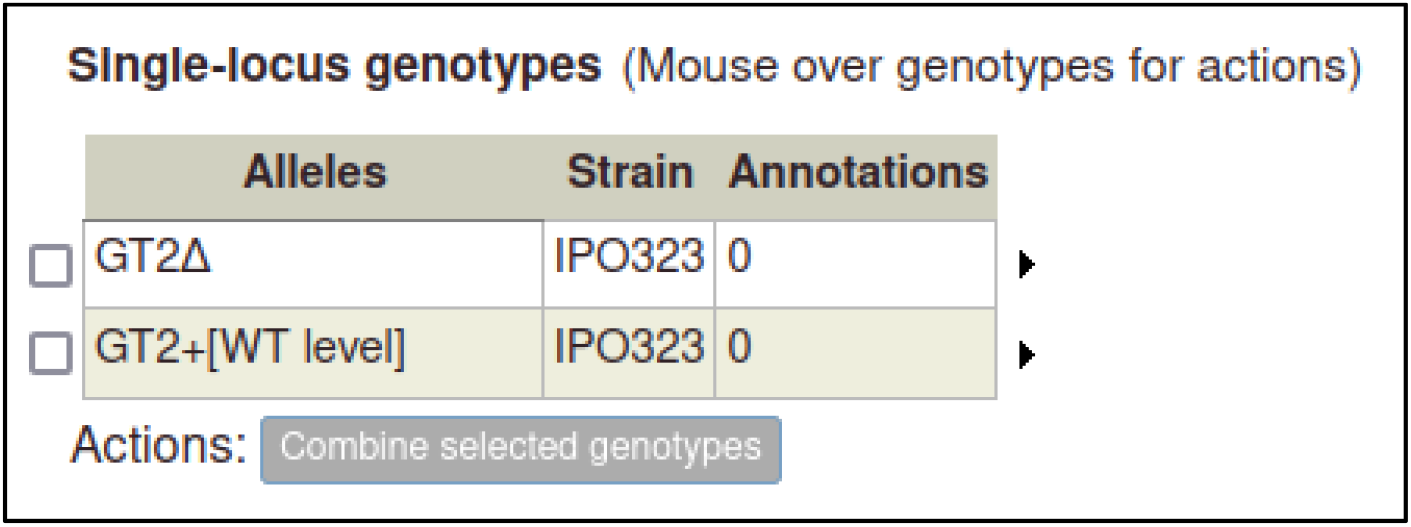

The curator can repeat the process above to create pathogen genotypes for *F. graminearum*.

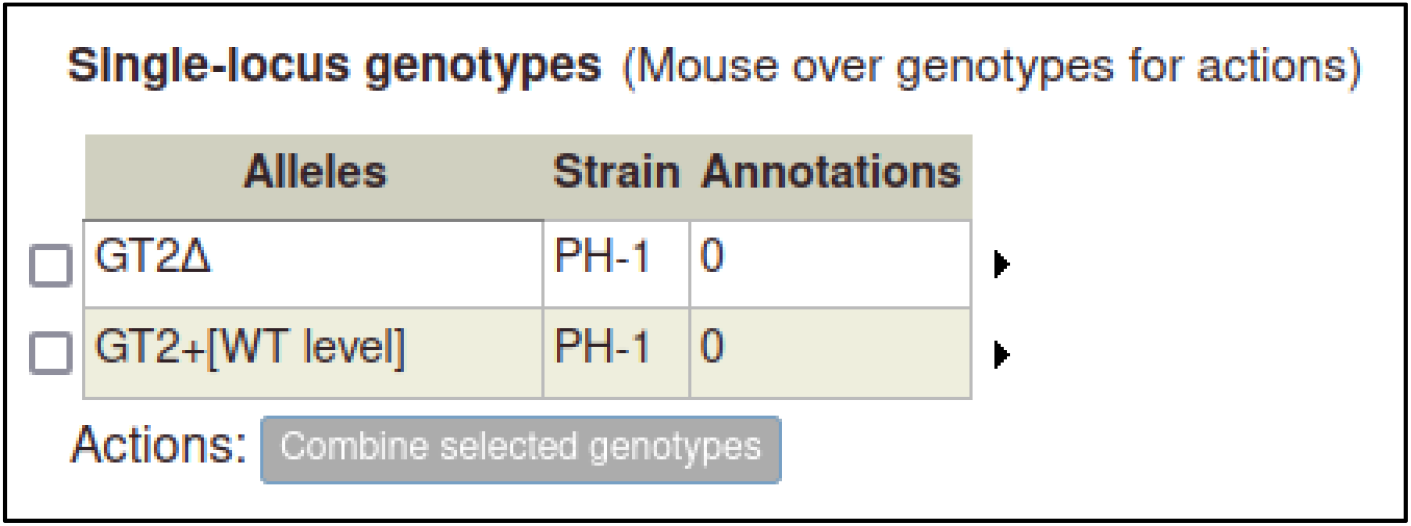

### Creating metagenotypes for pathogen–host interactions

Metagenotypes are created using the Metagenotype management page, where genotypes previously added to the curation session can be combined into a metagenotype. The curator can reach this page from the Curation Summary page, or from either the pathogen or host genotype management page.

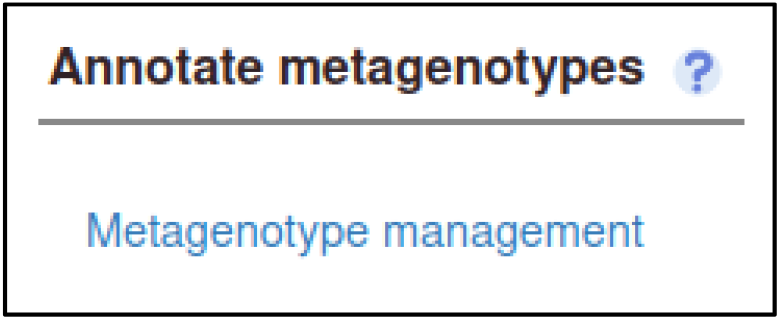

The curator starts by selecting a pathogen species from a drop-down menu.

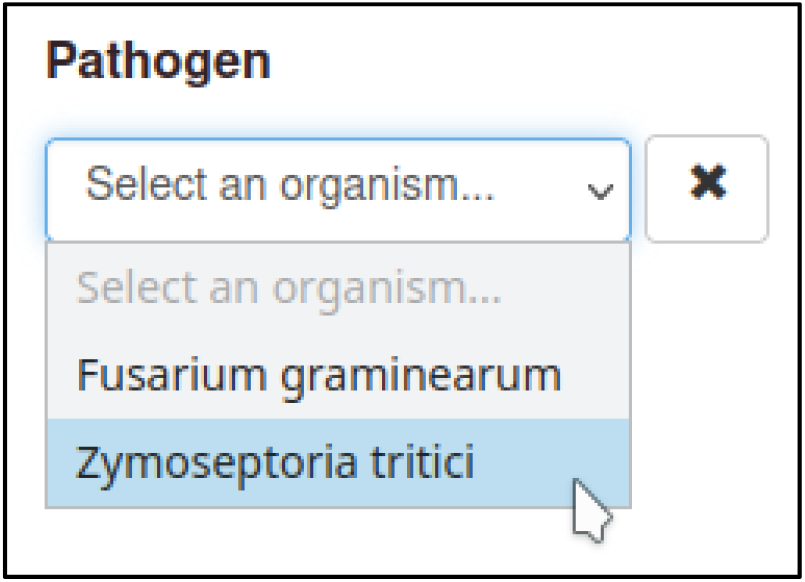

Then the curator selects a genotype from the table of pathogen genotypes.

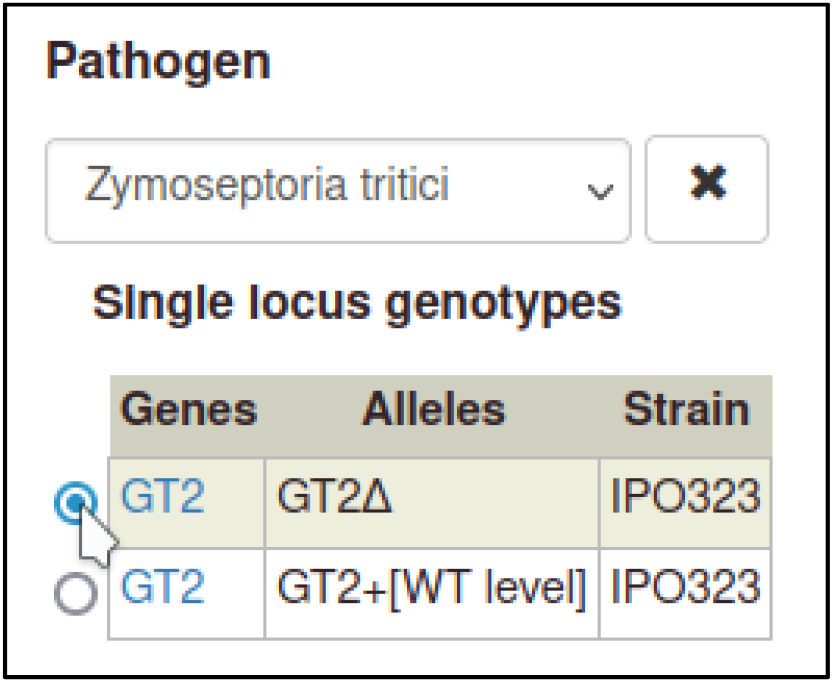

Then the curator selects a host genotype. For wild type hosts, PHI-Canto provides a shortcut where a strain can be selected without needing to create an allele as part of the genotype.

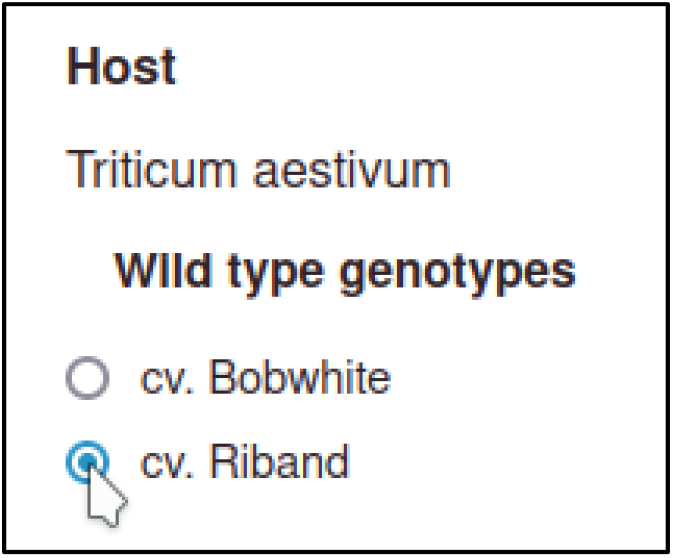

The curator selects ‘Make metagenotype’ to create the metagenotype for the interaction.

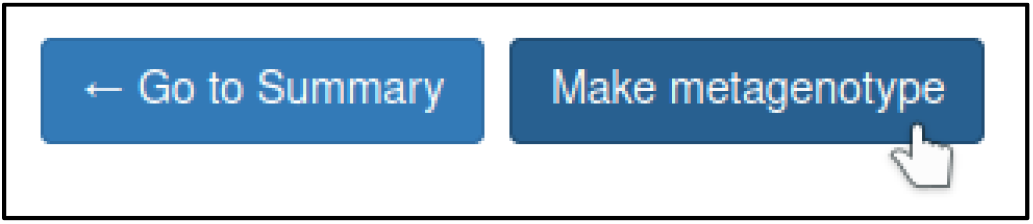

The metagenotype is displayed in a table as a combination of pathogen and host genotype.

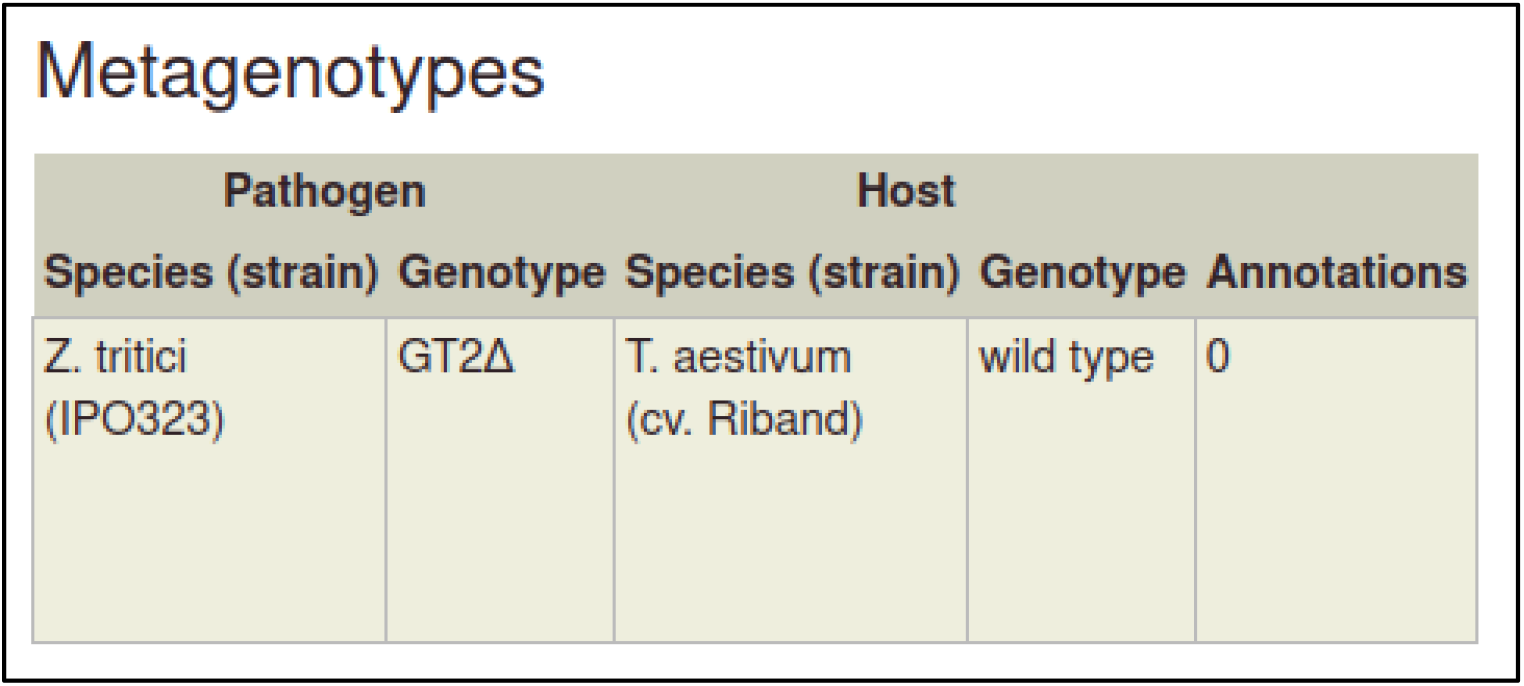

This process can be repeated to create the metagenotype for the wild type interaction between *Z. tritici* and *T. aestivum*. In this case, the pathogen genotype containing the wild type GT2 is selected instead of the deletion allele.

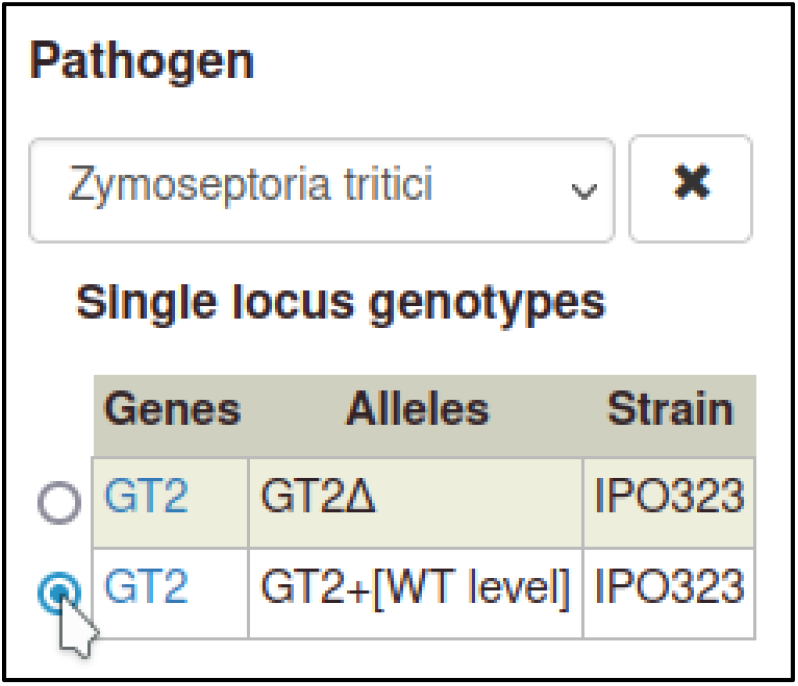

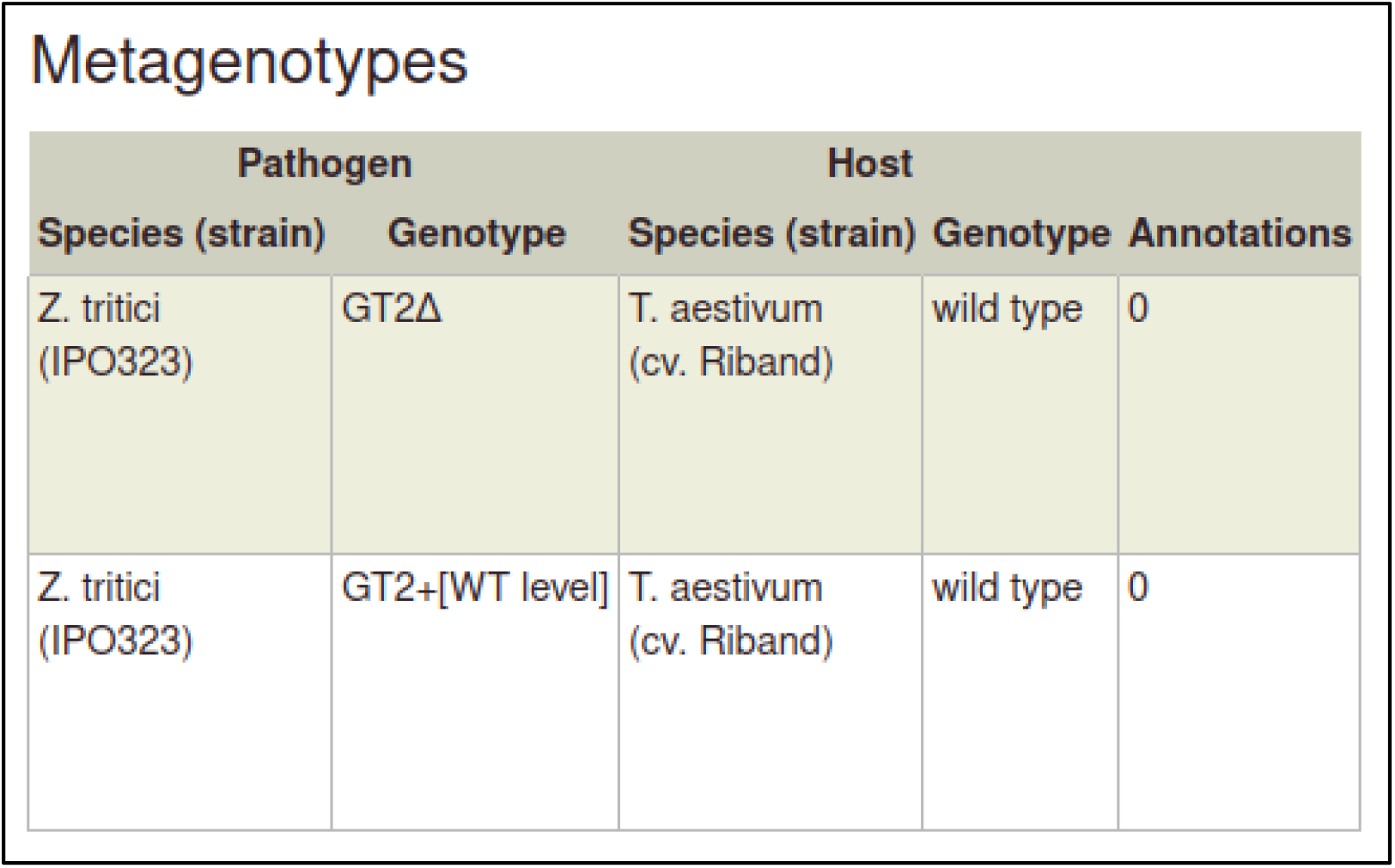

Creating the corresponding metagenotypes for *F. graminearum* and *T. aestivum* simply requires changing the pathogen species and selecting cv. Bobwhite for the host strain.

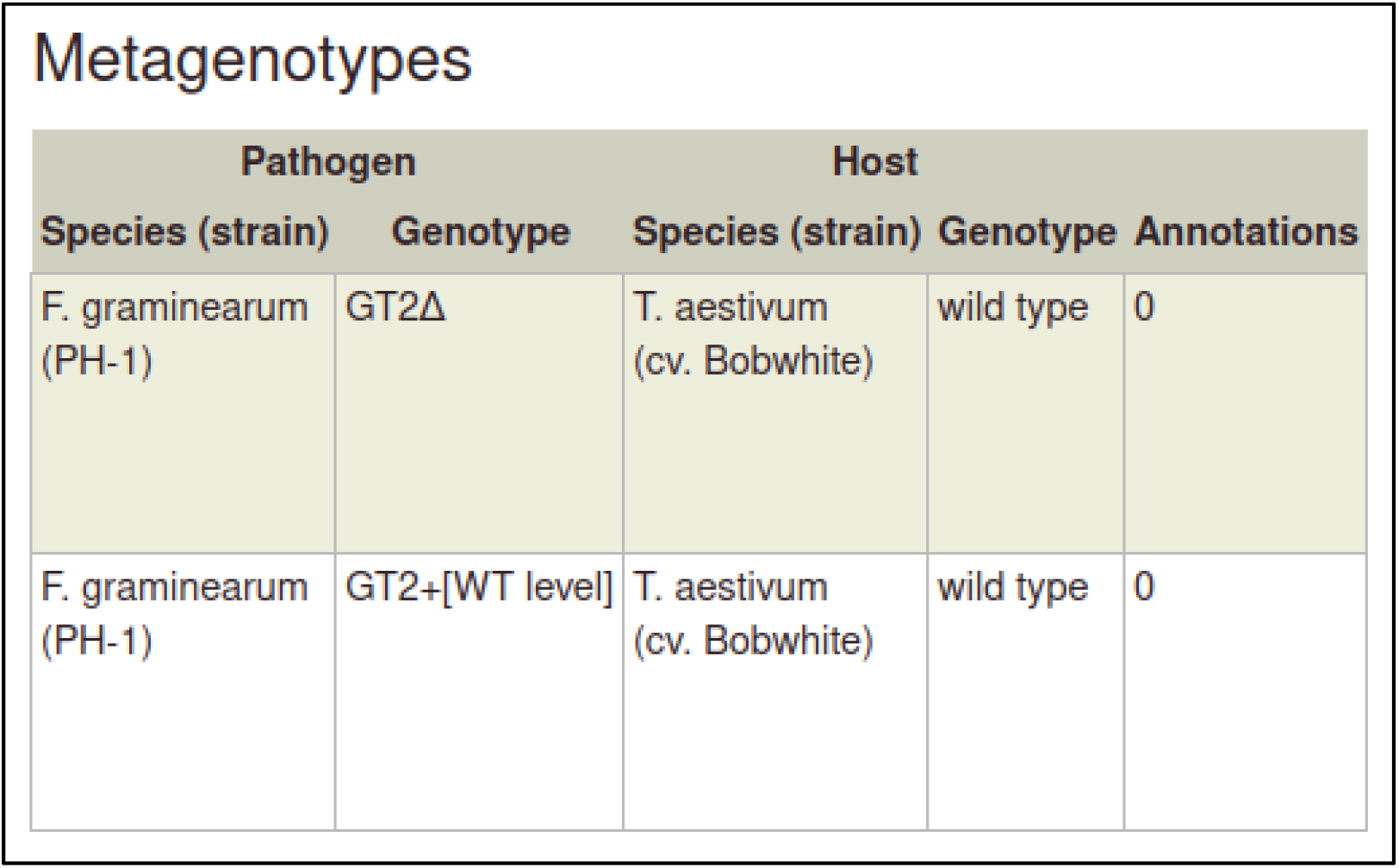

### Annotating pathogen–host interactions with phenotypes

Metagenotypes can be annotated with phenotypes by selecting the ‘Annotate pathogen-host interaction phenotype’ action.

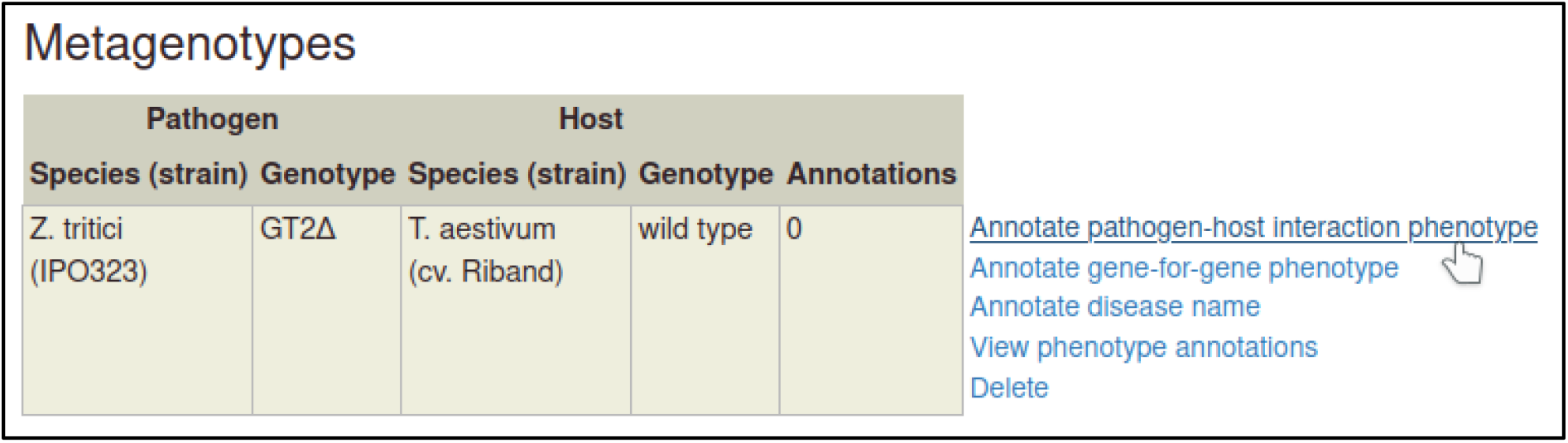

#### Phenotype and evidence

The first step is to select a term from a controlled vocabulary that describes the phenotype of the interaction. PHI-Canto uses terms from the Pathogen–Host Interaction Phenotype Ontology (PHIPO) for this purpose. The primary observed phenotype in this case is the *absence of pathogen-associated host lesions* (PHIPO:0000481).

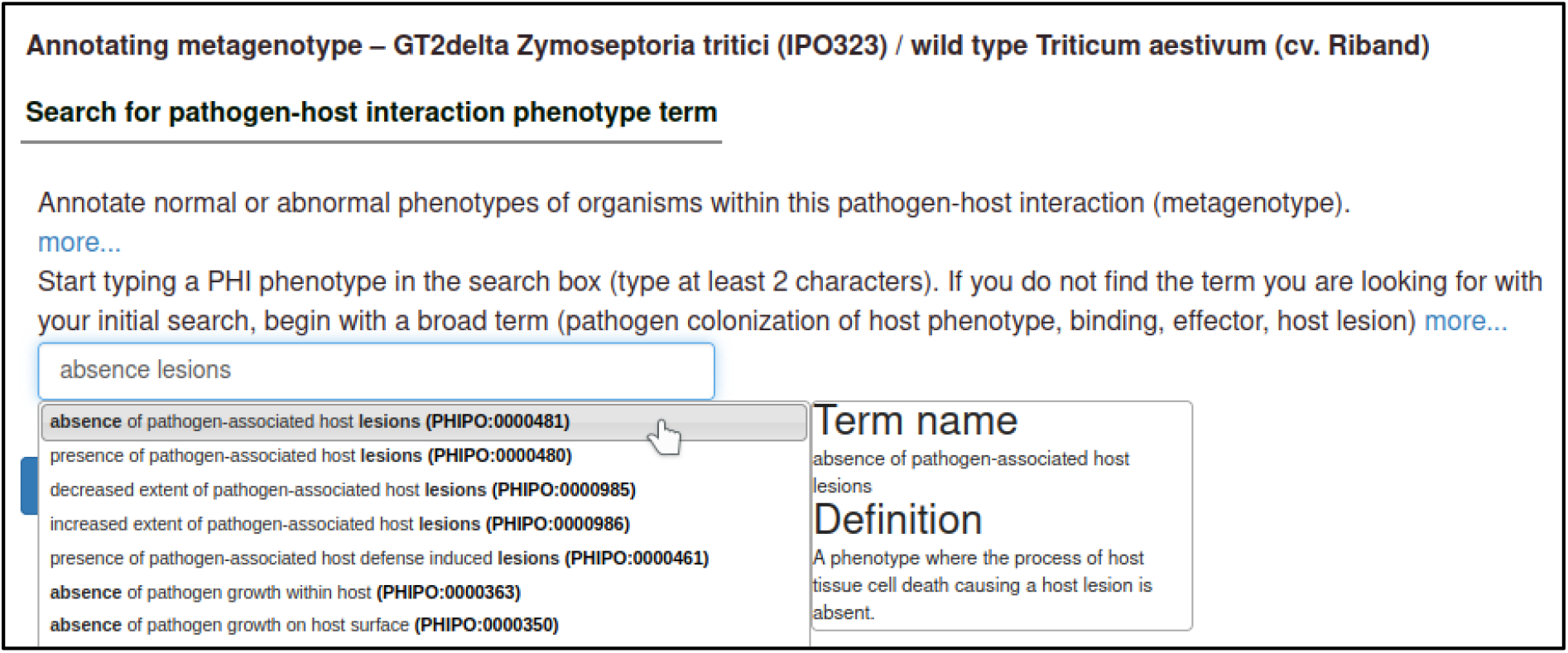

Upon selecting the term, the curator is shown a description of the term and its synonyms to help confirm that their chosen term is appropriate.

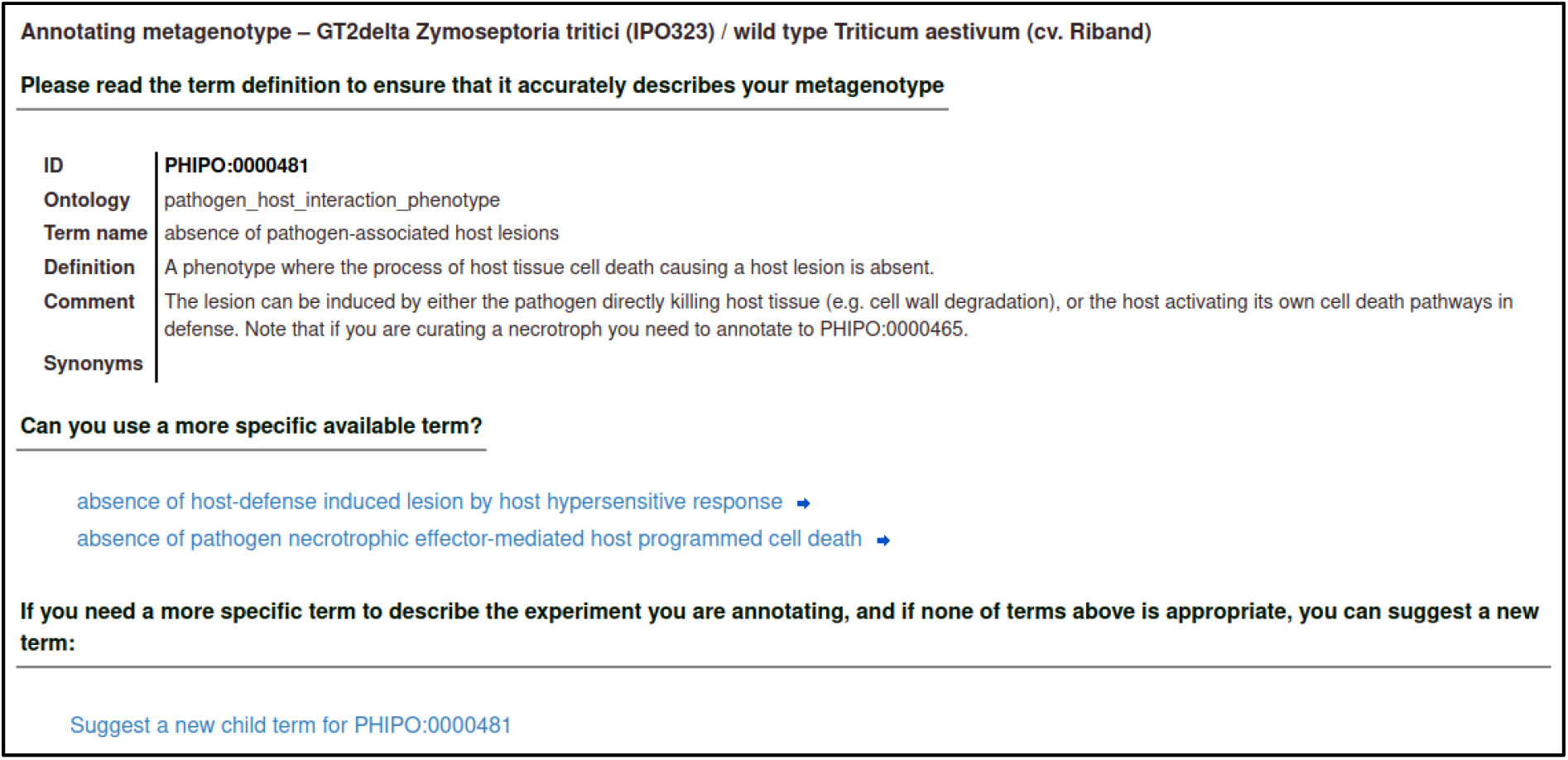

The curator must select an evidence code for the observation of the phenotype. In this case, the phenotype was observed macroscopically, and measured qualitatively.

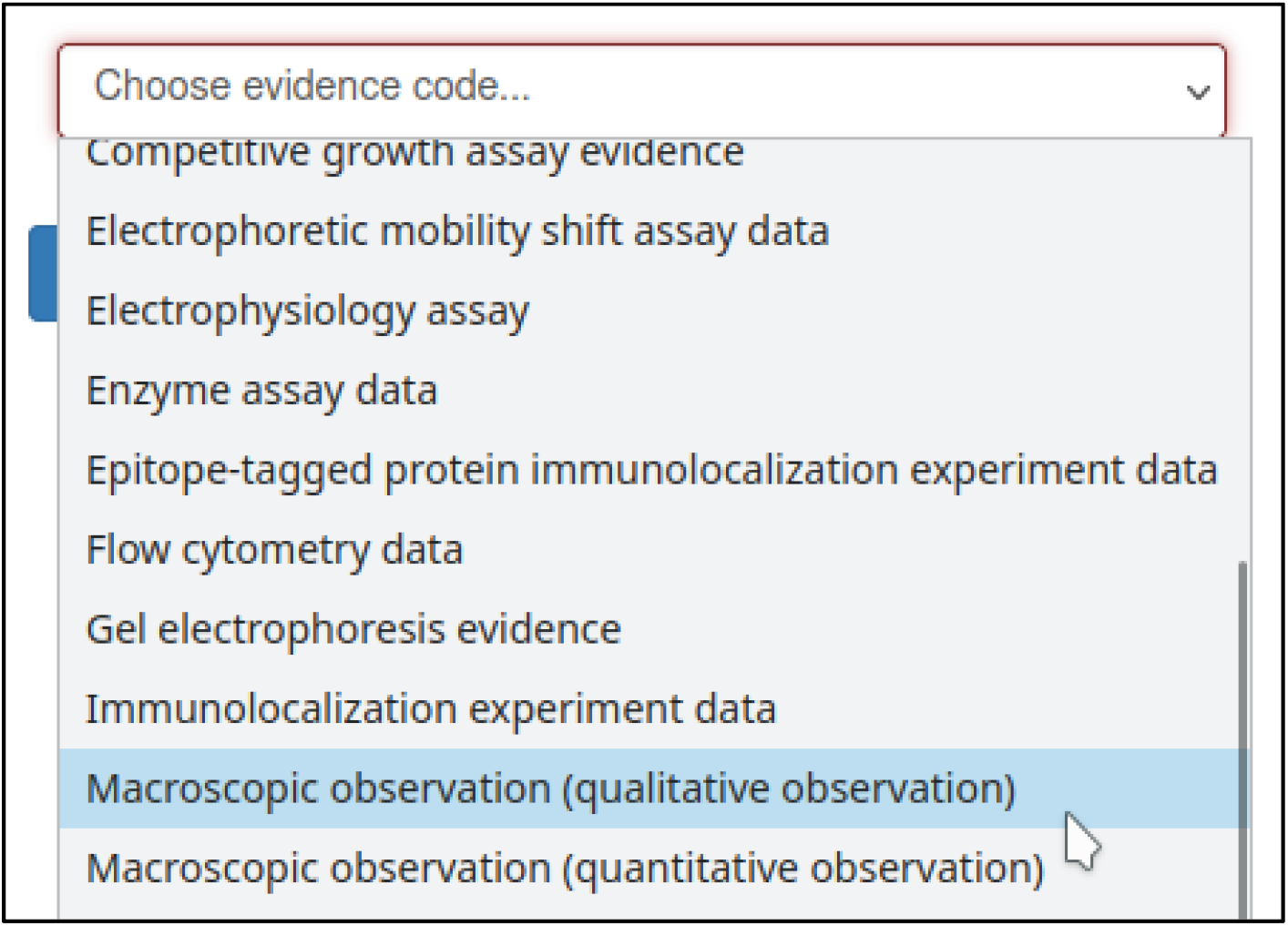

The curator may also specify experimental conditions for the experiment – such as the growth medium, or days elapsed after inoculation of the host. This annotation specifies that the assay was performed 14 days after inoculation with the *Z. tritici* GT2 deletion mutant.

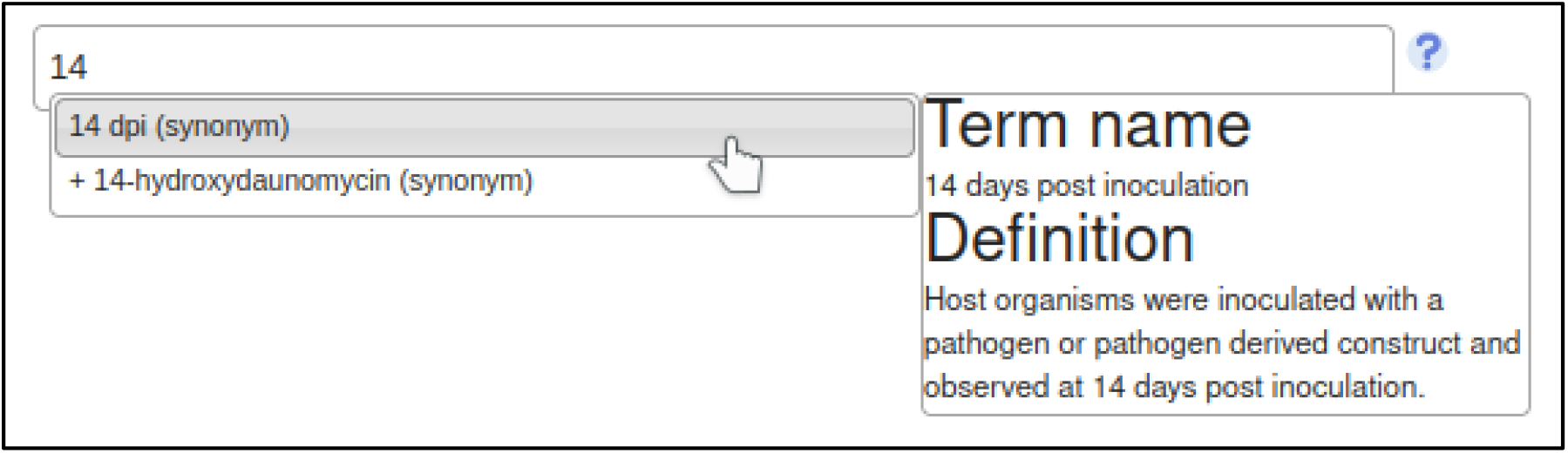

#### Annotation extensions

PHI-Canto uses annotation extensions to provide additional information about the conditions and outcome of the pathogen–host interaction. Of particular note are the host tissue infected, the changes to the infective ability of the pathogen, the presence (or absence) of disease, and the interaction used as a control for the interaction involving a mutant pathogen.

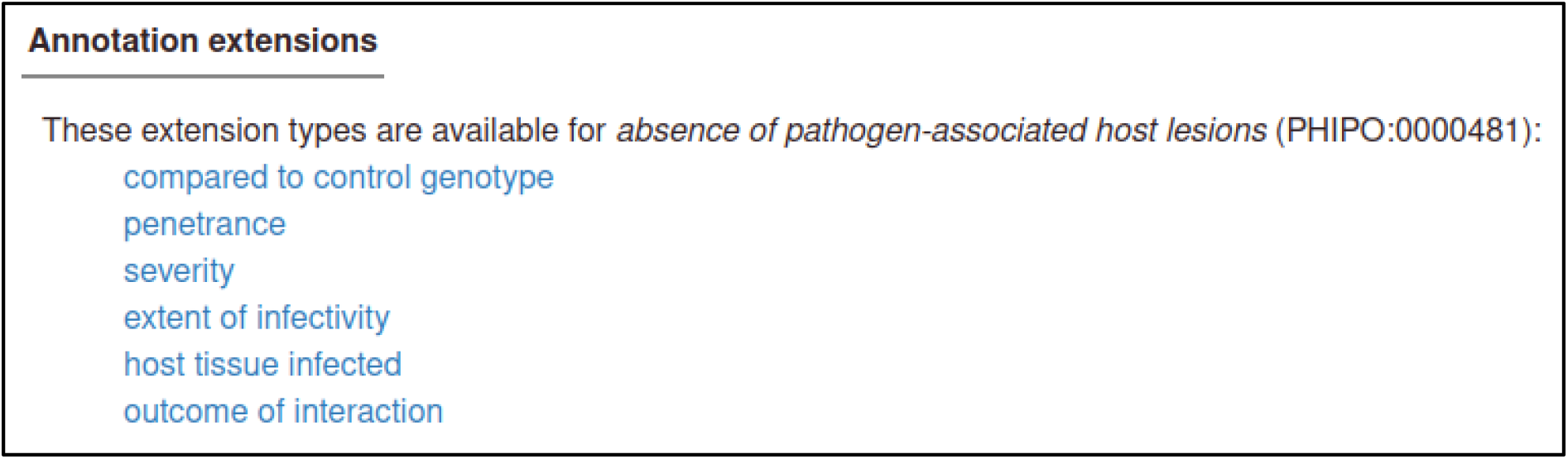

The host tissue that was infected during the interaction is annotated with the ‘host tissue infected’ annotation extension. This extension uses ontology terms from the BRENDA Tissue Ontology (BTO). In this case, the curator specifies that the *leaf* (BTO:0000713) of *T. aestivum* was infected.

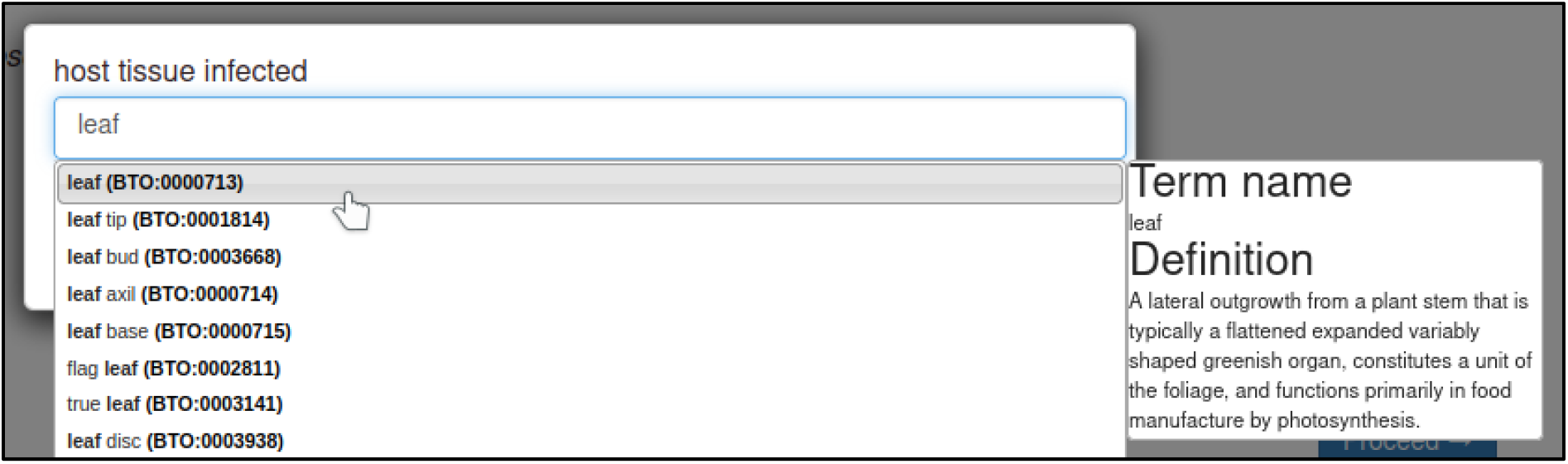

Changes in the infective ability of the pathogen are annotated with the ‘extent of infectivity’ annotation extension. This extension uses a subset of ontology terms from PHIPO. In this case, the curator specifies that the interaction resulted in a *loss of pathogenicity* (PHIPO:0000010).

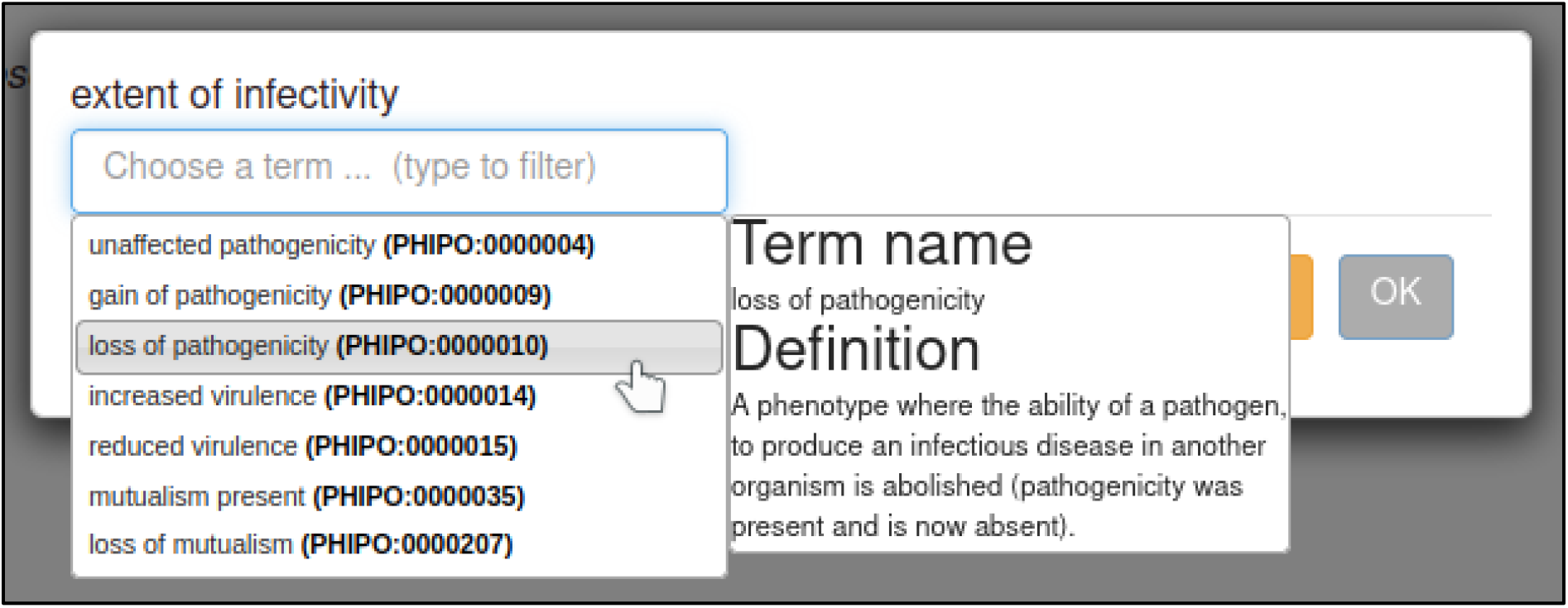

The control interaction (to which the interaction being annotated should be compared) can be annotated with the ‘compared to control genotype’ annotation extension. This annotation allows any metagenotype in the curation session to be designated as a control. In this case, the curator selects the wild type metagenotype that was created earlier.

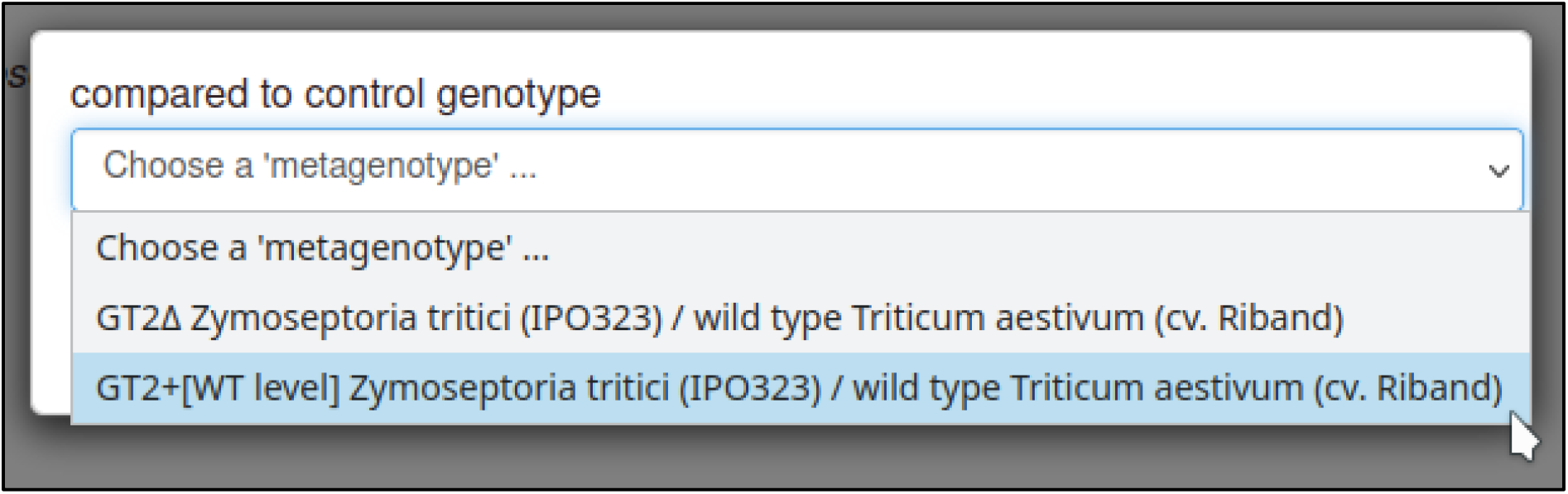

The presence or absence of disease resulting from the interaction can be annotated with the ‘outcome of interaction’ annotation extension. This extension uses a subset of ontology terms from PHIPO. In this case, the curator specifies that no disease was observed as a result of the interaction: *disease absent* (PHIPO:0001199).

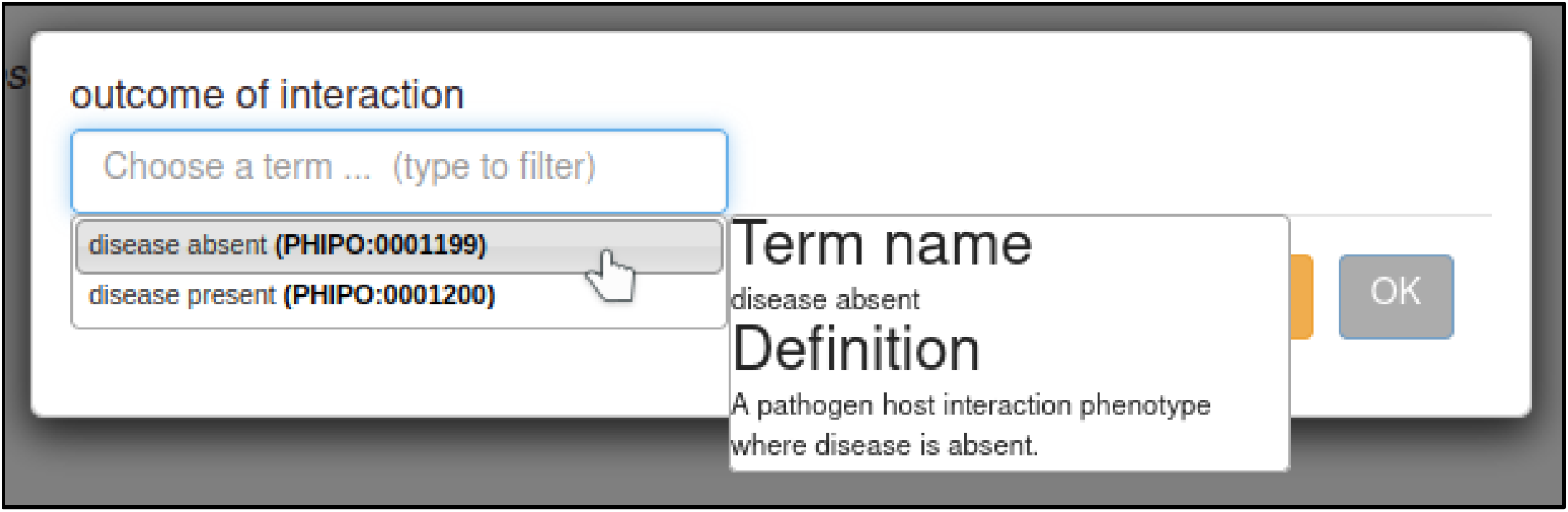

#### Figure numbers and comments

After adding annotation extensions, the curator has the option to provide the figure number from the publication (if any) that illustrates the phenotype. In this case, the figure was Figure 2E.

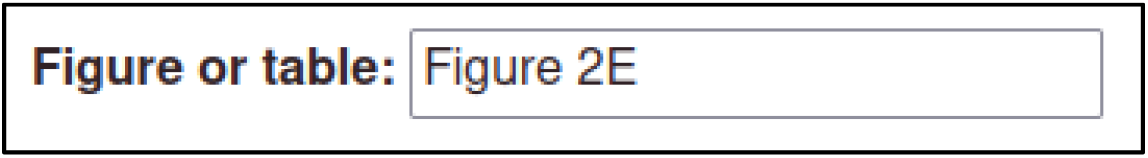

The curator can also provide additional information in a comments field, in case of details that are not appropriate for any other field.

Once the above steps are completed, the phenotype annotation is created.

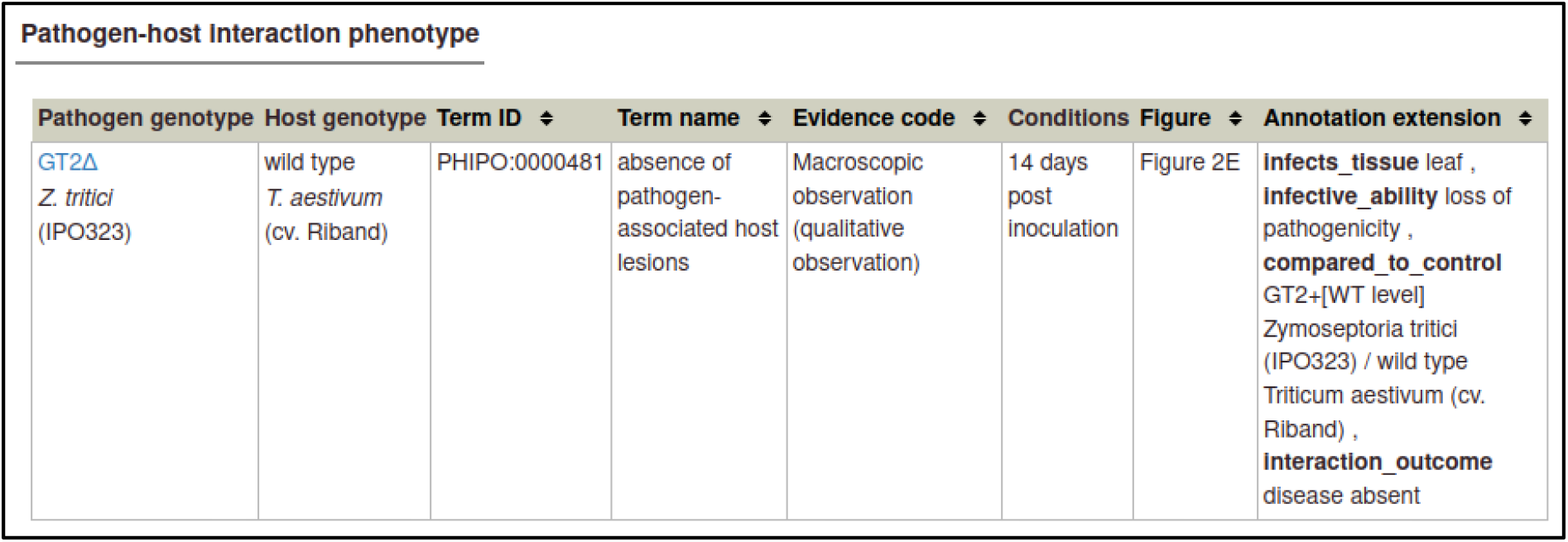

| Pathogen-host interaction phenotype |  |  |  |  |  |  |  |
| --- | --- | --- | --- | --- | --- | --- | --- |
| Pathogen genotype | Host genotype | Term ID | Term name | Evidence code | Conditions | Figure | Annotation extension |
| GT2Δ<br>*Z. tritici*<br>(IPO323) | wild type<br>*T. aestivum*<br>(cv. Riband) | PHIPO:0000481 | absence of pathogen-associated host lesions | Macroscopic observation (qualitative observation) | 14 days post inoculation | Figure 2E | **infects_tissue** leaf , **infective_ability** loss of pathogenicity , **compared_to_control** GT2+[WT level] *Zymoseptoria tritici* (IPO323) / wild type *Triticum aestivum* (cv. Riband) , **interaction_outcome** disease absent |

#### Copying annotations

The above annotation can be used as a template for the interaction between the wild type pathogen and host, since many of the variables are the same. PHI-Canto provides a ‘Copy and edit’ feature that allows curators to use one annotation as a template for creating another.

| Conditions | Figure | Annotation extension |  |
| --- | --- | --- | --- |
| 14 days post inoculation | Figure 2E | **infects_tissue** leaf , **infective_ability** loss of pathogenicity , **compared_to_control** GT2+[WT level] | View metagenotype<br>Edit<br>Copy and edit<br>Delete |

For the wild type interaction, the pathogen genotype is changed to wild type GT2, the phenotype term is changed to *presence of pathogen-associated host lesions* (PHIPO:0000480), the interaction outcome is changed to *disease present* (PHIPO:0001200), and the extensions for infective ability and control metagenotypes are removed, since they are not applicable.

| Pathogen genotype | Host genotype | Term ID | Term name | Evidence code | Conditions | Figure | Annotation extension |
| --- | --- | --- | --- | --- | --- | --- | --- |
| GT2+[WT level]<br>*Z. tritici*<br>(IPO323) | wild type<br>*T. aestivum*<br>(cv. Riband) | PHIPO:0000480 | presence of pathogen-associated host lesions | Macroscopic observation (qualitative observation) | 14 days post inoculation | Figure 2E | **infects_tissue** leaf , **interaction_outcome** disease present |

The interaction between *Z. tritici* and *T. aestivum* can also be used as a template for the interaction between *F. graminearum* and *T. aestivum*. Here, the pathogen genotype is changed to the GT2 deletion *F. graminearum*, the host strain is changed to cv. Bobwhite, the experimental condition is changed to ‘13 days post inoculation’, the host tissue infected is changed to *inflorescence* (BTO:0000628), the control metagenotype is updated accordingly, and the figure number is changed to 4E.

| Pathogen genotype | Host genotype | Term ID | Term name | Evidence code | Conditions | Figure | Annotation extension |
| --- | --- | --- | --- | --- | --- | --- | --- |
| GT2Δ<br>*F. graminearum*<br>(PH-1) | wild type<br>*T. aestivum*<br>(cv. Bobwhite) | PHIPO:0000481 | absence of pathogen-associated host lesions | Macroscopic observation (qualitative observation) | 13 days post inoculation | Figure 4E | **infects_tissue** inflorescence , **infective_ability** loss of pathogenicity , **compared_to_control** GT2+[WT level] *Fusarium graminearum* (PH-1) / wild type *Triticum aestivum* (cv. Bobwhite) , **interaction_outcome** disease absent |

The changes required for the wild type interaction between *F. graminearum* and *T. aestivum* are the same as those required for *Z. tritici* and *T. aestivum*, since the interaction outcome is the same (presence of pathogen-associated host lesions, and presence of disease).

| Pathogen genotype | Host genotype | Term ID | Term name | Evidence code | Conditions | Figure | Annotation extension |
| --- | --- | --- | --- | --- | --- | --- | --- |
| GT2+[WT level]<br>*F. graminearum*<br>(PH-1) | wild type<br>*T. aestivum*<br>(cv. Bobwhite) | PHIPO:0000480 | presence of pathogen-associated host lesions | Macroscopic observation (qualitative observation) | 13 days post inoculation | Figure 4E | **infects_tissue** leaf , **interaction_outcome** disease present |

Shown below is a table of all the pathogen–host interaction phenotypes from this curation example.

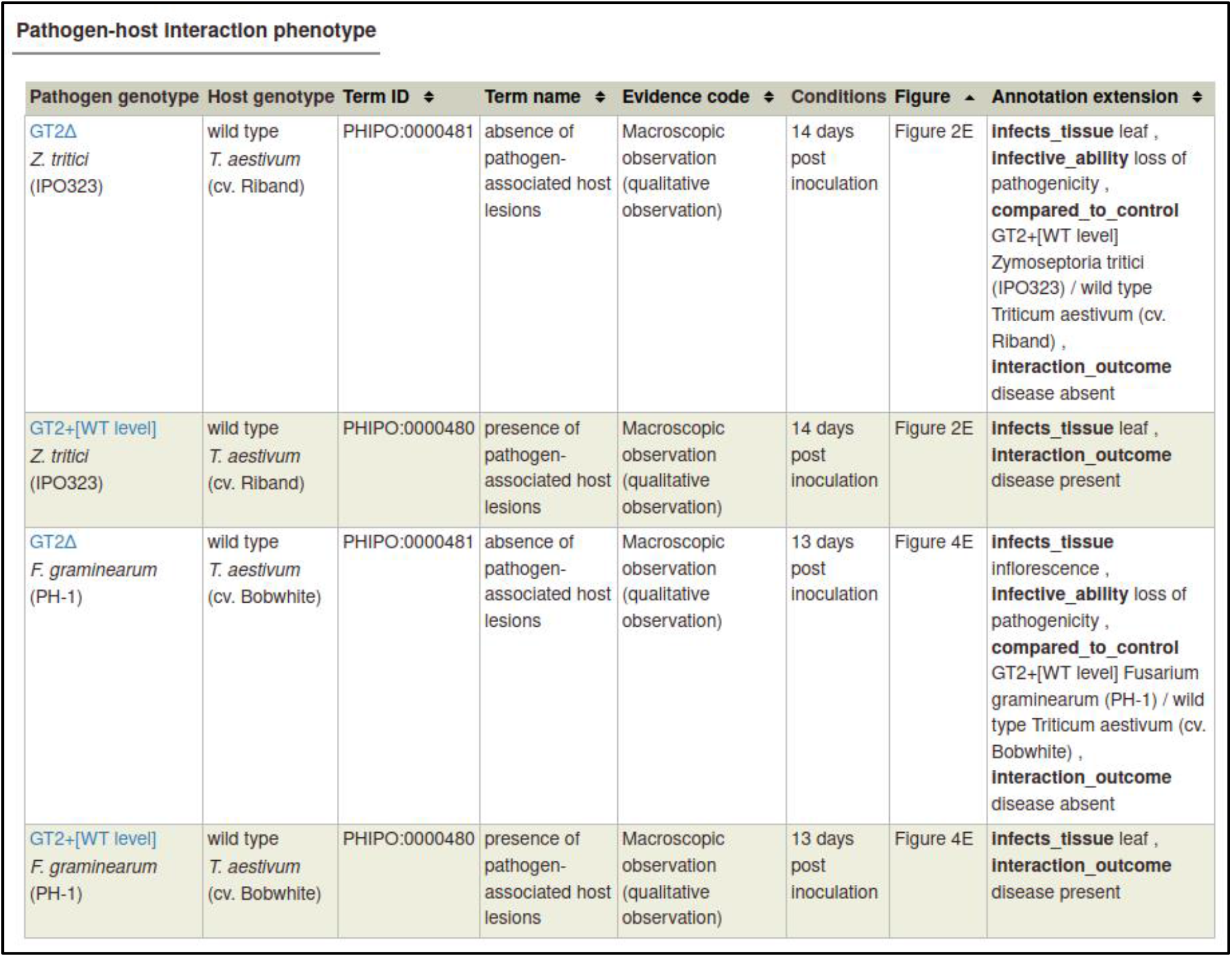

| Pathogen-host Interaction phenotype |  |  |  |  |  |  |  |
| --- | --- | --- | --- | --- | --- | --- | --- |
| Pathogen genotype | Host genotype | Term ID | Term name | Evidence code | Conditions | Figure | Annotation extension |
| GT2Δ<br>*Z. tritici*<br>(IPO323) | wild type<br>*T. aestivum*<br>(cv. Riband) | PHIPO:0000481 | absence of pathogen-associated host lesions | Macroscopic observation (qualitative observation) | 14 days post inoculation | Figure 2E | **infects_tissue** leaf , **infective_ability** loss of pathogenicity , **compared_to_control** GT2+[WT level] *Zymoseptoria tritici* (IPO323) / wild type *Triticum aestivum* (cv. Riband) , **interaction_outcome** disease absent |
| GT2+[WT level]<br>*Z. tritici*<br>(IPO323) | wild type<br>*T. aestivum*<br>(cv. Riband) | PHIPO:0000480 | presence of pathogen-associated host lesions | Macroscopic observation (qualitative observation) | 14 days post inoculation | Figure 2E | **infects_tissue** leaf , **interaction_outcome** disease present |
| GT2Δ<br>*F. graminearum*<br>(PH-1) | wild type<br>*T. aestivum*<br>(cv. Bobwhite) | PHIPO:0000481 | absence of pathogen-associated host lesions | Macroscopic observation (qualitative observation) | 13 days post inoculation | Figure 4E | **infects_tissue** inflorescence , **infective_ability** loss of pathogenicity , **compared_to_control** GT2+[WT level] *Fusarium graminearum* (PH-1) / wild type *Triticum aestivum* (cv. Bobwhite) , **interaction_outcome** disease absent |
| GT2+[WT level]<br>*F. graminearum*<br>(PH-1) | wild type<br>*T. aestivum*<br>(cv. Bobwhite) | PHIPO:0000480 | presence of pathogen-associated host lesions | Macroscopic observation (qualitative observation) | 13 days post inoculation | Figure 4E | **infects_tissue** leaf , **interaction_outcome** disease present |

### Disease annotation

PHI-Canto provides the ‘Disease name’ annotation type, which is used to annotate a disease to a pathogen–host interaction. These annotations highlight the fact that two different pathogens infecting different tissue types of the same host have been used in experiments within this publication.

Disease name annotations are made on the Metagenotype Management page, via the ‘Annotate disease name’ link.

| Pathogen |  | Host |  | Annotations |  |
| --- | --- | --- | --- | --- | --- |
| Species (strain) | Genotype | Species (strain) | Genotype |  |  |
| Z. tritici<br>(IPO323) | GT2+[WT level] | T. aestivum<br>(cv. Riband) | wild type | 1 | Annotate pathogen-host interaction phenotype<br>Annotate gene-for-gene phenotype<br>Annotate disease name<br>View phenotype annotations<br>Delete |

The curator can select a disease from a list of disease names provided by the PHI-base Disease List (PHIDO). For *Z. tritici*, the disease is *septoria leaf blotch* (PHIDO:0000329).

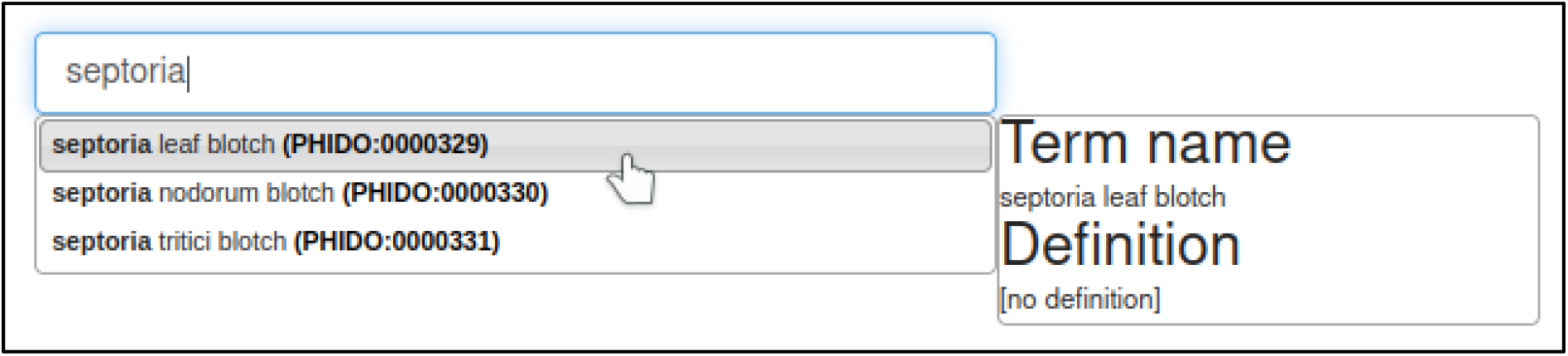

Disease name annotations also allow the host tissue infected to be specified. In this case, the tissue is the *leaf* (BTO:0000713).

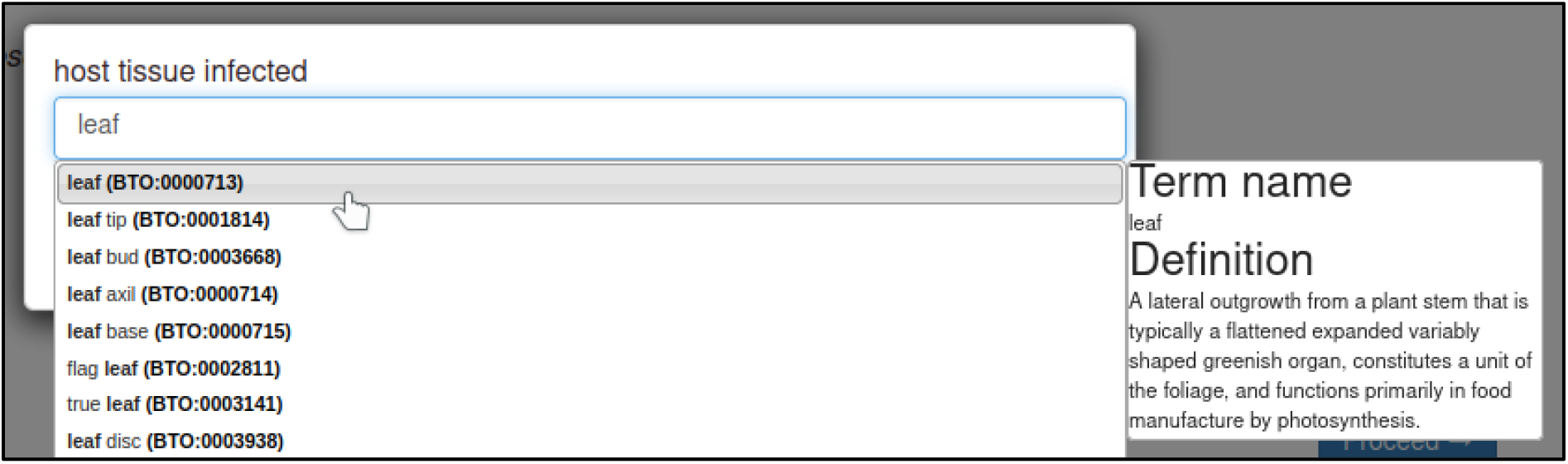

The curator has the option to provide the figure number and additional comments. In this case, the figure numbers are 1 and 2.

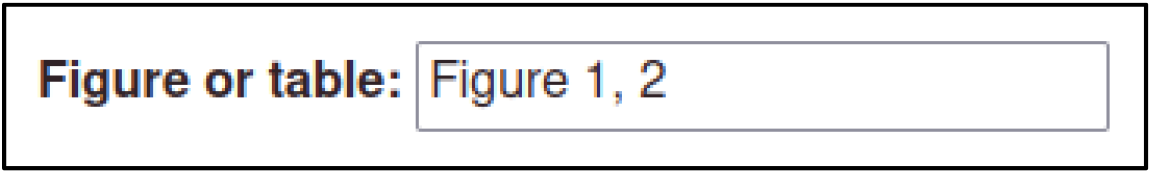

Once this step is completed, the disease name annotation is created.

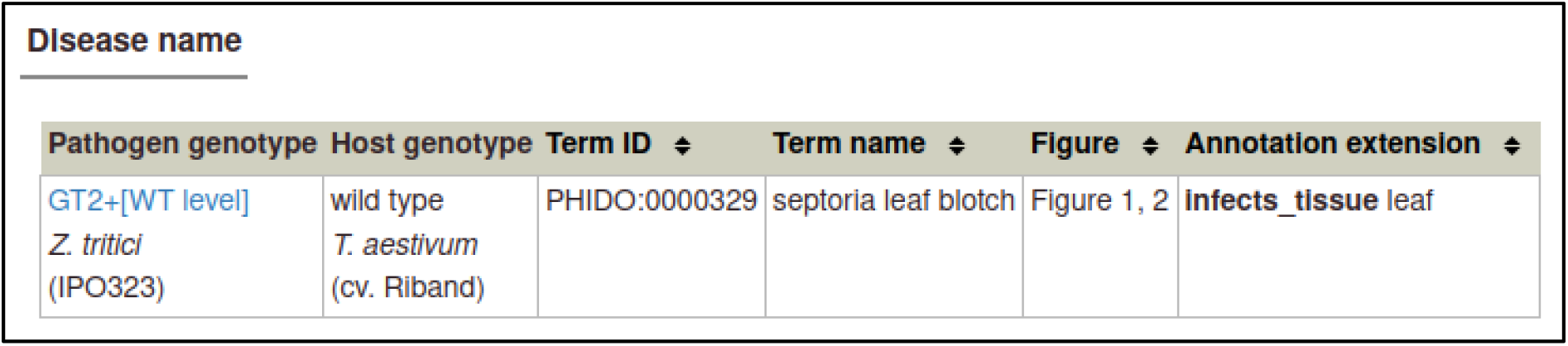

The same process can be followed to create the Disease name annotation for *F. graminearum*: the genotype is the wild type GT2, the host cultivar is *cv. Bobwhite*, the disease is *fusarium ear blight* (PHIDO:0000162), the host tissue infected is the *inflorescence* (BTO:0000628), and the figure number is 4.

| Disease name |  |  |  |  |  |
| --- | --- | --- | --- | --- | --- |
| Pathogen genotype | Host genotype | Term ID | Term name | Figure | Annotation extension |
| GT2+[WT level]<br>*F. graminearum*<br>(PH-1) | wild type<br>*T. aestivum*<br>(cv. Bobwhite) | PHIDO:0000162 | fusarium ear blight | Figure 4 | **infects_tissue** inflorescence |

### Gene Ontology annotation

PHI-Canto also provides the ability to annotate biological processes, molecular functions, and cellular components associated with wild type versions of genes, using terms from the Gene Ontology (GO). In this publication, GT2 is described as having glycotransferase activity as its molecular function, so the curator can annotate this.

Gene Ontology annotations are made by selecting the gene from the Curation Summary page.

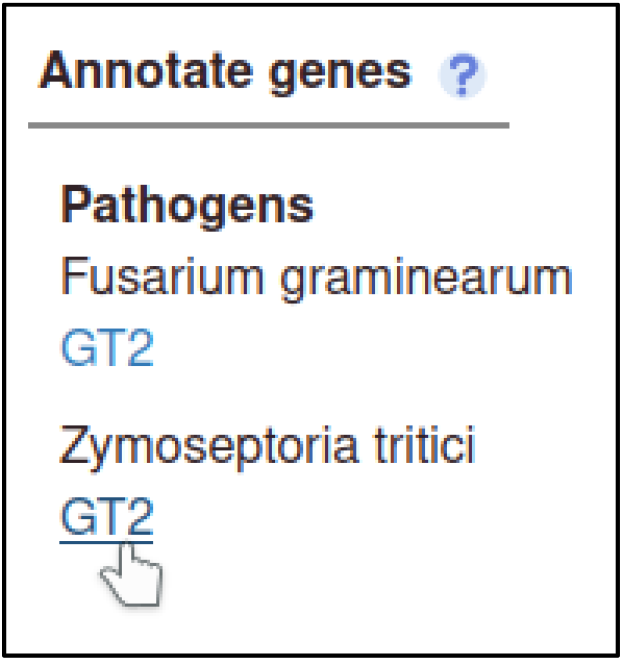

The gene details page has a list of available annotation types.

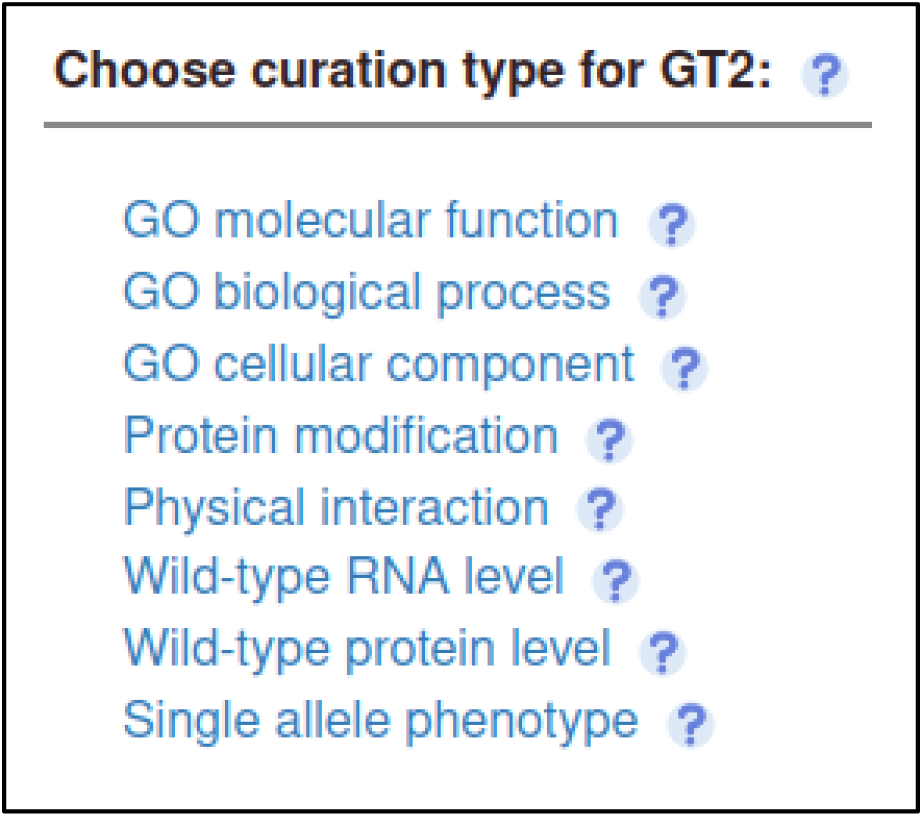

The curator selects the GO Molecular Function annotation type and is prompted for a term from the Gene Ontology. In this case, the correct term is *glycotransferase activity* (GO:0016757).

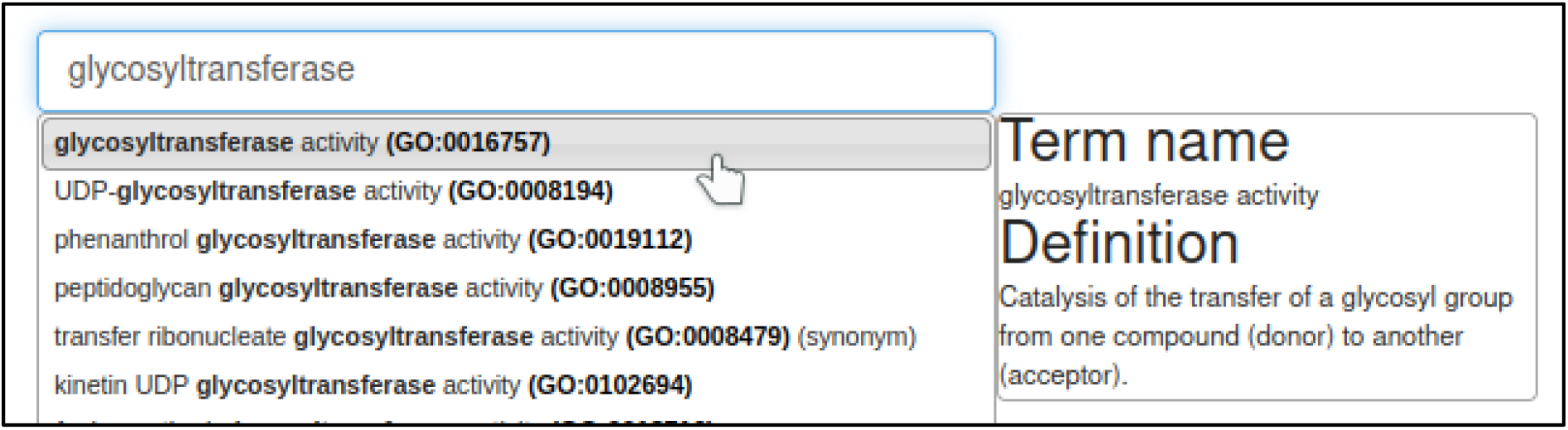

The curator must provide an evidence code from a controlled list specified by the Gene Ontology. The appropriate evidence code in this case is a *Traceable Author Statement* in the publication.

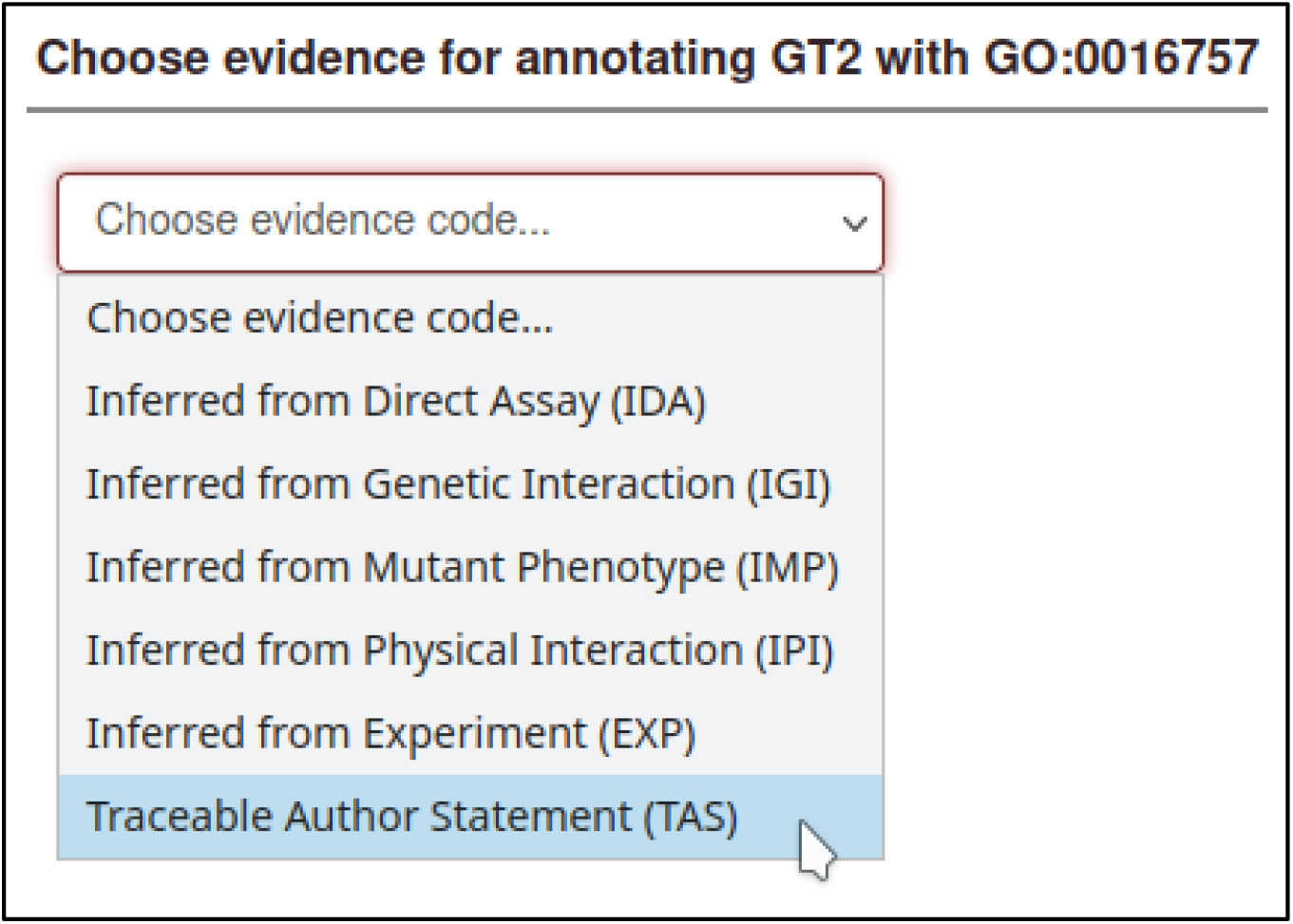

There are many annotation extensions available for GO annotations, but in this case, none of them are applicable (or required), so the curator skips this step.

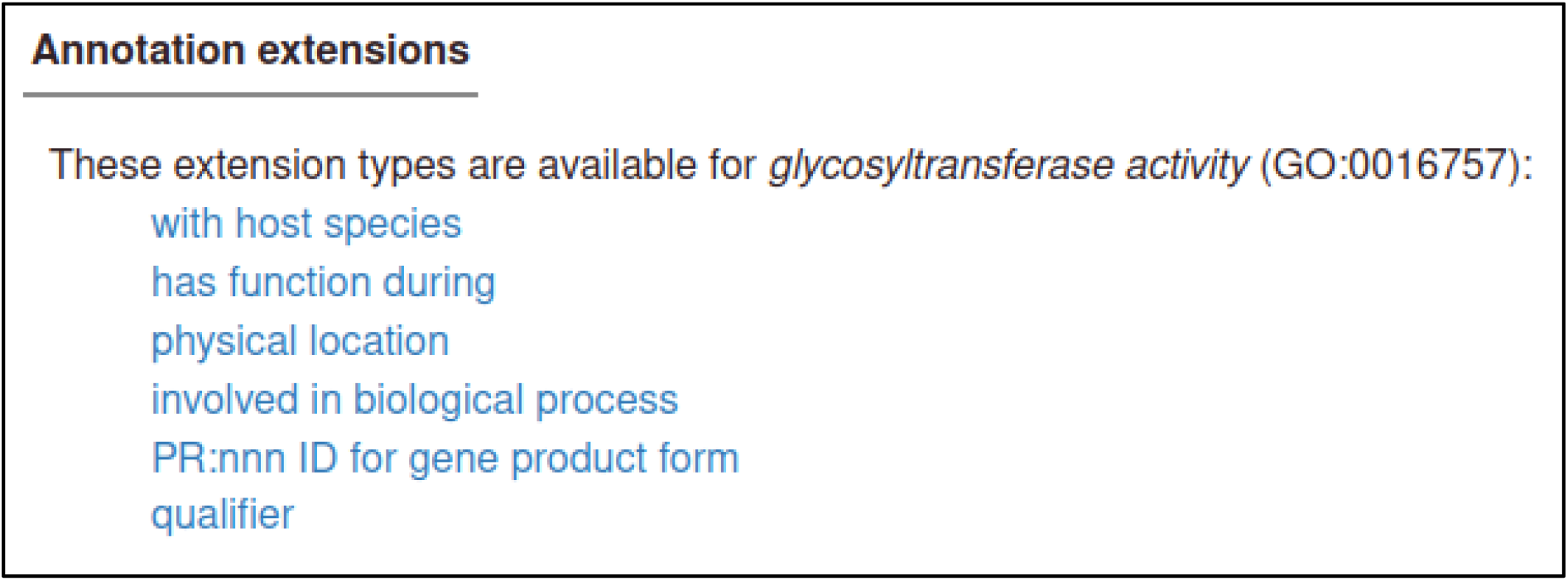

Figure numbers can be specified for GO annotations: in this case, the relevant figure is Figure 3.

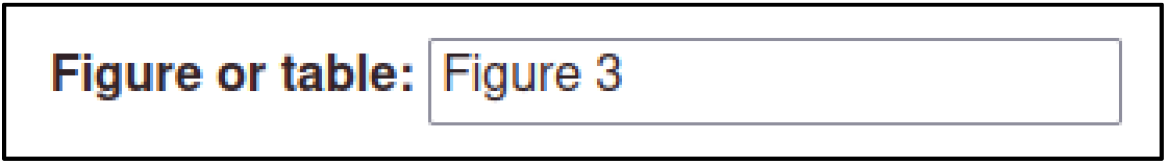

Once this step is completed, the molecular function annotation is created.

GO molecular function
| Species ⇅ | Gene ⇅ | Term ID ⇅ | Term name ⇅ | Evidence code ⇅ | Figure ⇅ |
| --- | --- | --- | --- | --- | --- |
| *Z. tritici* | GT2 | GO:0016757 | glycosyltransferase activity | TAS | Figure 3 |

#### Other annotation types

The publication contains other information which is not included in this worked example for the sake of brevity. In the real curation session, this other information is captured as the following annotations:

- **GO biological process** annotations indicate that GT2 is involved in the hyphal growth process.
- **GO cellular component** annotations indicate that GT2 is located in the hyphal cell wall.
- **Pathogen phenotype** annotations capture information about the pathogen *in vitro*, specifically normal and altered phenotypes for unicellular population growth, hyphal growth, cellular melanin accumulation, filament morphology, and so on.

All these annotation types use the same annotation process as the annotation types described above.

### Submitting the curation session

Once the curator has made all their annotations, the curation session is submitted to the PHI-base team for review.

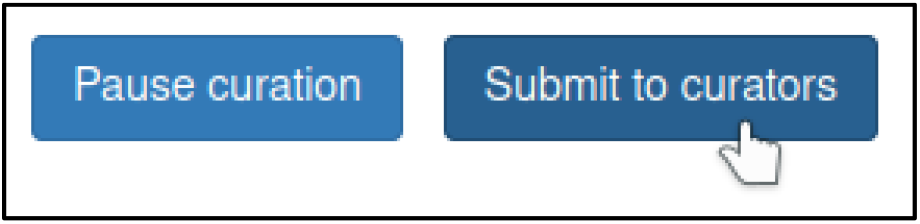

The curator can use a text box to provide any information that is outside the scope of the curation process before finishing the submission process. Once the submission process is finished, the curation session can no longer be edited except by members of the PHI-base team, who have the option to reactivate the session in case changes are required by the original curator.

## Appendix 3 Author checklist prior to publication

1. Use the most current gene name. Take care with synonyms. Prefix the gene name with the genus and species initials if the same gene name exists in multiple species.
2. If reporting on a new (gene) sequence, submit your sequence to NCBI GenBank or the European Nucleotide Archive (ENA), then obtain an accession number prior to publication. Record this accession number within the manuscript. If reporting on a gene with an existing accession number, make sure this is reported in the manuscript. Please record the UniProtKB accession number for the protein of the gene, where available. Provide or use any existing informative allele or line designations for mutations and transgenes.
3. Provide a binomial species name for pathogen and host organisms, not just a common name. If possible, please also include NCBI Taxonomy IDs for the pathogen and host organisms at the rank of species.
4. Describe the tissue or organ in which the experimental observations were made (controlled language can be found in the BRENDA Tissue Ontology, see https://www.ebi.ac.uk/ols/ontologies/bto).
5. Describe any experimental techniques used, and accurately record any chemicals or reagents used.
6. When writing an article, try to keep the use of descriptive language as accurate and controlled as possible. For example, do not use ‘reduced pathogenicity’ or ‘loss of virulence’, as these terms can be misleading: it would be more accurate to use ‘reduced virulence’ and ‘loss of pathogenicity’, respectively. Ideally, try to follow the terminology of an existing ontology: this will make the data easier to extract and reuse. Relevant ontologies include PHIPO and GO (https://www.ebi.ac.uk/ols/ontologies/phipo, https://www.ebi.ac.uk/ols/ontologies/go).
7. Document all the key information for the paper: do not rely on citing past papers for information on the pathogen used, or the strain used, and so on.
8. Think carefully when choosing keywords for your manuscript to ensure that the publication can be located by PHI-base’s keyword searches. One example of an ideal keyword is ‘pathogen-host interaction’.
9. Record the provenance of the pathogen strain: for example, whether it is a lab strain or a field isolate, or if the strain was obtained from a stock center or as a gift from another lab.

---

**Figure 3 – figure supplement 1.**
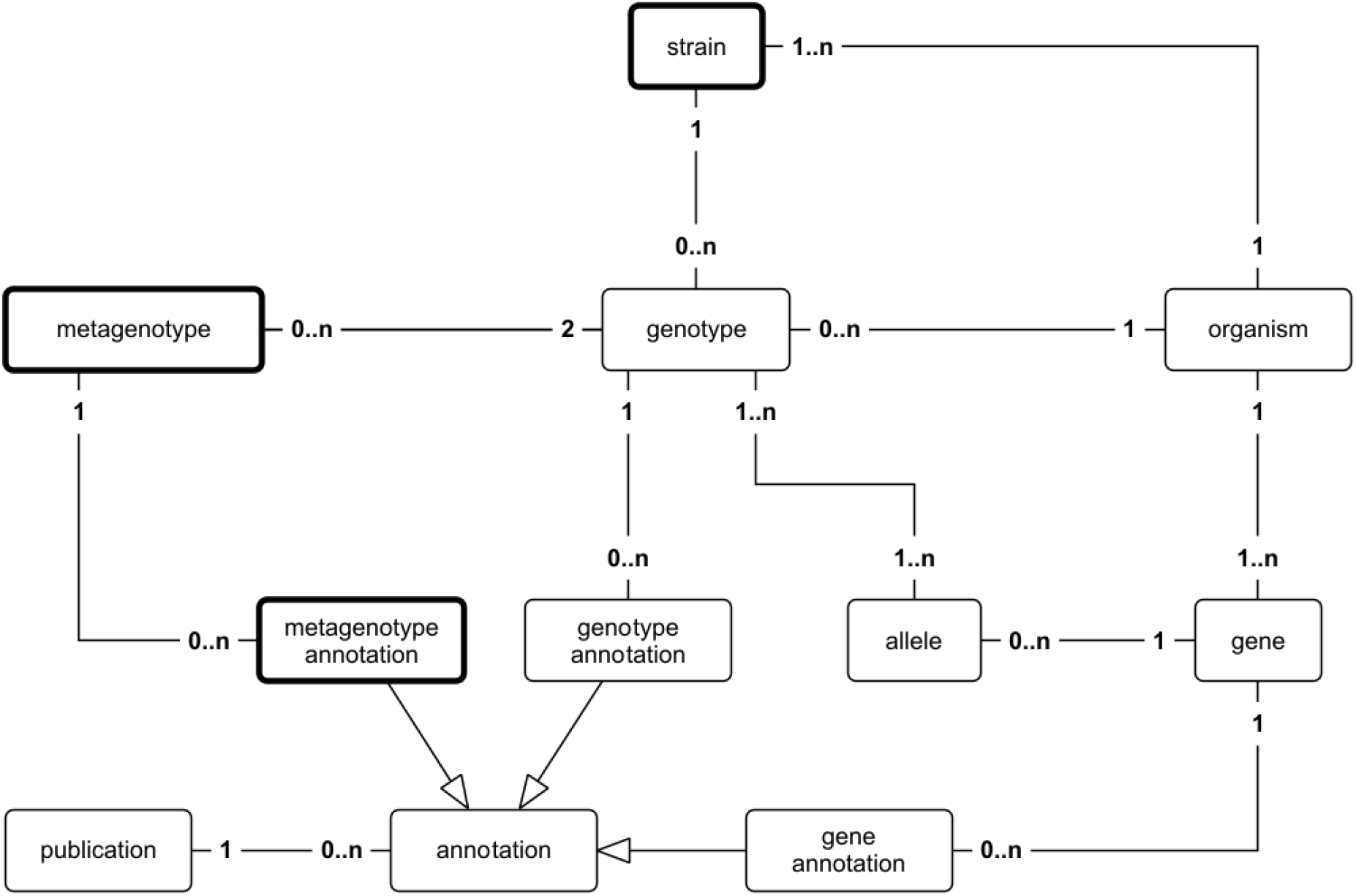
Canto entity relationship diagram. Simplified UML class diagram showing the relations between entities (things of interest) in a Canto curation session. The numbers on the connecting lines represent the cardinality of the relation, meaning how many of one entity can be related to another entity: 0..n means ‘zero or more’; 1..n means ‘one or more’. Lines with a hollow arrowhead indicate that the target entity (at the head of the arrow) is a generalization of the source entity (at the tail of the arrow). Boxes outlined in bold indicate new entities which were added to support curation in PHI-Canto.

**Figure 4 – figure supplement 1.**
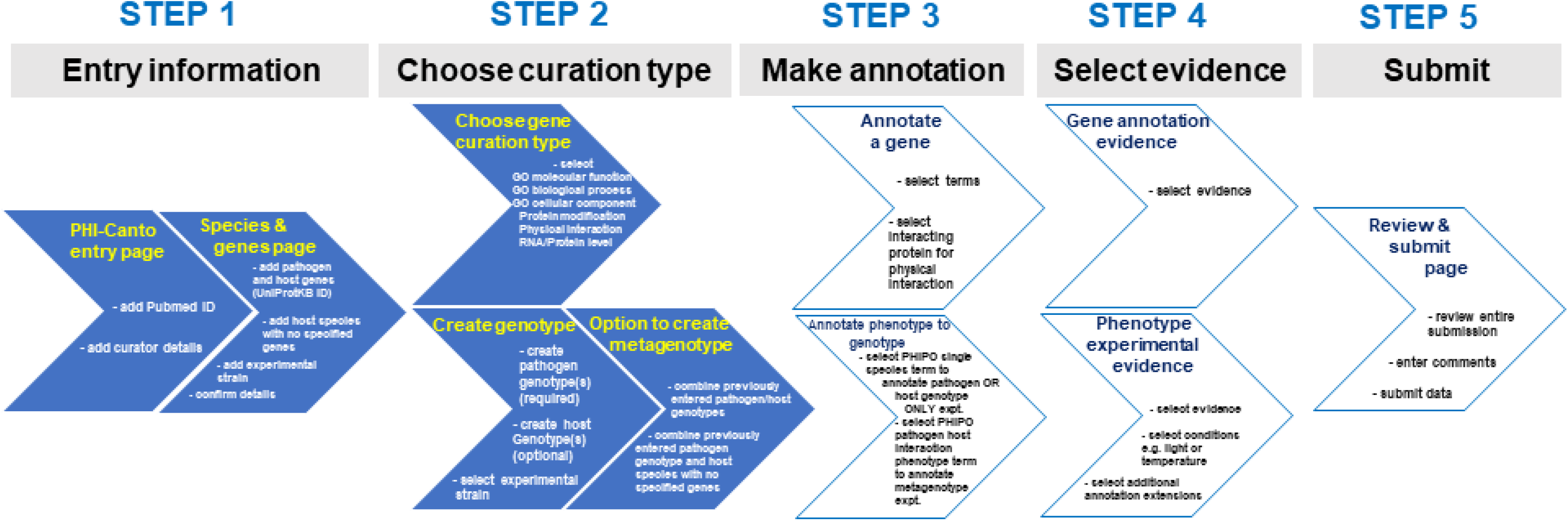
Alternative curation step workflow. The flow diagram represents the PHI-Canto curation process from beginning to end in 5 steps. It is an alternative representation to the image depicted in Figure 4. During step 2 of the workflow, the curator chooses either the gene annotation or genotype / metagenotype annotation process. Multiple annotations can be made using both annotation processes which can then be submitted for review.

**Figure 4 - figure supplement 2.** What you need to curate a publication into PHI-Canto.

1. The PubMed ID of the peer-reviewed publication.
2. Your email address, so we can contact you regarding your curation session.
3. UniProtKB accession numbers for the pathogen and host gene products studied within the publication.
4. The binomial names of the pathogen and host species studied within the publication.
5. Details of the experimental strains used within the publication.

**Figure 4 - figure supplement 3.** Instructions on how to look up a UniProtKB ID.

UniProtKB is divided into two sources: UniProtKB/Swiss-Prot, which contains manually annotated entries; and UniProtKB/TrEMBL, which contains unreviewed entries that are automatically annotated by prediction systems. PHI-Canto permits annotations on entries from either source, although UniProtKB/Swiss-Prot entries are preferred (owing to their higher quality).

## Finding genes in UniProtKB

PHI-Canto uses <u>UniProt Knowledgebase</u> (UniProtKB) gene accession numbers to disambiguate genes/proteins. This is to ensure that we are talking about the correct gene product – especially as the same names are sometimes used for different proteins – and to standardize entries, because not all strains are in UniProt.

**1. Identify the reference proteome** (we use the designated reference proteome to integrate different strain information at the gene level in PHI-base). In PHI-Canto you will be able to specify the strain you used.

Look up the reference proteome for your organism using the species name (https://www.uniprot.org/help/reference_proteome).

If there is no reference proteome use the strain studied.

**2. Identify the gene of interest** in the reference proteome:

Start from the <u>UniProt homepage</u>, then perform any of the following steps:

Search for the author assigned gene name/primary name (e.g., Tri5) or synonyms, plus species name (e.g., *Fusarium graminearum*).

OR

If the gene does not have a ‘given name’ but a locus ID is provided, search using the locus_id (e.g., FGRRES_03537) plus species name (e.g., *Fusarium graminearum*). If the entry identifier used is not the reference strain, copy the protein sequence and go to the BLAST step below.

OR

Search on a protein description (e.g., **Trichodiene synthase**)

OR

Obtain the protein sequence for your gene of interest and BLAST against UniprotKB (https://www.uniprot.org/blast/) with your protein sequence.

**Note:** If there are multiple entries for your gene product from the reference strain, please select the ‘Reviewed entry’. Use the left-hand filter for ‘Reviewed entries’.

OR

If the gene cannot be located in UniProt, contact the authors, UniProt, or PHI-base for help locating the canonical database entry.

**3. Add the entry into PHI-Canto.** Once the entry of interest is located, select the entry accession number (also called ‘Entry’) from column 1 of the results table, and use this to retrieve the entry into PHI-Canto on the gene entry page. **Caution:** Do not confuse the ‘Entry’ column with the ‘Entry name’ column. PHI-Canto uses the accession number to retrieve details (such as the gene name, gene product, and organism). If PHI-Canto is unable to find your entry, check for typos (e.g., 0 for O), ensure you are using the ‘entry’ not ‘entry name’, and check that your accession is from UniProtKB, not UniParc.

**Figure 5 – figure supplement 1.**
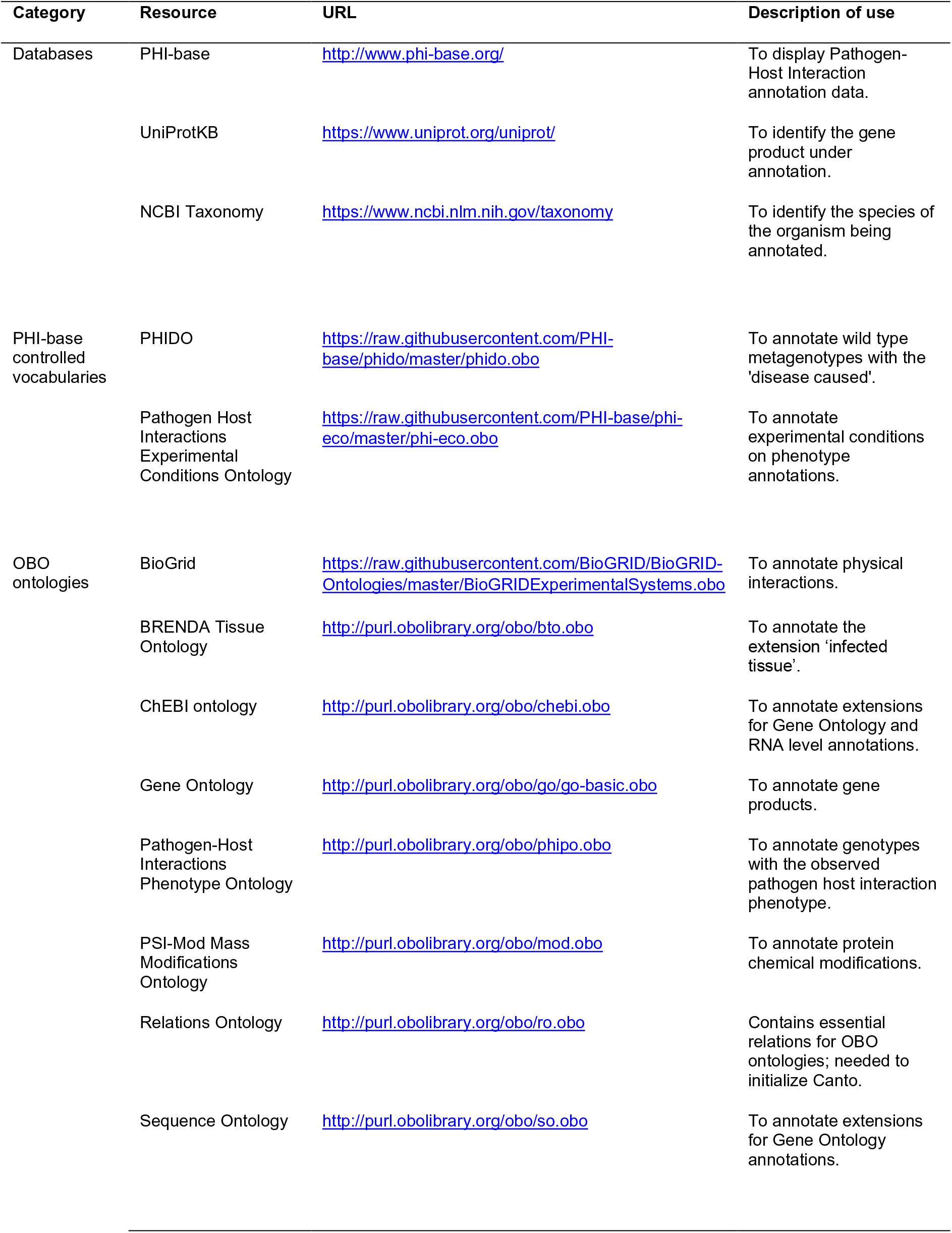
Resources relied upon by PHI-Canto.

## References

Alliance of Genome Resources Consortium. (2020). Alliance of Genome Resources Portal: unified model organism research platform. Nucleic Acids Res, 48(D1), D650–D658. doi:10.1093/nar/gkz813

Arita, M., Karsch-Mizrachi, I., & Cochrane, G. (2021). The international nucleotide sequence database collaboration. Nucleic Acids Res, 49(D1), D121–D124. doi:10.1093/nar/gkaa967

Ashburner, M., Ball, C. A., Blake, J. A., Botstein, D., Butler, H., Cherry, J. M., … Sherlock, G. (2000). Gene ontology: tool for the unification of biology. Nat Genet, 25(1), 25–29. doi:10.1038/75556

Bebber, D. P., Ramotowski, M. A. T., & Gurr, S. J. (2013). Crop pests and pathogens move polewards in a warming world. Nature Climate Change, 3(11), 985–988. doi:10.1038/Nclimate1990

Breen, S., Williams, S. J., Winterberg, B., Kobe, B., & Solomon, P. S. (2016). Wheat PR-1 proteins are targeted by necrotrophic pathogen effector proteins. Plant J, 88(1), 13–25. doi:10.1111/tpj.13228

Brown, G. D., Denning, D. W., Gow, N. A., Levitz, S. M., Netea, M. G., & White, T. C. (2012). Hidden killers: human fungal infections. Sci Transl Med, 4(165), 165rv113. doi:10.1126/scitranslmed.3004404

Chaloner, T. M., Gurr, S. J., & Bebber, D. P. (2021). Plant pathogen infection risk tracks global crop yields under climate change. Nature Climate Change, 11(8), 710-+. doi:10.1038/s41558-021-01104-8

Cook, N. M., Chng, S., Woodman, T. L., Warren, R., Oliver, R. P., & Saunders, D. G. (2021). High frequency of fungicide resistance-associated mutations in the wheat yellow rust pathogen *Puccinia striiformis f. sp. tritici*. Pest Manag Sci, 77(7), 3358–3371. doi:10.1002/ps.6380

Federhen, S., Clark, K., Barrett, T., Parkinson, H., Ostell, J., Kodama, Y., … Karsch-Mizrachi, I. (2014). Toward richer metadata for microbial sequences: replacing strain-level NCBI taxonomy taxids with BioProject, BioSample and Assembly records. Stand Genomic Sci, 9(3), 1275–1277. doi:10.4056/sigs.4851102

Fisher, M. C., Hawkins, N. J., Sanglard, D., & Gurr, S. J. (2018). Worldwide emergence of resistance to antifungal drugs challenges human health and food security. Science, 360(6390), 739–742. doi:10.1126/science.aap7999

Fisher, M. C., Henk, D. A., Briggs, C. J., Brownstein, J. S., Madoff, L. C., McCraw, S. L., & Gurr, S. J. (2012). Emerging fungal threats to animal, plant and ecosystem health. Nature, 484(7393), 186–194. doi:10.1038/nature10947

Flor, H. H. (1956). The complementary genic systems in Flax and Flax Rust. In M. Demerec (Ed.), Advances in Genetics (Vol. 8, pp. 29–54): Academic Press.

Gene Ontology Consortium. (2021). The Gene Ontology resource: enriching a GOld mine. Nucleic Acids Res, 49(D1), D325–D334. doi:10.1093/nar/gkaa1113

Giglio, M., Tauber, R., Nadendla, S., Munro, J., Olley, D., Ball, S., …Chibucos, M. C. (2019). ECO, the Evidence & Conclusion Ontology: community standard for evidence information. Nucleic Acids Res, 47(D1), D1186–D1194. doi:10.1093/nar/gky1036

Houterman, P. M., Ma, L., van Ooijen, G., de Vroomen, M. J., Cornelissen, B. J., Takken, F. L., & Rep, M. (2009). The effector protein Avr2 of the xylem-colonizing fungus Fusarium oxysporum activates the tomato resistance protein I-2 intracellularly. Plant J, 58(6), 970–978. doi:10.1111/j.1365-313X.2009.03838.x

Huntley, R. P., Harris, M. A., Alam-Faruque, Y., Blake, J. A., Carbon, S., Dietze, H., … Mungall, C. J. (2014). A method for increasing expressivity of Gene Ontology annotations using a compositional approach. BMC Bioinformatics, 15, 155. doi:10.1186/1471-2105-15-155

Huntley, R. P., Sawford, T., Mutowo-Meullenet, P., Shypitsyna, A., Bonilla, C., Martin, M. J., & O’Donovan, C. (2015). The GOA database: gene Ontology annotation updates for 2015. Nucleic Acids Res, 43(Database issue), D1057–1063. doi:10.1093/nar/gku1113

International Society for Biocuration. (2018). Biocuration: distilling data into knowledge. PLoS Biol, 16(4), e2002846. doi:10.1371/journal.pbio.2002846

Jackson, R., Matentzoglu, N., Overton, J. A., Vita, R., Balhoff, J. P., Buttigieg, P. L., … Peters, B. (2021). OBO Foundry in 2021: operationalizing open data principles to evaluate ontologies. Database (Oxford), 2021. doi:10.1093/database/baab069

Jackson, R. C., Balhoff, J. P., Douglass, E., Harris, N. L., Mungall, C. J., & Overton, J. A. (2019). ROBOT: a tool for automating ontology workflows. BMC Bioinformatics, 20(1), 407. doi:10.1186/s12859-019-3002-3

Jones, J. D., & Dangl, J. L. (2006). The plant immune system. Nature, 444(7117), 323–329. doi:10.1038/nature05286

Kanyuka, K., Igna, A. A., Solomon, P. S., & Oliver, R. P. (2022). The rise of necrotrophic effectors. New Phytol, 233(1), 11–14. doi:10.1111/nph.17811

Montecchi-Palazzi, L., Beavis, R., Binz, P. A., Chalkley, R. J., Cottrell, J., Creasy, D., … Garavelli, J. S. (2008). The PSI-MOD community standard for representation of protein modification data. Nat Biotechnol, 26(8), 864–866. doi:10.1038/nbt0808-864

Musen, M. A., & Protege Team. (2015). The Protege project: A look back and a look forward. AI Matters, 1(4), 4–12. doi:10.1145/2757001.2757003

Oughtred, R., Rust, J., Chang, C., Breitkreutz, B. J., Stark, C., Willems, A., … Tyers, M. (2021). The BioGRID database: A comprehensive biomedical resource of curated protein, genetic, and chemical interactions. Protein Sci, 30(1), 187–200. doi:10.1002/pro.3978

Rodriguez-Iglesias, A., Rodriguez-Gonzalez, A., Irvine, A. G., Sesma, A., Urban, M., Hammond-Kosack, K. E., & Wilkinson, M. D. (2016). Publishing FAIR data: an exemplar methodology utilizing PHI-base. Front Plant Sci, 7, 641. doi:10.3389/fpls.2016.00641

Rutherford, K. M., Harris, M. A., Lock, A., Oliver, S. G., & Wood, V. (2014). Canto: an online tool for community literature curation. Bioinformatics, 30(12), 1791–1792. doi:10.1093/bioinformatics/btu103

Schoch, C. L., Ciufo, S., Domrachev, M., Hotton, C. L., Kannan, S., Khovanskaya, R., … Karsch-Mizrachi, I. (2020). NCBI Taxonomy: a comprehensive update on curation, resources and tools. Database (Oxford), 2020. doi:10.1093/database/baaa062

Scholthof, K. B. (2007). The disease triangle: pathogens, the environment and society. Nat Rev Microbiol, 5(2), 152–156. doi:10.1038/nrmicro1596

Shefchek, K. A., Harris, N. L., Gargano, M., Matentzoglu, N., Unni, D., Brush, M., … Osumi-Sutherland, D. (2020). The Monarch Initiative in 2019: an integrative data and analytic platform connecting phenotypes to genotypes across species. Nucleic Acids Res, 48(D1), D704–D715. doi:10.1093/nar/gkz997

Smith, K. M., Machalaba, C. C., Seifman, R., Feferholtz, Y., & Karesh, W. B. (2019). Infectious disease and economics: The case for considering multi-sectoral impacts. One Health, 7, 100080. doi:10.1016/j.onehlt.2018.100080

UniProt Consortium. (2021). UniProt: the universal protein knowledgebase in 2021. Nucleic Acids Res, 49(D1), D480–D489. doi:10.1093/nar/gkaa1100

Urban, M., Cuzick, A., Rutherford, K., Irvine, A., Pedro, H., Pant, R., … Hammond-Kosack, K. E. (2017). PHI-base: a new interface and further additions for the multi-species pathogen-host interactions database. Nucleic Acids Res, 45(D1), D604–D610. doi:10.1093/nar/gkw1089

Urban, M., Cuzick, A., Seager, J., Wood, V., Rutherford, K., Venkatesh, S. Y., … Hammond-Kosack, K. E. (2020). PHI-base: the pathogen-host interactions database. Nucleic Acids Res, 48(D1), D613–D620. doi:10.1093/nar/gkz904

Urban, M., Cuzick, A., Seager, J., Wood, V., Rutherford, K., Venkatesh, S. Y., … Hammond-Kosack, K. E. (2021). PHI-base in 2022: a multi-species phenotype database for Pathogen-Host Interactions. Nucleic Acids Res. doi:10.1093/nar/gkab1037

Urban, M., Pant, R., Raghunath, A., Irvine, A. G., Pedro, H., & Hammond-Kosack, K. E. (2015). The Pathogen-Host Interactions database (PHI-base): additions and future developments. Nucleic Acids Res, 43(Database issue), D645–655. doi:10.1093/nar/gku1165

Wilkinson, M. D., Dumontier, M., Aalbersberg, I. J., Appleton, G., Axton, M., Baak, A., … Mons, B. (2016). The FAIR Guiding Principles for scientific data management and stewardship. Sci Data, 3, 160018. doi:10.1038/sdata.2016.18

Winnenburg, R., Baldwin, T. K., Urban, M., Rawlings, C., Kohler, J., & Hammond-Kosack, K. E. (2006). PHI-base: a new database for pathogen host interactions. Nucleic Acids Res, 34(Database issue), D459–464. doi:10.1093/nar/gkj047

Wood, V., Sternberg, P. W., & Lipshitz, H. D. (2022). Making biological knowledge useful for humans and machines. Genetics, 220(4). doi:10.1093/genetics/iyac001

Yates, A. D., Allen, J., Amode, R. M., Azov, A. G., Barba, M., Becerra, A., … Flicek, P. (2021). Ensembl Genomes 2022: an expanding genome resource for non-vertebrates. Nucleic Acids Res. doi:10.1093/nar/gkab1007

